# Physiological Aging and Cardiovascular Disease Distinctly Remodel the Human Cardiac Lymphatic Endothelium

**DOI:** 10.64898/2026.09.22.750171

**Authors:** Haotian Chen, Sydney Chao, Yanxi Shi, Zihui Yang, Jason D Roh, Paul B Yu, Peng Xia

## Abstract

**Aims:** Cardiac lymphatic endothelial cells (LECs) regulate tissue homeostasis, immune trafficking, and remodeling, but how human cardiac LEC heterogeneity changes with physiological aging and cardiovascular disease remains poorly defined. We sought to define age- and disease-associated LEC states, their regulatory and microenvironmental programs, and the functional relevance of selected pathways.

**Methods and results:** We integrated human cardiac single-nucleus transcriptomic datasets spanning physiological aging, dilated and hypertrophic cardiomyopathy, doxorubicin-associated cardiomyopathy, and heart failure with preserved ejection fraction. Aging redistributed LECs among pre-existing states, with expansion of a homeostatic immune-surveillance state and loss of cytokine-responsive and polarity-associated states. Cardiovascular diseases showed a distinct trajectory, with depletion of a non-failing homeostatic state and expansion of shared and disease-associated remodeling states. SCENIC and NicheNet/MultiNicheNet identified state-specific regulatory programs and disease-dependent signaling networks, with recurrent BMP/TGF-β/activin-family signaling. ACVR2A was prioritized in human doxorubicin-associated cardiomyopathy; endothelial Acvr2a deletion in aged doxorubicin-treated mice was associated with improved cardiac function and increased open cardiac lymphatic vessels. Independent human spatial transcriptomic analyses supported selected aging- and disease-associated programs.

**Conclusion:** Physiological aging and cardiovascular disease remodel cardiac LECs through distinct changes in cellular state, transcriptional regulation, and microenvironmental signaling. Aging is associated with redistribution and reduced representation of specialized LEC programs, whereas cardiovascular diseases redirect LECs through disease-associated signaling environments, highlighting context-dependent remodeling of the cardiac lymphatic endothelium across aging and disease.

**Translational Perspective:** Cardiac lymphatic dysfunction may contribute to impaired immune resolution and myocardial remodeling, but human disease-associated LEC biology remains poorly understood. We identify distinct LEC remodeling trajectories during physiological aging and cardiovascular disease and identify age-associated loss of specialized cytokine-responsive and structural LEC programs. Disease-specific microenvironmental analyses nominate actionable signaling pathways, while endothelial Acvr2a deletion preserves cardiac lymphatic remodeling and improves cardiac function following doxorubicin injury. These findings highlight LEC state remodeling and disease-associated signaling as potential targets for preserving cardiac lymphatic function during aging and cardiovascular disease.

## Introduction

The cardiac lymphatic vasculature is an essential component of cardiovascular homeostasis, regulating interstitial fluid balance, immune-cell trafficking, lipid transport, and tissue remodeling^1–4^. Lymphatic endothelial cells (LECs), which form the lymphatic vascular network, are increasingly recognized as active participants in tissue-specific immune and stromal communication rather than passive conduits for fluid drainage^3, 5–7^. In the heart, disruption of lymphatic integrity or function can promote edema, persistent inflammation, fibrosis, and adverse cardiac remodeling, whereas enhancement of lymphatic growth or function can facilitate resolution after cardiovascular injury^1, 4^. Despite this emerging importance, the cellular and molecular heterogeneity of human cardiac LECs remains incompletely defined, particularly across physiological aging and different forms of cardiovascular disease^4, 8^.

Aging represents a major biological determinant of cardiovascular vulnerability and is accompanied by progressive alterations in endothelial function, immune homeostasis, extracellular matrix organization, and tissue repair^7, 9, 10^. These changes are likely to influence cardiac lymphatic biology, yet how physiological aging remodels human cardiac LEC populations is poorly understood. Importantly, age-related endothelial dysfunction may not simply reflect loss of vascular structure^9^. LECs can adopt distinct transcriptional and functional programs in response to cytokines, extracellular matrix signals, metabolic demands, and signals from neighboring cells. Aging could therefore alter the distribution or responsiveness of specific LEC states, thereby changing how the cardiac lymphatic endothelium adapts to its local and systemic environment. Whether such age-associated remodeling occurs in the human heart, which LEC states are most affected, and whether these changes are associated with altered regulatory and intercellular communication programs remain unresolved.

Cardiovascular disease presents an additional level of complexity. Dilated cardiomyopathy (DCM), hypertrophic cardiomyopathy (HCM), anthracycline-associated cardiomyopathy (DoxCM), and heart failure with preserved ejection fraction (HFpEF) arise from biologically distinct insults and involve different combinations of inflammation, mechanical stress, metabolic dysfunction, extracellular matrix remodeling, and vascular injury^11–15^. The lymphatic endothelium is exposed to these disease-specific microenvironments and may consequently undergo both shared and context-dependent adaptations. However, it remains unclear whether cardiovascular diseases converge on a common pathological LEC phenotype or instead generate distinct LEC states and signaling networks. Resolving this distinction is important for understanding whether cardiac lymphatic dysfunction represents a common downstream feature of cardiovascular disease or a collection of disease-specific endothelial responses.

Recent single-cell and single-nucleus transcriptomic approaches provide an opportunity to address these questions by resolving rare endothelial populations within complex human cardiac tissue^8^. Nevertheless, defining transcriptional clusters alone provides limited insight into their biological significance. A more comprehensive understanding requires integration of LEC state abundance with transcriptional regulatory programs, pathway activity, and predicted communication between LECs and the surrounding cardiac microenvironment^16, 17^. Furthermore, because dissociation-based transcriptomic analyses lose spatial information and computationally inferred signaling does not establish functional relevance, orthogonal tissue-level and experimental approaches are needed to determine whether prioritized LEC programs are supported in intact human myocardium and associated with functional endothelial phenotypes.

Here, we integrated human cardiac single-nucleus RNA-sequencing datasets to define LEC states across physiological aging and multiple cardiovascular diseases using a standardized endothelial-to-LEC analytical framework. We first determined how LEC state composition changes across adult aging and characterized the marker, pathway, and transcriptional regulatory programs associated with age-sensitive states. We further integrated transcriptional regulatory and intercellular communication analyses to characterize intrinsic and microenvironmental programs associated with age-sensitive LEC states. In parallel, we defined LEC remodeling across DCM, HCM, DoxCM, and HFpEF and integrated transcriptional regulatory and intercellular communication analyses to distinguish shared from disease-specific programs and prioritize candidate signaling pathways. Selected aging- and disease-associated programs were further examined in independent spatial transcriptomic datasets, while endothelial *Acvr2a* perturbation *in vivo* provided functional assessment of a prioritized disease-associated signaling pathway.

Together, our findings reveal that physiological aging and cardiovascular disease do not produce a uniform cardiac LEC phenotype. Instead, they remodel the human cardiac lymphatic endothelium through distinct but partially convergent changes in LEC state composition, transcriptional regulation, and extracellular communication. Aging was associated with redistribution of specific LEC states and altered regulatory and microenvironmental signaling programs, whereas cardiovascular diseases exhibited both shared remodeling and disease-specific regulatory and signaling programs. Independent spatial and functional analyses provide orthogonal support for selected findings and identify candidate pathways linking disease-associated endothelial signaling to lymphatic and cardiac remodeling. These findings establish a framework for understanding cardiac LEC heterogeneity across aging and cardiovascular disease and highlight the lymphatic endothelium as a dynamic component of the cardiac microenvironment.

## Results

### Integrated human cardiac datasets define lymphatic endothelial states across physiological aging and cardiovascular disease

To characterize lymphatic endothelial cell (LEC) heterogeneity across human cardiac aging and cardiovascular disease, we integrated cardiac single-nucleus RNA-sequencing datasets spanning physiologically aging non-failing hearts and four cardiovascular disease groups, including dilated cardiomyopathy (DCM), hypertrophic cardiomyopathy (HCM), doxorubicin-associated cardiomyopathy (DoxCM), and heart failure with preserved ejection fraction (HFpEF)^8, 18–22^ (Fig. 1A, S1, and Table S1). Following standardized quality control, endothelial cells were first identified from the complete cardiac cellular populations and subsequently reclustered to resolve cardiac LECs using canonical lymphatic markers, including PROX1, FLT4, LYVE1, PDPN, CCL21, and MMRN1 (Fig. 1B, C)^8, 18^. This two-step strategy preserved rare LEC populations while minimizing contamination from other endothelial populations.

**Figure 1.**
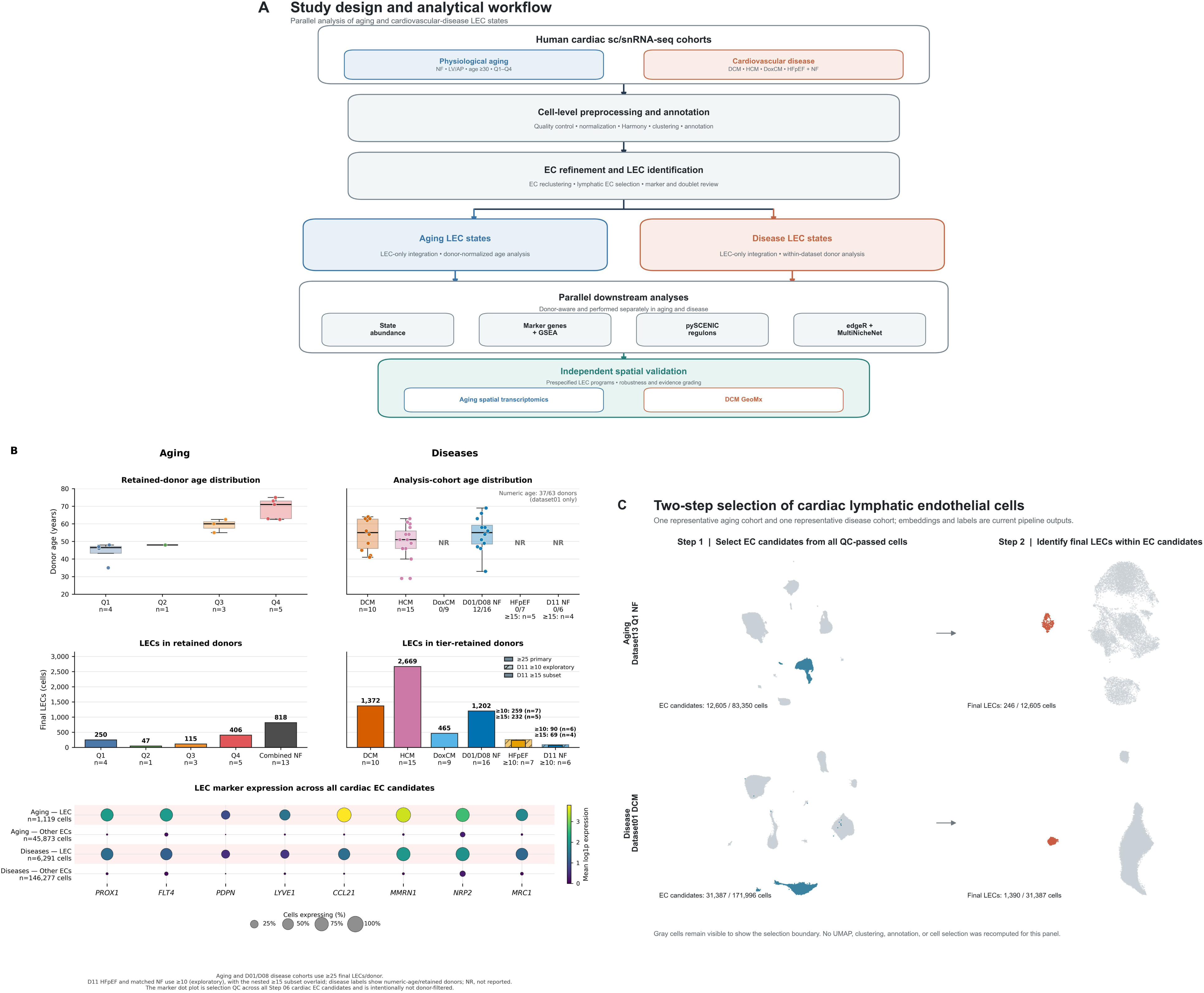
Integrated human cardiac transcriptomic framework for defining lymphatic endothelial cell states across physiological aging and cardiovascular disease. (A) Study design and analytical workflow. Human cardiac single-cell/single-nucleus RNA-sequencing datasets were assembled into parallel physiological aging and cardiovascular disease cohorts. Endothelial cells (ECs) were first identified from the integrated cardiac datasets, followed by LEC-specific reclustering. Downstream analyses included LEC state identification, donor-normalized state composition, marker-gene and pathway analyses, transcriptional regulatory network inference, intercellular communication analysis, and independent spatial transcriptomic assessment. Cardiovascular disease cohorts included non-failing (NF) hearts, dilated cardiomyopathy (DCM), hypertrophic cardiomyopathy (HCM), doxorubicin-associated cardiomyopathy (DoxCM), and heart failure with preserved ejection fraction (HFpEF). (B) Overview of the physiological aging and cardiovascular disease cohorts, including donor distribution, retained LEC numbers, and relevant clinical/sample characteristics. (C) Two-step EC-to-LEC identification strategy. Endothelial cells were initially resolved from the complete cardiac cellular compartment and subsequently reclustered to identify LECs based on canonical lymphatic endothelial markers, including PROX1, FLT4, LYVE1, PDPN, CCL21, and MMRN1. (D) UMAP visualization of the final LEC populations used for downstream physiological aging and cardiovascular disease analyses.

Because physiological aging and overt cardiovascular disease represent biologically distinct processes, subsequent analyses were performed in parallel rather than assuming that age-associated and disease-associated LEC states were equivalent. Within each cohort, we evaluated LEC state composition, marker genes and pathway enrichment, transcription factor regulons using SCENIC, and predicted microenvironmental communication using NicheNet/MultiNicheNet, followed by independent spatial analyses of selected findings.

### Physiological aging redistributes cardiac LECs among pre-existing functional states

Reclustering of LECs from the physiological aging cohort resolved 11 transcriptional states with distinct marker, pathway, and regulatory profiles (Fig. 2A–C, S2-3, Table S3). Both cell-level and donor-normalized analyses demonstrated that aging was associated predominantly with redistribution among existing LEC states rather than emergence of a novel aging-specific population. Young hearts contained a relatively diverse mixture of LEC states, whereas middle-aged and older hearts became increasingly dominated by a smaller subset of transcriptional programs (Fig. 2B).

**Figure 2.**
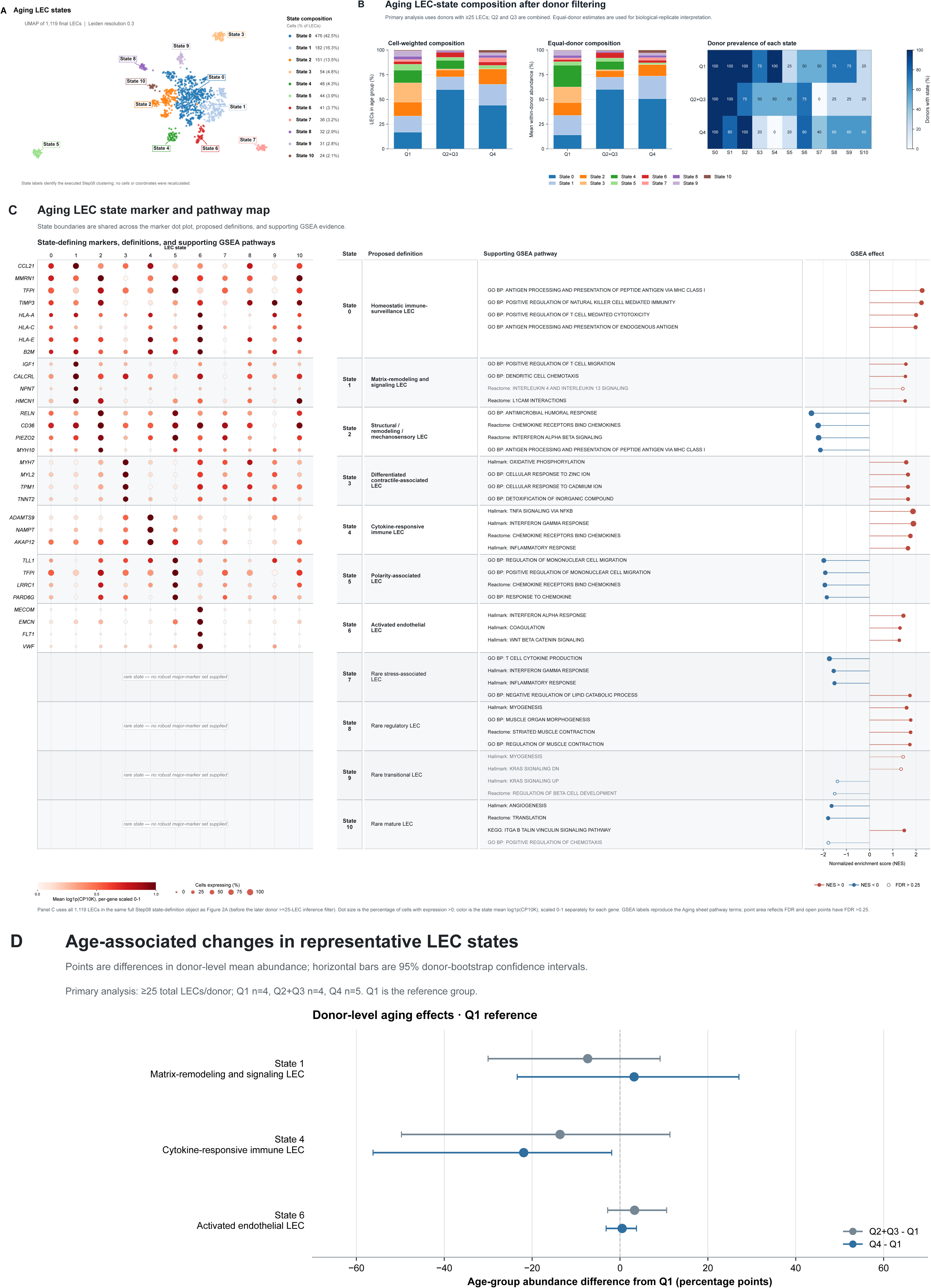
Physiological aging remodels cardiac LEC states. (A) UMAP representation of cardiac LECs from the physiological aging cohort, identifying 11 transcriptional states (States 0–10). (B) Distribution of LEC states across young (Q1), middle-aged (Q2+Q3), and old (Q4) donors. Stacked bars show donor-normalized LEC state composition by age group. (C) Molecular and functional annotation of aging-associated LEC states based on representative marker genes and pathway enrichment. State 0 was characterized as a homeostatic immune-surveillance LEC state; State 1 as a matrix-remodeling/signaling state; State 2 as a structural-remodeling/mechanosensory state; State 3 as a differentiated contractile-associated state; State 4 as a cytokine-responsive immune state; State 5 as a polarity-associated state; and State 6 as an activated endothelial state. States 7–10 represented low-frequency specialized populations. (D) Donor-level abundance of selected age-associated LEC states. State 0 increased with age, whereas cytokine-responsive State 4 and polarity-associated State 5 declined. States 1 and 2 showed increased representation in advanced aging. Each point represents a difference in equal-donor mean state abundance (Q2+Q3 minus Q1 or Q4 minus Q1), expressed in percentage points; horizontal bars indicate 95% percentile confidence intervals from 10,000 within-group donor-bootstrap resamples. Donors with ≥25 final LECs were included (Q1, n = 4; Q2+Q3, n = 4; Q4, n = 5). Pairwise comparisons used two-sided Mann– Whitney U tests with Benjamini–Hochberg correction across all 22 state-by-contrast tests (11 states × two contrasts), not only the displayed states. P values and FDR are retained in the source data; significance levels are not displayed in this panel.

The most prominent age-associated change involved State 0, a homeostatic immune-surveillance LEC state characterized by CCL21, MMRN1, TFPI, TIMP3, HLA-A, HLA-C, HLA-E, and B2M. Enrichment of antigen processing and presentation, immune-surveillance-related pathways, and translational programs further supported a mature lymphatic endothelial phenotype capable of coordinating tissue immune homeostasis. State 0 increased markedly from young to middle age and remained the predominant LEC population in older hearts, indicating that physiological aging preferentially preserves and expands a core lymphatic maintenance program (Fig. 2C,D).

In contrast, State 4 exhibited a progressive decline with age. State 4 expressed ADAMTS9, NAMPT, and AKAP12 and was enriched for interferon-γ response and JAK–STAT-related signaling, consistent with a cytokine-responsive immune LEC program. Importantly, this transcriptional profile is more consistent with endothelial responsiveness to inflammatory cues than with constitutive inflammatory activation. Its progressive depletion therefore suggests that aging may reduce the representation of LECs capable of mounting adaptive responses to inflammatory or immune signals.

A second age-sensitive population, State 5, expressed TLL1, TFPI, LRRC1, and PARD6G and was associated with endothelial polarity and structural organization. State 5 also declined with age, suggesting progressive loss of a structural maintenance program in the aging lymphatic endothelium. By comparison, States 1 and 2 showed greater representation in advanced aging. State 1 expressed IGF1, CALCRL, NPNT, and HMCN1 and was associated with Hedgehog, GPCR, and cell-migration signaling, whereas State 2 expressed RELN, CD36, PIEZO2, and MYH10, together with developmental guidance and mechanosensory programs. These populations therefore appear to represent remodeling-associated states recruited during structural adaptation of the aging cardiac microenvironment.

State 3 was preferentially represented in younger hearts and expressed several contractile-associated transcripts, including MYH7, MYL2, TPM1, and TNNT2. Because of the unusual contractile signature, we interpret this population conservatively as a differentiated contractile-associated transcriptional state rather than assigning a definitive specialized LEC function. States 6–10 were comparatively uncommon and showed limited age-dependent redistribution.

Together, these findings indicate that physiological aging does not simply induce an inflammatory or senescent LEC phenotype. Instead, aging progressively restructures the cardiac lymphatic endothelial compartment through expansion of a dominant immune-surveillance/homeostatic state, loss of cytokine-responsive and polarity-associated states, and later enrichment of remodeling-associated programs.

### Age-associated LEC states are supported by distinct transcriptional regulatory programs

We next used SCENIC to determine whether the transcriptionally defined aging states were supported by distinct regulatory architectures (Fig. 3A–C, S4)^16^. Individual LEC states showed characteristic regulon profiles, and the major state-associated regulons remained reproducible across donor-bootstrap analyses, indicating that these patterns were not driven by individual donors.

**Figure 3.**
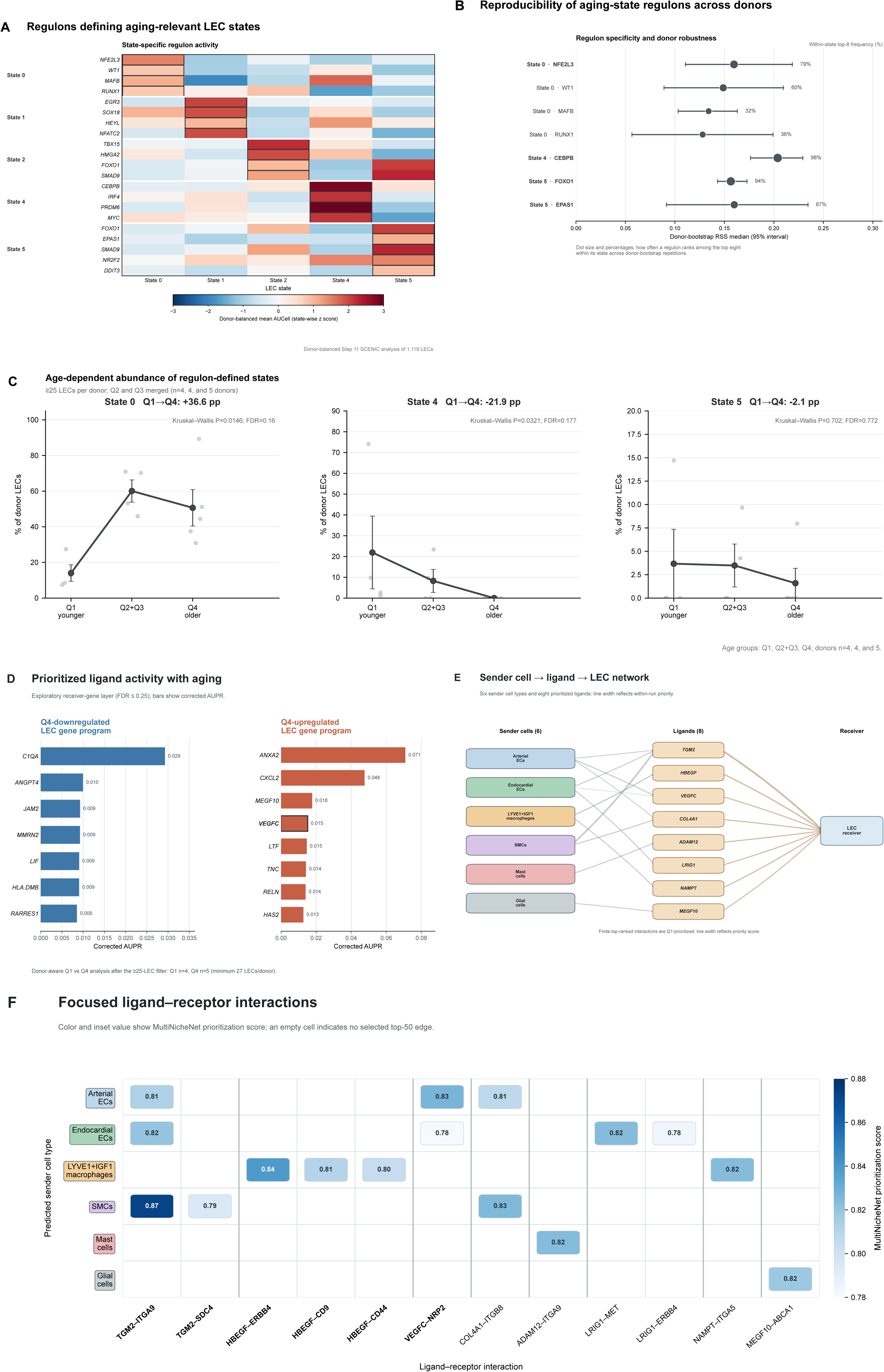
Physiological aging remodels transcriptional regulatory programs and microenvironmental signaling to cardiac LECs. (A) SCENIC analysis of transcription factor regulon activity across aging-associated LEC states. Heatmap shows state-enriched regulons, demonstrating distinct transcriptional regulatory architectures among LEC populations. (B) Representative transcription factor regulons associated with major aging-sensitive LEC states. State 0 was associated with NFE2L3, WT1, MAFB, RUNX1, and AR; State 1 with SOX18, EGR3, NFATC2, HEYL, and PLAG1; State 2 with TBX15, HMGA2, FOXO1, TCF4, NR2F1, and SMAD9; cytokine-responsive State 4 with CEBPB, IRF4, PRDM6, ETV1, and MYC; and State 5 with FOXO1, EPAS1, NR2F2, SMAD9, and DDIT3. (C) Donor-level abundance of selected LEC states across age groups. Regulon enrichment was recalculated across donor resampling iterations to assess reproducibility of state specificity and reduce sensitivity to individual donors. (D) NicheNet ligand-activity analysis of age-associated LEC transcriptional changes following MultiNicheNet-based communication analysis. Ligand activities were calculated using predict_ligand_activities for receiver-gene sets defined by FDR ≤0.25 and absolute log₂FC ≥0.25, with up to 50 genes retained per direction. (E) Major sender-cell populations associated with young- and old-enriched LEC transcriptional programs. Predicted signals originated from multiple vascular, immune, smooth-muscle, and stromal populations, demonstrating age-associated remodeling of the cardiac LEC microenvironment. (F) Selected ligand–receptor interactions prioritized by MultiNicheNet. Candidate interactions included VEGFC–NRP2 and growth-factor-, extracellular-matrix-, and vascular-remodeling-associated signaling involving TGM2, HBEGF, COL4A1, FN1, and related pathways. Interaction scores represent computational prioritization and do not establish direct biochemical signaling.

The age-expanded State 0 exhibited a regulatory program involving NFE2L3, WT1, MAFB, RUNX1, and AR, consistent with maintenance of endothelial identity, stress adaptation, and immune regulation. The preservation of this regulatory architecture despite increasing State 0 abundance suggests that aging predominantly expands an existing endothelial program rather than generating a distinct aging-specific transcriptional identity.

State 1 was associated with SOX18, EGR3, NFATC2, HEYL, and PLAG1, consistent with endothelial differentiation and signal-responsive remodeling. State 2 showed enrichment of TBX15, HMGA2, FOXO1, TCF4, NR2F1, and SMAD9, supporting a structurally adaptive program involving developmental and environmental-response pathways.

The cytokine-responsive State 4 displayed a distinct regulatory architecture that included CEBPB, IRF4, PRDM6, ETV1, and MYC, with particularly reproducible CEBPB-associated activity across donors. When considered together with interferon-γ/JAK–STAT pathway enrichment and the progressive decline of this state with age, these findings further support State 4 as a cytokine-responsive endothelial program that becomes less represented during physiological aging. State 5 showed regulatory activity involving FOXO1, EPAS1, NR2F2, SMAD9, and DDIT3, linking its structural and polarity-associated phenotype to endothelial maintenance, oxygen sensing, morphogenic signaling, and stress adaptation.

Thus, physiological aging was accompanied not only by altered LEC state composition but also by a contraction of the repertoire of specialized transcriptional regulatory programs represented within the cardiac lymphatic endothelium.

### Aging alters predicted microenvironmental signals to cardiac LECs

Because endothelial state composition is influenced by signals from surrounding cardiac cells, we next used MultiNicheNet to identify candidate microenvironmental inputs associated with age-dependent LEC transcriptional changes (Fig. 3D–F)^17^. The predicted signaling landscape differed substantially between young and aged hearts, indicating that aging affects both intrinsic LEC regulatory states and the extracellular signals to which LECs are exposed.

Signals associated with younger LEC programs were preferentially linked to vascular endothelial populations, LYVE1-positive macrophages, and other cardiac cell populations, whereas aged LEC programs received predicted input from arterial and endocardial endothelial cells, smooth muscle cells, and stromal populations. Among the prioritized interactions were VEGFC–NRP2, growth-factor-associated signaling, and extracellular matrix interactions involving molecules such as TGM2, COL4A1, and FN1.

Integration with SCENIC suggested convergence between extracellular and intracellular regulatory layers. In particular, predicted extracellular signaling capable of engaging developmental and vascular-remodeling pathways accompanied SMAD-associated regulon activity in selected LEC states. These analyses therefore support a model in which physiological aging reshapes cardiac LEC biology through coordinated alteration of both the microenvironmental signaling landscape and the intrinsic transcriptional regulatory repertoire, rather than through a single dominant age-associated signaling pathway.

### Cardiovascular diseases redistribute cardiac LECs through shared and disease-specific remodeling programs

We next examined whether pathological cardiac remodeling reproduced the aging trajectory or generated distinct LEC responses. Reclustering of LECs across NF, DCM, HCM, DoxCM, and HFpEF hearts identified nine disease-cohort LEC states (Fig. 4A–C, S5, Table S4). As in physiological aging, cardiovascular disease was characterized primarily by redistribution among transcriptional states rather than appearance of disease-exclusive cell populations. However, the pattern of redistribution differed substantially from physiological aging.

**Figure 4.**
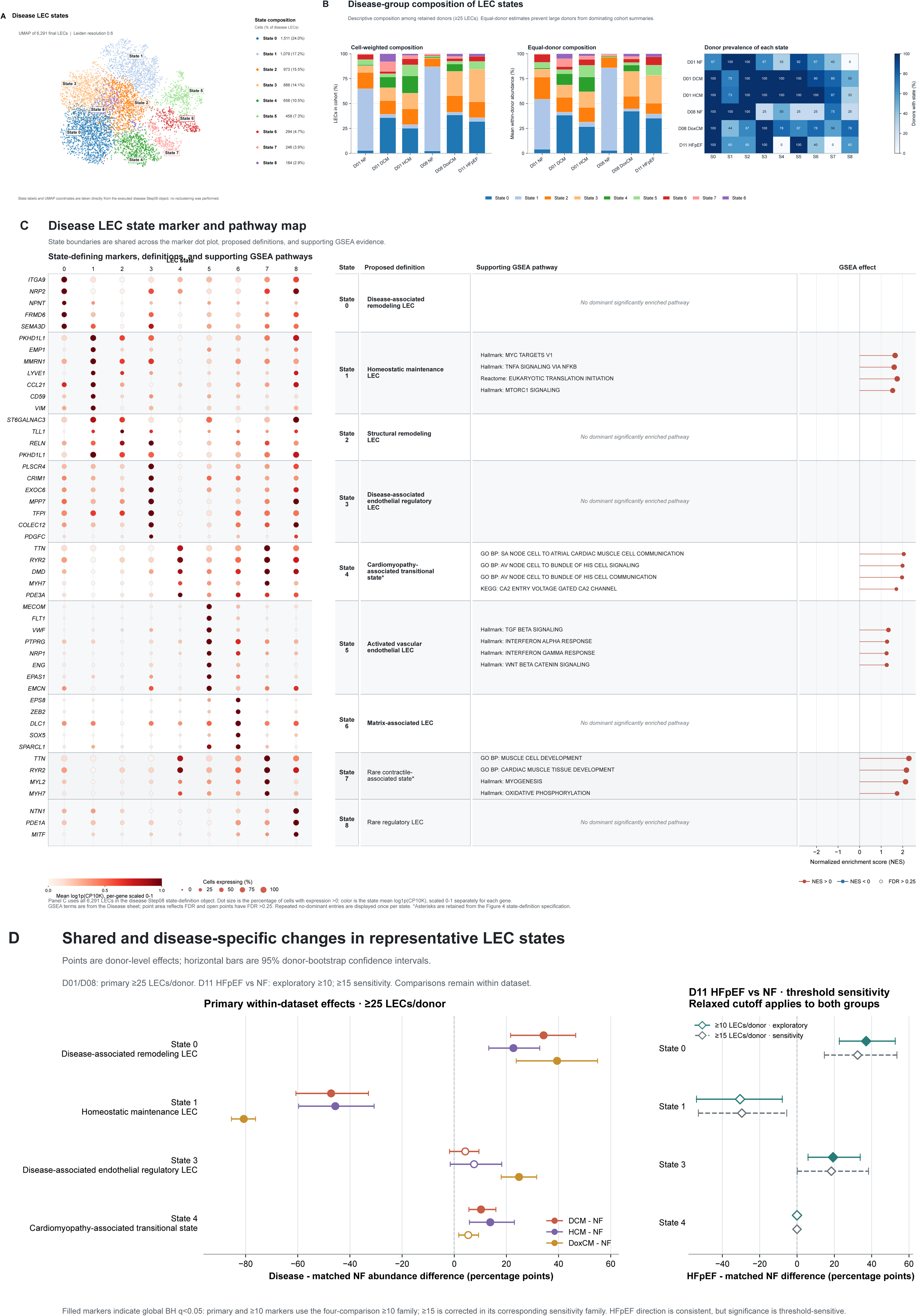
Cardiovascular diseases induce shared and disease-specific remodeling of the cardiac LEC state landscape. (A) UMAP representation of cardiac LECs from NF, DCM, HCM, DoxCM, and HFpEF hearts, identifying nine transcriptional states (States 0–8). (B) Donor-normalized LEC state composition across NF and cardiovascular disease cohorts. Diverse cardiovascular diseases were characterized by redistribution among pre-existing LEC states rather than emergence of disease-exclusive populations. (C) Molecular annotation of disease-associated LEC states based on representative marker genes and pathway enrichment. State 0 represented a disease-associated remodeling state expressing ITGA9, NRP2, NPNT, FRMD6, and SEMA3D; State 1 a homeostatic maintenance state expressing PKHD1L1, EMP1, MMRN1, LYVE1, CCL21, CD59, and VIM; State 2 a structural-remodeling state; State 3 a disease-associated endothelial regulatory state; State 4 a cardiomyopathy-associated transitional transcriptional state; State 5 an activated vascular endothelial state; and States 6–8 low-frequency specialized populations. (D) Donor-level abundance of selected LEC states across disease cohorts. Homeostatic State 1 was reduced across cardiovascular diseases, whereas remodeling-associated State 0 expanded broadly. State 3 was particularly represented in DoxCM and HFpEF, and activated State 5 was enriched in selected disease groups. State 4 was detected in DCM, HCM, and DoxCM but was minimally represented in HFpEF. Each point represents the disease-minus-NF difference in equal-donor mean state abundance, expressed in percentage points; horizontal bars indicate 95% percentile confidence intervals from 10,000 within-group donor-bootstrap resamples. Comparisons used two-sided Mann–Whitney U tests on donor-level state percentages. DCM (n = 10) and HCM (n = 15) were compared with NF donors (n = 12) within dataset01; DoxCM (n = 9) was compared with NF donors (n = 4) within dataset08. These comparisons required ≥25 final LECs per donor. Within dataset11, HFpEF versus NF was exploratory at ≥10 LECs per donor (n = 7 and 6, respectively), with ≥15 used for sensitivity analysis (n = 5 and 4); the threshold was applied to both groups. Benjamini–Hochberg correction was applied globally across the 35 testable state-by-comparison tests in the nine-state, four-comparison family, including non-displayed states. The all-zero HFpEF State 4 contrast was not tested. Primary and HFpEF ≥10 symbols use the family containing the ≥10 comparison; HFpEF ≥15 symbols use a separately corrected family with the ≥15 comparison substituted. Filled symbols indicate BH-adjusted q < 0.05; open symbols indicate q ≥ 0.05 or an untestable contrast.

The most consistent disease-associated change was depletion of State 1, which predominated in NF hearts but decreased across all four cardiovascular disease groups (Fig. 4B, D). State 1 expressed PKHD1L1, EMP1, MMRN1, LYVE1, CCL21, CD59, and VIM, and was enriched for metabolic, translational, MYC, TNFα/NFκB, mTORC1, unfolded-protein-response, oxidative-phosphorylation, and cellular stress-response pathways. We therefore interpret State 1 as a metabolically active homeostatic maintenance LEC state rather than a transcriptionally quiescent population. Its consistent depletion suggests that disruption of normal LEC homeostasis is a common feature across cardiovascular diseases.

Concomitantly, State 0 expanded across disease groups. This state expressed ITGA9, NRP2, NPNT, FRMD6, and SEMA3D, consistent with structural endothelial remodeling. Although no single pathway dominated its enrichment profile, its reproducible expansion across distinct disease etiologies indicates a shared remodeling-associated endothelial response.

State 3, characterized by PLSCR4, CRIM1, EXOC6, MPP7, TFPI, COLEC12, and PDGFC, was particularly increased in DoxCM and HFpEF, suggesting another adaptive endothelial program associated with chronic cardiac injury. In contrast, State 5 exhibited a clear activated vascular phenotype, expressing MECOM, FLT1, VWF, PTPRG, NRP1, ENG, EPAS1, and EMCN and showing enrichment of TGF-β, interferon-α, interferon-γ, and WNT/β-catenin signaling. State 5 was particularly represented in HCM and HFpEF, indicating disease-dependent recruitment of an activated endothelial program.

One notable difference among disease classes involved State 4, which was detected in DCM, HCM, and DoxCM but was minimal in HFpEF. State 4 expressed several contractile/cardiomyocyte-associated genes, including TTN, RYR2, DMD, MYH7, and PDE3A, and was enriched for cardiac-conduction-associated pathways. Accordingly, its precise cellular interpretation remains uncertain, and contributions from ambient or transitional transcriptional signals cannot be excluded. Nevertheless, its reproducible distribution across the three cardiomyopathy groups but not HFpEF indicates a disease-class-associated transcriptional pattern worthy of further investigation.

These observations demonstrate that cardiovascular diseases share a common restructuring of the LEC compartment, particularly loss of the NF-associated homeostatic State 1 and expansion of remodeling-associated states, while retaining disease-specific differences in endothelial state composition.

### Distinct regulatory networks reinforce shared and disease-specific LEC remodeling

SCENIC analysis demonstrated that disease-associated LEC states were supported by distinct and donor-reproducible transcriptional regulatory programs (Fig. 5A–C, S6). The NF-enriched State 1 was characterized by JUN, JUNB, JUND, FOS, KLF2, REL, and CEBPD regulons, consistent with a responsive endothelial program integrating environmental, inflammatory, and homeostatic signals. Its depletion across cardiovascular diseases therefore reflected loss of a coordinated regulatory state rather than reduction of individual homeostatic genes alone.

**Figure 5.**
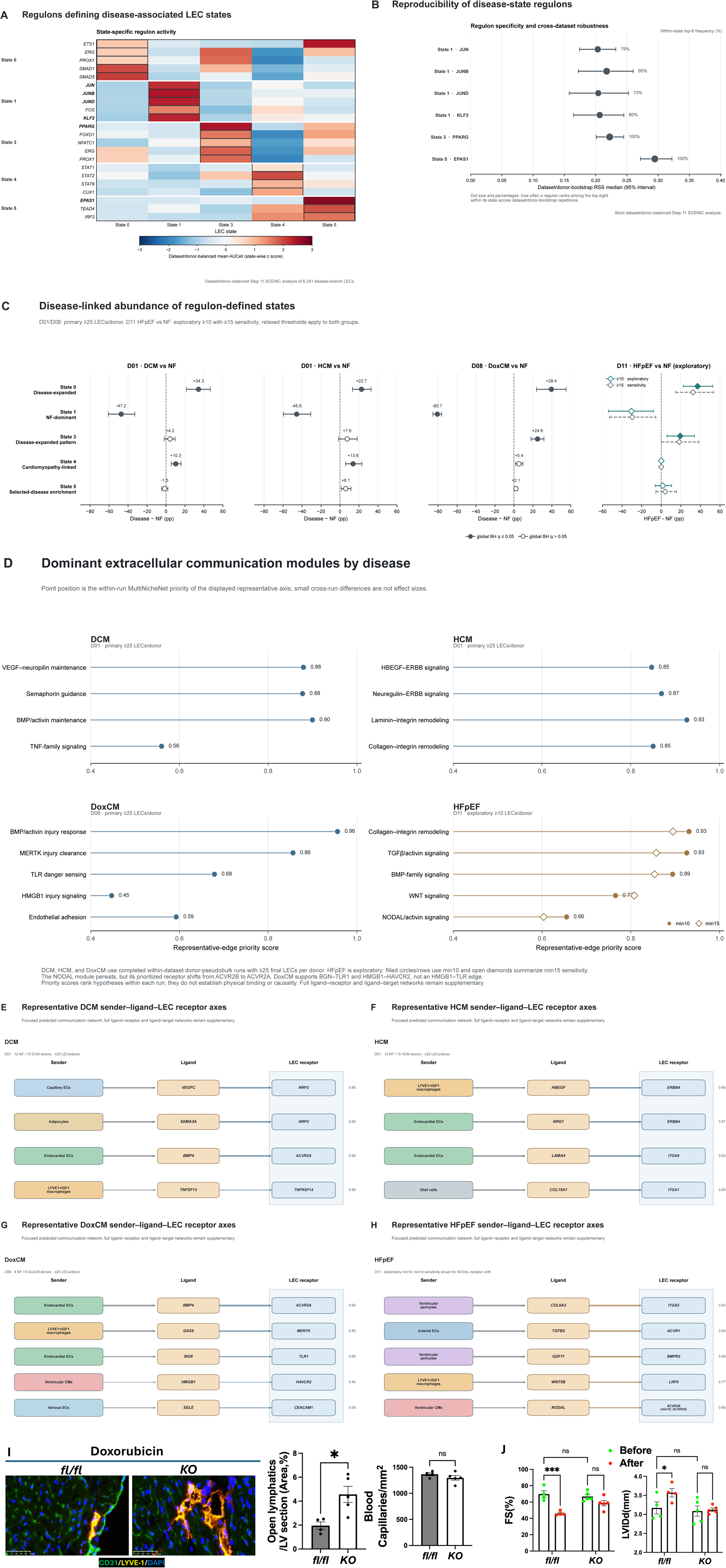
Disease-specific regulatory and microenvironmental programs identify BMP/activin–ACVR2A signaling as a candidate regulator of LEC remodeling. (A) SCENIC analysis of state-specific transcription factor regulon activity across disease-associated cardiac lymphatic endothelial cell (LEC) states. Heatmap shows representative regulons enriched within major LEC states, illustrating distinct regulatory architectures associated with homeostatic, remodeling, cardiomyopathy-associated, and activated endothelial programs. (B) Donor-bootstrap analysis of representative state-associated regulons. Points indicate estimated regulon specificity and error-corrected robustness across donor resampling; percentages indicate donor-bootstrap reproducibility. (C) Donor-level differences in the abundance of regulon-defined LEC states between cardiovascular disease groups and non-failing (NF) controls. Selected representative regulons are shown for DCM, HCM, DoxCM, and HFpEF comparisons. Filled symbols indicate significant disease-associated differences; open symbols indicate exploratory associations. (D) Dominant extracellular communication modules predicted by MultiNicheNet for DCM, HCM, DoxCM, and HFpEF. Point position represents within-run priority of representative signaling axes and is intended for within-disease interpretation rather than direct quantitative comparison across diseases. DCM was characterized by vascular-remodeling and guidance programs; HCM by growth-factor and extracellular-matrix communication; DoxCM by BMP/activin, injury-clearance, Toll-like receptor, HMGB1, and endothelial-adhesion programs; and HFpEF by extracellular-matrix, TGF-β/activin, BMP-family, WNT, and NODAL/activin-associated signaling. (E–H) Representative sender–ligand–LEC receptor axes prioritized within each disease-specific communication network. (E) DCM, showing representative vascular-guidance, extracellular-matrix, and BMP/activin-related interactions. (F) HCM, showing representative growth-factor, extracellular-matrix, and remodeling-associated interactions. (G) DoxCM, showing representative BMP/activin and injury-associated signaling, including prioritized ACVR2A-associated communication. (H) HFpEF, showing representative BMP/TGF-β/activin, developmental, and extracellular-matrix-associated interactions. Sender populations, candidate ligands, and corresponding LEC receptors are indicated. MultiNicheNet analyses represent computationally predicted communication relationships and do not establish direct biochemical ligand–receptor interactions. (I) Representative cardiac sections and vascular quantification from 18-month-old Acvr2a^fl/fl control and endothelial Acvr2a-deficient mice following doxorubicin treatment. Mice received doxorubicin (5 mg/kg once weekly for four weeks), with analysis one week after the final dose. Representative immunofluorescence images show cardiac lymphatic and blood vessels stained for LYVE-1 and CD31, respectively. Endothelial Acvr2a deletion increased the proportion of open lymphatic vessels without a corresponding difference in blood capillary density. (J) Cardiac functional responses to doxorubicin in Acvr2a^fl/fl control and endothelial Acvr2a-deficient mice. Fractional shortening (FS) and left ventricular internal diameter at end-diastole (LVIDd) were assessed before and after doxorubicin treatment. Doxorubicin induced significant functional deterioration in control mice, whereas these changes were attenuated following endothelial Acvr2a deletion. For the mouse experiments, individual points represent individual animals and data are presented as mean ± SEM; n = 4–5 per group. Statistical comparisons were performed using Student’s t-test or two-way ANOVA, as appropriate. P values or significance levels are indicated in the figure; ns, not significant.

In contrast, the disease-expanded State 0 was characterized by ETS1, ERG, PROX1, GABPA, NFIA, and SMAD1/5-associated regulatory activity, supporting preservation of endothelial and lymphatic identity together with engagement of developmental and vascular-remodeling programs. State 3 showed activity of PPARG, FOXO1, NFATC1, ERG, and PROX1, consistent with metabolic adaptation and vascular plasticity, whereas activated State 5 was characterized by EPAS1, TEAD4, IRF3, and EBF1, complementing its pathway enrichment for vascular activation and environmental stress signaling.

State 4 exhibited a distinct STAT1/STAT2/STAT6-centered regulatory signature together with CUX1 and IKZF2. Although the contractile-associated transcript profile of this state warrants caution, the presence of a coherent regulatory architecture argues against interpreting the population solely as stochastic transcriptional contamination. Instead, cardiomyopathies may preferentially engage a cytokine-responsive transitional program that is less represented in HFpEF.

Together, these findings demonstrate that disease-associated LEC states are supported by distinct regulatory architectures and that cardiovascular disease remodels the lymphatic endothelium through coordinated redistribution among transcriptionally coherent endothelial programs.

### Distinct cardiovascular diseases establish disease-specific signaling environments around cardiac LECs

We next used MultiNicheNet to determine whether the shared remodeling of LEC states occurred within common or disease-specific cardiac microenvironments (Fig. 5D–H). Despite convergence at the level of LEC state remodeling, the predicted extracellular signaling landscape differed substantially among DCM, HCM, DoxCM, and HFpEF, indicating that similar endothelial states can be engaged within biologically distinct myocardial environments.

DCM was characterized predominantly by vascular-remodeling and endothelial-guidance signaling, including VEGF receptor-associated communication together with neuropilin/semaphorin and TNF-family interactions. HCM exhibited stronger growth-factor and extracellular-matrix communication, including HBEGF–EGFR/ERBB signaling, neuropilin/semaphorin interactions, and collagen- and adhesion-associated pathways. These patterns were consistent with coordinated trophic and structural remodeling within the hypertrophic myocardial environment.

In contrast, DoxCM showed a prominent injury-response communication landscape. Prioritized interactions included HMGB1- and TLR2/TLR4-associated signaling, MERTK/PROS1-mediated injury-response pathways, CXCL12-associated communication, and integrin/adhesion networks, consistent with tissue injury and reparative responses following anthracycline exposure. HFpEF exhibited a broader chronic remodeling architecture involving BMP/TGF-β/activin-family, WNT/NODAL-related, and extracellular-matrix signaling.

Thus, although cardiovascular diseases shared redistribution of major LEC states, the extracellular environments associated with these states remained disease dependent. DCM preferentially engaged vascular guidance and remodeling, HCM growth-factor and matrix communication, DoxCM injury and repair signaling, and HFpEF chronic developmental and remodeling pathways. These findings suggest that disease-associated LEC phenotypes arise through integration of conserved endothelial response programs with distinct myocardial signaling environments.

### BMP/activin signaling emerges as a recurrent component of disease-associated LEC remodeling

In the unbiased MultiNicheNet analysis, BMP/activin-related signaling emerged among the prioritized disease-associated communication programs and was particularly prominent in DoxCM. We therefore examined this signaling family in greater detail in Fig. 5. At the regulatory level, SMAD1/5-associated activity was identified within disease-expanded State 0, linking a major remodeling-associated LEC state with transcriptional machinery responsive to BMP-family signaling. At the extracellular level, MultiNicheNet prioritized multiple components of BMP/activin-family communication across the disease cohorts, although the specific ligands, receptors, and relative contribution of this network varied among diseases.

This convergence was particularly relevant to DoxCM. In addition to its prominent injury- and innate-immune-associated signaling environment, DoxCM showed prioritized BMP/activin-family interactions, with ACVR2A among the prominent receptor components of the predicted communication network. Thus, the human DoxCM analysis suggested that anthracycline-associated injury occurs within a signaling environment in which damage-response pathways coexist with BMP/activin receptor signaling.

Together, the identification of extracellular BMP/activin-family communication and intracellular SMAD-associated regulation suggests that this signaling family represents a recurrent regulatory layer within disease-remodeled cardiac LECs rather than a pathway restricted to a single disease state. The prioritization of ACVR2A in the human DoxCM network further provided a direct rationale for examining its functional relevance in a matched experimental model of doxorubicin cardiotoxicity.

### Endothelial Acvr2a deletion preserves lymphatic remodeling and cardiac function during doxorubicin cardiotoxicity

To functionally examine the relevance of the ACVR2A-associated signaling network identified in human DoxCM, we analyzed an independent experimental model using 18-month-old aged mice with endothelial Acvr2a deletion, generated and characterized as previously described^23^, and corresponding *fl/fl* controls subjected to doxorubicin treatment (Fig. 5I,J, Table S7).

Endothelial *Acvr2a* deletion was associated with a marked increase in the proportion of open cardiac lymphatic vessels following doxorubicin exposure (Fig. 5I). In contrast, blood capillary density did not differ significantly between genotypes, indicating that the vascular phenotype was not accompanied by generalized expansion of the cardiac blood vasculature and was preferentially associated with lymphatic remodeling under these experimental conditions.

The lymphatic phenotype was accompanied by preservation of cardiac function (Fig. 5J). Doxorubicin treatment produced a marked reduction in fractional shortening in control mice, whereas this decline was attenuated in endothelial *Acvr2a*-deficient mice. Similarly, the increase in left ventricular end-diastolic dimension observed following doxorubicin exposure in control mice was not evident in the knockout group. Thus, endothelial *Acvr2a* deletion was associated with preservation of both cardiac lymphatic architecture and cardiac functional responses during cardiotoxic stress.

These experimental findings provide an orthogonal functional extension of the human DoxCM analysis. Human DoxCM LECs were embedded within an injury-associated signaling environment that included prioritized BMP/activin–ACVR2A communication, while endothelial Acvr2a perturbation modified lymphatic and cardiac responses to the same class of cardiotoxic stress in vivo. Together with the SMAD-associated regulatory architecture identified in disease-remodeled human LECs, these observations support BMP/activin–ACVR2A signaling as a candidate component of pathological lymphatic endothelial remodeling.

Because the genetic model targets the broader endothelial compartment, these experiments do not establish an LEC-autonomous function of ACVR2A. Likewise, the computational analyses do not establish a causal relationship between an individual predicted ligand and ACVR2A. Rather, the concordance between the human transcriptomic analyses and the matched *in vivo* doxorubicin model provides functional support for the biological relevance of endothelial ACVR2A within the broader BMP/activin signaling network prioritized in human cardiovascular disease.

### Independent spatial datasets support selected aging- and disease-associated LEC programs in human cardiac tissue

Finally, we asked whether selected transcriptional findings from the single-nucleus analyses were detectable in independent human spatial transcriptomic datasets^24, 25^ (Fig. 6, S7-8, Table S5).

**Figure 6.**
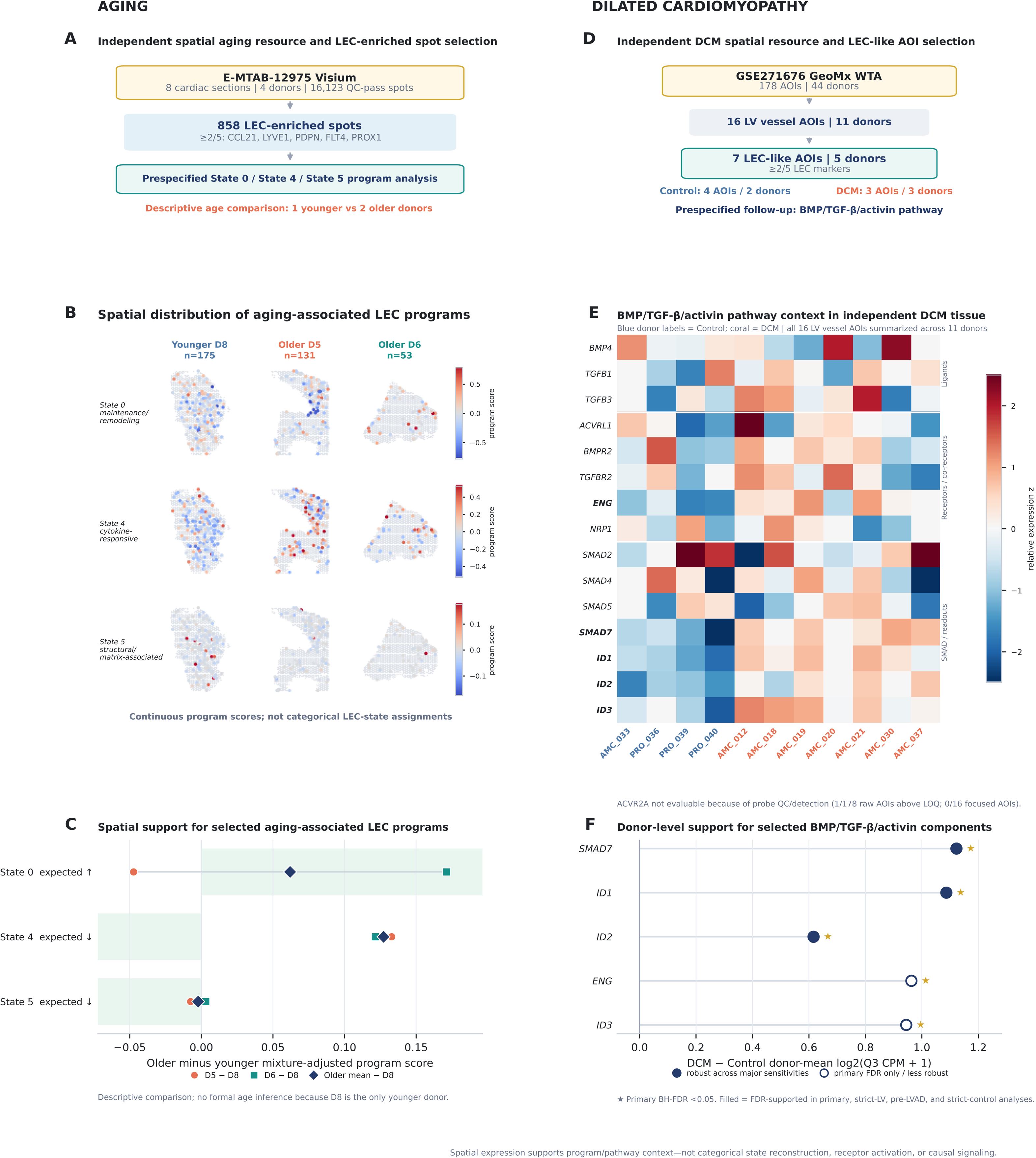
Independent human spatial transcriptomic datasets provide tissue-level support for selected aging- and disease-associated LEC programs. (A) Independent spatial transcriptomic resource and LEC-enriched spot selection for physiological aging. Eight cardiac sections from four donors in the E-MTAB-12975 Visium dataset contained 16,123 quality-control-passing spots. LEC-enriched spots were defined by detection of at least two of five lymphatic markers (CCL21, LYVE1, PDPN, FLT4, PROX1), identifying 858 spots for the primary analysis. Prespecified continuous program scores corresponding to aging-associated States 0, 4, and 5 were evaluated without categorical assignment of mixed Visium spots. The age comparison was descriptive because only one younger donor was available. (B) Spatial maps of State 0 homeostatic/maintenance, State 4 cytokine-responsive, and State 5 structural/polarity-associated program scores in the prespecified younger and older cardiac sections. Colored spots indicate LEC-enriched regions and gray spots indicate other QC-passing tissue regions. Scores represent continuous transcriptional programs within mixed Visium spots and do not represent categorical assignment of individual spots to LEC states. (C) Older-versus-younger effects for the prespecified State 0, State 4, and State 5 programs. Effects are shown separately for each older donor relative to the younger donor and for the equal-donor older mean. The expected directions derived from the snRNA-seq analysis were increased State 0 and decreased States 4 and 5 with age. No formal age-associated P value was calculated because the dataset contained only one younger donor. (D) Independent GeoMx whole-transcriptome resource and LEC-like AOI selection for DCM. The GSE271676 dataset contained 178 AOIs from 44 donors/90 cores after author-defined probe quality control. Sixteen LV vessel AOIs from 11 donors were selected, of which seven AOIs from five donors met the prespecified LEC-like criterion of at least two of five canonical LEC markers above the AOI-specific limit of quantification. The LEC-like subset included four control AOIs from two donors and three DCM AOIs from three donors. (E) Expression of selected detectable components of the BMP/TGF-β/activin signaling network in control and DCM LV vessel/LEC-like AOIs. The analysis was prespecified as a targeted follow-up of the pathway prioritized by the disease communication analysis in Fig. 5. ACVR2A was not independently evaluable because of insufficient probe detection and failure of the dataset-specific probe-quality criterion. (F) Donor-level DCM-versus-control effects for selected BMP/TGF-β/activin pathway components. Reproducible disease-associated changes included SMAD7, ID1, ID2, ENG, and ID3, with statistical significance determined across the prespecified detection-eligible family and robustness evaluated in sensitivity analyses presented in the Supplementary Data. Spatial analyses provide independent tissue-level support for selected transcriptional programs and pathway components but do not establish discrete spatial LEC states or causal intercellular signaling.

For physiological aging, an independent Visium dataset containing eight cardiac sections from four donors was analyzed using a prespecified LEC-enrichment strategy based on expression of at least two canonical lymphatic markers. This identified 858 LEC-enriched spots for spatial program analysis. Because the dataset contained only one younger donor, the analysis was deliberately treated as a descriptive tissue-level assessment rather than a formal replication of the donor-level aging analysis.

Continuous program scores corresponding to the age-associated State 0, State 4, and State 5 signatures were mapped within cardiac tissue rather than assigning mixed Visium spots to discrete LEC states. At the overall cohort level, the spatial patterns were directionally consistent with selected age-associated changes identified by snRNA-seq, including higher State 0 and lower State 4/5 program scores in older tissue. However, donor-level and sensitivity analyses showed heterogeneity among older donors, including inconsistent directionality for some programs. Accordingly, these spatial data were interpreted as descriptive tissue-level context rather than independent validation of the aging-associated state changes.

For disease validation, we analyzed an independent GeoMx whole-transcriptome dataset from control and DCM hearts. From the larger dataset, 16 LV vessel AOIs from 11 donors were identified, including seven LEC-like AOIs from five donors using prespecified lymphatic marker criteria. We specifically interrogated the BMP/TGF-β/activin pathway emerging from the disease communication analyses rather than performing another unbiased discovery analysis.

Donor-level analyses demonstrated altered expression of multiple detectable pathway components in DCM, including SMAD7, ID1, ID2, ENG, and ID3, with several signals remaining significant across prespecified sensitivity analyses. ACVR2A itself could not be independently assessed because its GeoMx probe showed insufficient detection and failed the dataset-specific probe-quality criterion. Thus, the spatial analyses provide tissue-level support for selected aging-associated LEC programs and for broader remodeling of BMP/TGF-β/activin signaling in human DCM, while avoiding inference of causal ligand–receptor signaling from spatial expression alone.

### Overall Results interpretation

Collectively, these analyses reveal that physiological aging and cardiovascular disease remodel cardiac LECs through related but non-equivalent biological processes. Physiological aging primarily redistributes LECs toward a dominant homeostatic immune-surveillance state while progressively reducing cytokine-responsive, polarity-associated, and other specialized programs.

Cardiovascular disease, in contrast, is characterized by loss of an NF-associated homeostatic state and recruitment of multiple structural, regulatory, and activated endothelial programs. Although these changes share common features across DCM, HCM, DoxCM, and HFpEF, each disease is associated with a distinct predicted microenvironmental signaling architecture. Integration of cell-state analysis, SCENIC, MultiNicheNet, independent human spatial datasets, and focused experimental validation therefore supports a model in which cardiac LEC phenotype emerges from the interaction between intrinsic transcriptional regulatory state and disease- or age-dependent microenvironmental cues, with selected pathways—including endothelial ACVR2A-associated signaling—contributing to functional lymphatic and cardiac remodeling.

## Discussion

In this study, we define the cellular, regulatory, and microenvironmental architecture of human cardiac lymphatic endothelial cells (LECs) across physiological aging and multiple cardiovascular diseases. Integration of human cardiac single-nucleus transcriptomic datasets revealed that both aging and disease remodel the lymphatic endothelial compartment predominantly through redistribution among pre-existing transcriptional states rather than generation of entirely new LEC populations. However, physiological aging and cardiovascular disease followed fundamentally different remodeling trajectories. Aging was characterized by increasing representation of a homeostatic immune-surveillance state together with progressive loss of cytokine-responsive, polarity-associated, and other specialized LEC programs. In contrast, DCM, HCM, DoxCM, and HFpEF shared depletion of a predominant non-failing homeostatic state while recruiting multiple remodeling-associated and activated endothelial programs. Integration of SCENIC and intercellular communication analyses further demonstrated that these cellular changes were accompanied by distinct transcriptional regulatory architectures and microenvironmental signaling networks. Endothelial *Acvr2a* perturbation *in vivo* and independent spatial transcriptomic analyses provided complementary support for selected components of this framework. Collectively, our findings suggest that cardiac LEC phenotype reflects the integration of intrinsic endothelial regulatory state with age- and disease-dependent microenvironmental cues, providing a framework for understanding lymphatic remodeling across the adult human cardiovascular lifespan.

### Physiological aging is associated with contraction of specialized LEC programs

A major finding of this study is that physiological aging did not produce a discrete “aged LEC” population. Instead, aging progressively altered the relative representation of existing LEC states. The dominant age-associated change was expansion of State 0, characterized by CCL21, MMRN1, TFPI, TIMP3, MHC class I genes, and transcriptional programs associated with immune surveillance and lymphatic endothelial maintenance. In parallel, several more specialized populations became less represented, most notably the cytokine-responsive State 4 and polarity-associated State 5. Thus, aging appears to preserve core lymphatic endothelial identity while narrowing the repertoire of specialized LEC programs represented within the cardiac lymphatic compartment.

This distinction may be important for understanding age-associated lymphatic dysfunction. Aging is frequently associated with chronic low-grade inflammation and vascular dysfunction, while experimental studies demonstrate age-related impairment of lymphatic transport and cardiac lymphatic remodeling^9, 10, 26^. However, our findings argue against interpreting the aging cardiac lymphatic endothelium simply as increasingly inflammatory. State 4, which was enriched for interferon-γ and JAK–STAT-related programs and regulated by a distinct cytokine-responsive transcriptional network, progressively declined with age. We therefore interpret this state as reflecting responsiveness to inflammatory cues rather than the magnitude of inflammatory exposure itself. LECs actively participate in immune-cell trafficking and inflammatory resolution rather than functioning solely as passive conduits, including within the injured heart^6, 27^. Under this model, an increasingly inflammatory aging tissue environment may coexist with reduced representation of LEC transcriptional programs associated with cytokine responsiveness. Whether this redistribution translates into impaired functional responses to inflammatory challenge will require direct experimental investigation.

The expansion of States 1 and 2 in advanced aging further suggests that aging is not simply characterized by loss of cellular complexity. These states exhibited extracellular signaling, structural remodeling, mechanosensory, and developmental-guidance programs, potentially reflecting compensatory adaptation to progressive changes in myocardial structure and extracellular matrix composition. Accordingly, the aging lymphatic endothelium may undergo a transition from a heterogeneous repertoire containing multiple specialized response states toward an organization increasingly weighted toward homeostatic maintenance and structural adaptation. Whether this redistribution preserves lymphatic function or ultimately contributes to declining lymphatic reserve will require direct functional investigation.

### Aging remodels both intrinsic LEC regulation and the surrounding cardiac microenvironment

The age-associated redistribution of LEC states was accompanied by distinct transcriptional regulatory architectures. SCENIC identified coherent state-associated regulons rather than a uniform aging transcriptional program. The age-expanded State 0 retained a stable maintenance-associated regulatory network, whereas cytokine-responsive, structural, and signaling-associated states were governed by distinct transcription factor programs. These findings suggest that aging primarily alters the representation of regulatory states rather than globally replacing the transcriptional machinery of cardiac LECs.

The intercellular communication analysis extends this concept beyond cell-intrinsic regulation. MultiNicheNet predicted substantial age-dependent reorganization of signals originating from vascular, immune, and stromal populations and directed toward LECs. Among the prioritized interactions were VEGF-C/neuropilin-related signaling and multiple extracellular matrix and growth-factor-associated interactions. Given the established importance of VEGF-C signaling in lymphatic endothelial survival, maintenance, and remodeling, the prioritization of VEGFC-associated communication provides biological plausibility to the inferred network^4, 27, 28^. However, the broader pattern is perhaps more informative than any individual ligand–receptor pair: the aging LEC phenotype appears to arise within a progressively changing multicellular cardiac environment rather than through a single cell-autonomous aging pathway.

This interpretation is also consistent with the integration of NicheNet and SCENIC. Extracellular signals predicted from surrounding cardiac populations converged with intracellular regulatory programs involving developmental, endothelial-maintenance, and SMAD-associated networks. We therefore propose that age-associated LEC remodeling reflects the interaction between microenvironmental cues and intrinsic transcriptional regulatory capacity. With aging, changes in both components may progressively alter the repertoire of endothelial states available when the heart subsequently encounters pathological stress, a possibility that will require direct functional testing.

### Cardiovascular disease remodels LECs through a trajectory distinct from physiological aging

A second major finding is that cardiovascular disease did not simply accelerate the LEC remodeling trajectory observed during physiological aging. Across DCM, HCM, DoxCM, and HFpEF, the most consistent disease-associated alteration was depletion of State 1, the predominant LEC population in non-failing hearts, together with expansion of several remodeling-associated states. State 1 retained canonical lymphatic markers while exhibiting metabolic, translational, immediate-early, and stress-responsive regulatory programs, suggesting that normal cardiac LEC homeostasis represents an active rather than transcriptionally quiescent state. Its depletion across four etiologically distinct cardiovascular diseases therefore identifies disruption of normal lymphatic endothelial homeostasis as a shared feature of pathological cardiac remodeling.

Conversely, disease-expanded State 0 retained endothelial and lymphatic regulatory programs involving ETS1, ERG, PROX1, and SMAD-associated factors, whereas State 3 exhibited regulatory features consistent with metabolic and vascular adaptation. State 5 combined an activated vascular phenotype with TGF-β, interferon, and WNT-associated pathway enrichment and a distinct EPAS1-centered regulatory architecture. These observations indicate that cardiovascular disease does not simply erode LEC identity. Rather, pathological cardiac environments redistribute LECs toward alternative endothelial programs that retain lymphatic/endothelial identity while engaging structural remodeling, metabolic adaptation, vascular activation, and stress-responsive functions.

This distinction between physiological aging and disease is conceptually important. Aging was associated predominantly with contraction of specialized adaptive programs and increasing dominance of a maintenance-oriented state, whereas cardiovascular disease recruited multiple remodeling-associated regulatory programs. We therefore propose that physiological aging and pathological remodeling represent related but non-equivalent dimensions of cardiac lymphatic biology. Aging may alter the baseline repertoire and responsiveness of LEC states, whereas cardiovascular disease imposes additional signals that redirect LECs toward context-dependent pathological or compensatory programs. Longitudinal studies will be required to determine whether aging directly constrains subsequent disease-associated state transitions, but the present data establish that the two processes should not be considered interchangeable.

### Disease-specific microenvironments shape distinct LEC responses

Despite shared remodeling at the cell-state level, the predicted extracellular signaling environments differed substantially among cardiovascular diseases. MultiNicheNet identified BMP/activin-related signaling across multiple disease contexts, together with additional signaling modules whose relative prominence differed according to disease etiology. Thus, disease specificity was reflected not simply by the presence or absence of individual pathways, but by the broader combinations of extracellular cues predicted to regulate LECs.

DCM was characterized by prominent vascular-guidance and remodeling signals, including VEGF receptor and neuropilin/semaphorin pathways, together with TNF-family, extracellular-matrix, and BMP/activin-related communication^29^. HCM similarly exhibited BMP/activin-family signaling within a broader network enriched for growth-factor and extracellular-matrix communication, including HBEGF–EGFR/ERBB, neuropilin/semaphorin, collagen, and adhesion-associated interactions. DoxCM exhibited a particularly injury-oriented signaling environment, with HMGB1-, Toll-like receptor-, MERTK/PROS1-, CXCL12-, and adhesion-associated communication occurring alongside prominent BMP/activin-family receptor signaling, including ACVR2A-associated interactions. HFpEF also showed broad BMP/TGF-β/activin-family communication embedded within a broader chronic remodeling network involving WNT/NODAL-related, developmental, and extracellular-matrix pathways.

These patterns indicate that cardiovascular diseases do not expose LECs to entirely unrelated signaling environments. Rather, shared signaling components are embedded within distinct disease-specific communication networks. DCM preferentially combines these signals with vascular guidance and remodeling, HCM with growth-factor and matrix communication, DoxCM with tissue-injury and repair programs, and HFpEF with chronic developmental and extracellular-matrix remodeling. These differences broadly parallel the distinct pathophysiological environments represented by the disease cohorts. Anthracycline cardiotoxicity involves direct cellular injury together with inflammatory and reparative responses^14^, whereas HCM is characterized by chronic structural and mechanical remodeling^30^, and HFpEF by prolonged cardiometabolic, vascular, inflammatory, and fibrotic stress^15, 31^. Our findings therefore suggest that cardiac LECs integrate both conserved and disease-specific extracellular signals to generate context-dependent transcriptional and regulatory responses rather than adopting a universal pathological lymphatic phenotype.

The distinction between cardiomyopathies and HFpEF was also evident at the cell-state level. State 4 was represented in DCM, HCM, and DoxCM but was minimal in HFpEF. Because this state contains multiple cardiomyocyte-associated transcripts, including TTN, RYR2, DMD, and MYH7, its biological identity requires particular caution. Contributions from ambient RNA, doublets, or transcriptional signals arising from close cardiomyocyte–endothelial interactions cannot be excluded. Nevertheless, its reproducible disease distribution together with a coherent STAT-centered regulatory architecture suggests that the signal is unlikely to reflect stochastic transcriptional noise alone. We therefore regard State 4 conservatively as a cardiomyopathy-associated transitional transcriptional state rather than assigning it a definitive LEC identity or function.

### BMP/activin signaling emerges as a recurrent regulatory component of disease-associated LEC remodeling

Against this background of disease-specific communication, integration of MultiNicheNet and SCENIC identified BMP/activin signaling as a recurrent component of disease-associated LEC remodeling. Rather than being restricted to single disease context, BMP/activin-family interactions were represented across disease cohorts, although the specific ligands, receptors, and relative contribution of this network varied among diseases. Complementing these extracellular predictions, SCENIC identified SMAD1/5-associated regulatory activity within disease-expanded State 0, providing an intracellular regulatory correlate of BMP-family signaling. Together, these observations identify BMP/activin signaling as a candidate shared regulatory layer embedded within otherwise distinct disease-specific cardiac microenvironments^32, 33^.

This convergence was particularly relevant to DoxCM. In addition to its prominent injury- and innate-immune-associated communication network, the human DoxCM analysis prioritized BMP/activin-family interactions, with ACVR2A emerging as a prominent receptor component of the predicted signaling network. This finding provided a disease-matched connection between the human computational analysis and our independent experimental model of anthracycline cardiotoxicity. Rather than implying that ACVR2A represents the sole mediator of the broader BMP/activin network, these data nominate ACVR2A as one candidate receptor through which this recurrent signaling family may influence endothelial responses to cardiotoxic injury.

Consistent with this interpretation, endothelial *Acvr2a* deletion in aged mice subjected to doxorubicin was associated with attenuation of doxorubicin-associated cardiac dysfunction and an increased proportion of open cardiac lymphatic vessels, without a corresponding increase in blood-vessel abundance. The preferential lymphatic phenotype is notable because it connects perturbation of a receptor prioritized from the human disease signaling network with altered cardiac lymphatic remodeling in vivo. Together, the human DoxCM analysis and matched mouse experiment therefore provide complementary evidence linking endothelial ACVR2A-associated signaling to the lymphatic and cardiac response to anthracycline injury.

Several qualifications remain important. The endothelial genetic model targets the broader vascular endothelium and therefore does not establish an LEC-autonomous requirement for ACVR2A. Similarly, MultiNicheNet predicts potential communication relationships and does not demonstrate that a specific ligand directly activates ACVR2A in human cardiac LECs. The mouse experiment should therefore not be interpreted as validation of an individual computationally predicted ligand–receptor interaction. Rather, it provides orthogonal functional support for the biological relevance of endothelial ACVR2A within the broader BMP/activin signaling network prioritized by the human disease analyses. Defining the responsible ligand sources, endothelial cell specificity, and downstream signaling mechanisms will require dedicated mechanistic studies.

### Independent spatial analyses provide orthogonal tissue-level support while revealing important limitations

Spatial transcriptomic analyses were used as an independent tissue-level assessment of selected findings rather than as a second discovery dataset. In the aging Visium dataset, continuous scores corresponding to States 0, 4, and 5 could be localized within LEC-enriched cardiac regions and showed directional support for selected age-associated changes identified by single-nucleus analysis. However, the dataset contained only one younger donor, precluding formal age inference, and Visium spots represent mixtures of neighboring cell populations. Accordingly, these results support the presence and spatial distribution of selected transcriptional programs but cannot establish discrete spatial LEC states or independently confirm population-level age effects.

The independent DCM GeoMx dataset similarly provided tissue-level evidence for remodeling of detectable components of the BMP/TGF-β/activin pathway. Several downstream or pathway-associated genes showed reproducible donor-level differences, although the relatively small LEC-like subset limits inference. Importantly, ACVR2A itself could not be evaluated because of inadequate probe performance and detection, preventing direct spatial confirmation of the receptor identified in the human disease communication analysis and subsequently examined in vivo. We therefore view the spatial analyses as orthogonal support for selected transcriptional and pathway-level observations, rather than evidence for causal cell–cell signaling.

### Study strengths and limitations

Several features strengthen the present study. First, we analyzed human cardiac LECs across both physiological aging and multiple forms of cardiovascular disease, allowing aging-associated remodeling to be distinguished from pathological remodeling rather than assuming that disease represents accelerated aging. Second, donor-normalized analyses reduced the potential for large individual samples to dominate cell-level comparisons. Third, integration of marker expression, pathway enrichment, SCENIC, and NicheNet/MultiNicheNet provided complementary cellular, regulatory, and microenvironmental perspectives on LEC heterogeneity. Finally, selected observations were evaluated using orthogonal approaches, including endothelial genetic perturbation in vivo and independent human spatial transcriptomic datasets.

Several limitations should also be acknowledged. The study integrates datasets generated across different cohorts and experimental platforms, and residual technical or cohort-specific effects cannot be completely eliminated. Human cardiac datasets are cross-sectional and therefore cannot determine whether individual LEC states transition sequentially during aging or disease. Some rare states are represented by relatively few cells or donors, and states containing contractile-associated transcripts require cautious interpretation because ambient RNA or other technical contributions cannot be excluded. SCENIC and NicheNet infer regulatory and intercellular relationships from transcriptomic data and therefore identify candidate mechanisms rather than direct biochemical interactions. The endothelial Acvr2a model does not distinguish LEC-autonomous from broader vascular effects and was evaluated in a focused experimental cohort. Finally, the independent spatial datasets have limited cellular resolution and donor representation, particularly for physiological aging, and therefore provide supportive rather than definitive validation.

## Conclusions

In summary, our study reveals that the human cardiac lymphatic endothelium undergoes substantial but context-dependent remodeling across physiological aging and cardiovascular disease. Physiological aging is characterized by increasing dominance of a homeostatic immune-surveillance program together with progressive loss of specialized cytokine-responsive and structural LEC states, accompanied by diminished functional adaptation to an inflammatory circulating environment. Cardiovascular diseases follow a distinct trajectory characterized by loss of a non-failing homeostatic state and recruitment of shared and disease-specific remodeling programs shaped by different myocardial microenvironments. Integration of transcriptional regulatory and intercellular communication analyses identifies candidate pathways connecting these extracellular environments to LEC state remodeling, with independent spatial analyses and focused experimental studies providing orthogonal support for selected findings.

Together, these observations support a model in which cardiac LEC identity is dynamic rather than fixed and reflects the interaction between intrinsic transcriptional state and the surrounding myocardial microenvironment. Aging appears to narrow the repertoire of specialized adaptive LEC programs, whereas cardiovascular diseases redirect the remaining endothelial landscape through disease-specific signaling environments. Defining how these state transitions influence lymphatic transport, immune resolution, and myocardial remodeling may provide new opportunities to preserve lymphatic endothelial function and cardiovascular resilience across aging and disease.

## METHODS

### Code transparency

Custom code supporting the principal computational and statistical analyses is being deposited in the study GitHub repository(https://github.com/Jack2825/LEC-research), and the complete analysis code will be made publicly available upon publication. Publicly available human transcriptomic datasets can be accessed through their original repositories using the accession numbers listed in Supplementary Table S1. Software and package versions, major analytical parameters, and additional information supporting computational reproducibility are provided in Supplementary Table S6 and the Supplementary Methods.

### Study design and human cardiac transcriptomic datasets

Publicly available human cardiac single-cell and single-nucleus RNA-sequencing datasets were integrated to characterize lymphatic endothelial cell (LEC) heterogeneity across physiological aging and cardiovascular disease. Physiological aging was examined in non-failing human hearts spanning the adult age range, whereas disease analyses included non-failing (NF) controls and hearts from patients with dilated cardiomyopathy (DCM), hypertrophic cardiomyopathy (HCM), doxorubicin-associated cardiomyopathy (DoxCM), and heart failure with preserved ejection fraction (HFpEF). Because physiological aging and overt cardiovascular disease represent biologically distinct processes, aging and disease cohorts were analyzed independently. Dataset sources, accession numbers, donor characteristics, and inclusion criteria are provided in Supplementary Table S1 and the Supplementary Methods.

### Identification and characterization of cardiac LEC states

Following dataset-specific quality control and integration, endothelial cells were identified from the complete cardiac cell populations and subsequently reclustered to resolve cardiac LECs. LEC identity was established using canonical lymphatic endothelial markers, including PROX1, FLT4, LYVE1, PDPN, CCL21, and MMRN1, together with exclusion of non-lymphatic endothelial and other cardiac cell populations. LECs from the physiological aging and disease cohorts were independently reclustered and characterized using marker-gene expression and pathway enrichment analyses. State abundance was calculated at the donor level to avoid treating individual cells as independent biological replicates. For aging analyses, non-failing donors were grouped as young, middle-aged, and older adults according to prespecified age criteria.

### Transcriptional regulatory and intercellular communication analyses

Transcriptional regulatory programs underlying LEC states were inferred using SCENIC, with donor-bootstrap analyses used to evaluate reproducibility of major state-associated regulons. Microenvironmental communication between surrounding cardiac cell populations and LECs was investigated using NicheNet/MultiNicheNet. Candidate signaling relationships were prioritized by integrating sender-cell ligand expression, receptor expression in LECs, and predicted downstream transcriptional responses. Aging and individual cardiovascular disease comparisons were evaluated separately.

### Endothelial Acvr2a deletion during doxorubicin cardiotoxicity

To functionally evaluate a disease-associated signaling pathway prioritized by the human transcriptomic analyses, 18-month-old mice with endothelial-specific Acvr2a deletion, generated and characterized as previously described^23^, and corresponding control mice were subjected to doxorubicin treatment. Cardiac function was assessed by echocardiography, and cardiac lymphatic and blood vascular phenotypes were quantified by immunostaining and image analysis. Because Acvr2a deletion targeted the broader endothelial compartment rather than LECs specifically, these experiments were interpreted as assessing the contribution of endothelial Acvr2a signaling to cardiac and lymphatic responses to doxorubicin rather than establishing an LEC-autonomous mechanism.

### Spatial transcriptomic analyses

Independent human cardiac spatial transcriptomic datasets were used for orthogonal tissue-level assessment of selected findings. For physiological aging, Visium cardiac sections from four donors were analyzed using a prespecified LEC-enrichment strategy based on canonical lymphatic markers. Continuous program scores corresponding to selected age-associated LEC states were evaluated within LEC-enriched regions rather than assigning mixed Visium spots to discrete cell states.

For disease assessment, an independent GeoMx whole-transcriptome dataset from control and DCM hearts was analyzed. Vessel-associated areas of illumination (AOIs) were screened using prespecified lymphatic marker criteria to identify LEC-like regions, and detectable components of the BMP/TGF-β/activin signaling network were evaluated at the donor level. Probe-level quality control and sensitivity analyses are described in the Supplementary Methods.

### Statistical analysis

Statistical analyses were performed using donors or independent biological samples as the experimental unit whenever appropriate. LEC-state abundance and human transcriptomic comparisons were evaluated using donor-normalized measurements rather than individual cells as independent replicates. Statistical tests were selected according to experimental design and data distribution, and multiple-comparison correction was applied where appropriate. Spatial analyses were performed using donor-level aggregation with prespecified sensitivity analyses. Exact statistical tests, sample sizes, definitions of biological replicates, and adjusted P-value procedures are provided in the Supplementary Methods and figure legends. All tests were two-sided unless otherwise indicated, with P<0.05 considered statistically significant.

## Acknowledge

This work was supported by the National Institutes of Health (NIH)/National Institute on Aging (K01AG080077 to P.X.); the NIH/National Heart, Lung, and Blood Institute (R01HL170058 to J.D.R. and R01HL159443 to P.B.Y.); the NIH/National Institute of Arthritis and Musculoskeletal and Skin Diseases (R01AR057374 to P.B.Y.); the American Heart Association (25SFRNPC-KMS1463897 to J.D.R.); and the Fred and Ines Yeatts Fund for Innovative Research (to J.D.R.).

## References

1. Brakenhielm E, Alitalo K. Cardiac lymphatics in health and disease. Nat Rev Cardiol. 2019;16(1):56–68. doi: 10.1038/s41569-018-0087-8. PubMed PMID: 30333526.

2. Heron C, Ratajska A, Brakenhielm E. Cardiac lymphatics: state of the art. Curr Opin Hematol. 2022;29(3):156–65. Epub 20220225. doi: 10.1097/MOH.0000000000000713. PubMed PMID: 35220321.

3. Card CM, Yu SS, Swartz MA. Emerging roles of lymphatic endothelium in regulating adaptive immunity. J Clin Invest. 2014;124(3):943–52. Epub 20140303. doi: 10.1172/JCI73316. PubMed PMID: 24590280; PMCID: PMC3938271.

4. Klotz L, Norman S, Vieira JM, Masters M, Rohling M, Dube KN, Bollini S, Matsuzaki F, Carr CA, Riley PR. Cardiac lymphatics are heterogeneous in origin and respond to injury. Nature. 2015;522(7554):62–7. doi: 10.1038/nature14483. PubMed PMID: 25992544; PMCID: PMC4458138.

5. Johnson LA, Jackson DG. Cell traffic and the lymphatic endothelium. Ann N Y Acad Sci. 2008;1131:119–33. doi: 10.1196/annals.1413.011. PubMed PMID: 18519965.

6. Jalkanen S, Salmi M. Lymphatic endothelial cells of the lymph node. Nat Rev Immunol. 2020;20(9):566–78. Epub 20200224. doi: 10.1038/s41577-020-0281-x. PubMed PMID: 32094869.

7. Amersfoort J, Eelen G, Carmeliet P. Immunomodulation by endothelial cells - partnering up with the immune system? Nat Rev Immunol. 2022;22(9):576–88. Epub 20220314. doi: 10.1038/s41577-022-00694-4. PubMed PMID: 35288707; PMCID: PMC8920067.

8. Litvinukova M, Talavera-Lopez C, Maatz H, Reichart D, Worth CL, Lindberg EL, Kanda M, Polanski K, Heinig M, Lee M, Nadelmann ER, Roberts K, Tuck L, Fasouli ES, DeLaughter DM, McDonough B, Wakimoto H, Gorham JM, Samari S, Mahbubani KT, Saeb-Parsy K, Patone G, Boyle JJ, Zhang H, Zhang H, Viveiros A, Oudit GY, Bayraktar OA, Seidman JG, Seidman CE, Noseda M, Hubner N, Teichmann SA. Cells of the adult human heart. Nature. 2020;588(7838):466–72. Epub 20200924. doi: 10.1038/s41586-020-2797-4. PubMed PMID: 32971526; PMCID: PMC7681775.

9. Ungvari Z, Tarantini S, Kiss T, Wren JD, Giles CB, Griffin CT, Murfee WL, Pacher P, Csiszar A. Endothelial dysfunction and angiogenesis impairment in the ageing vasculature. Nat Rev Cardiol. 2018;15(9):555–65. doi: 10.1038/s41569-018-0030-z. PubMed PMID: 29795441; PMCID: PMC6612360.

10. Roh K, Li H, Freeman RN, Zazzeron L, Lee A, Zhou C, Shen S, Xia P, Guerra JRB, Sheffield C, Padera TP, Zhou Y, Kim S, Aguirre A, Houstis N, Roh JD, Ichinose F, Malhotra R, Rosenzweig A, Rhee J. Exercise-Induced Cardiac Lymphatic Remodeling Mitigates Inflammation in the Aging Heart. Aging Cell. 2025;24(6):e70043. Epub 20250313. doi: 10.1111/acel.70043. PubMed PMID: 40083143; PMCID: PMC12151892.

11. Schultheiss HP, Fairweather D, Caforio ALP, Escher F, Hershberger RE, Lipshultz SE, Liu PP, Matsumori A, Mazzanti A, McMurray J, Priori SG. Dilated cardiomyopathy. Nat Rev Dis Primers. 2019;5(1):32. Epub 20190509. doi: 10.1038/s41572-019-0084-1. PubMed PMID: 31073128; PMCID: PMC7096917.

12. Gigli M, Stolfo D, Merlo M, Sinagra G, Taylor MRG, Mestroni L. Pathophysiology of dilated cardiomyopathy: from mechanisms to precision medicine. Nat Rev Cardiol. 2025;22(3):183–98. Epub 20241011. doi: 10.1038/s41569-024-01074-2. PubMed PMID: 39394525; PMCID: PMC12046608.

13. Argiro A, Parikh V, Jurcut R, Finocchiaro G, Kaski JP, Adler E, Olivotto I. Hypertrophic cardiomyopathy. Nat Rev Dis Primers. 2025;11(1):58. Epub 20250814. doi: 10.1038/s41572-025-00643-0. PubMed PMID: 40813376.

14. Clayton ZS, Brunt VE, Hutton DA, VanDongen NS, D’Alessandro A, Reisz JA, Ziemba BP, Seals DR. Doxorubicin-Induced Oxidative Stress and Endothelial Dysfunction in Conduit Arteries Is Prevented by Mitochondrial-Specific Antioxidant Treatment. JACC CardioOncol. 2020;2(3):475–88. Epub 20200915. doi: 10.1016/j.jaccao.2020.06.010. PubMed PMID: 33073250; PMCID: PMC7561020.

15. Franssen C, Chen S, Unger A, Korkmaz HI, De Keulenaer GW, Tschope C, Leite-Moreira AF, Musters R, Niessen HW, Linke WA, Paulus WJ, Hamdani N. Myocardial Microvascular Inflammatory Endothelial Activation in Heart Failure With Preserved Ejection Fraction. JACC Heart Fail. 2016;4(4):312–24. Epub 20151209. doi: 10.1016/j.jchf.2015.10.007. PubMed PMID: 26682792.

16. Aibar S, Gonzalez-Blas CB, Moerman T, Huynh-Thu VA, Imrichova H, Hulselmans G, Rambow F, Marine JC, Geurts P, Aerts J, van den Oord J, Atak ZK, Wouters J, Aerts S. SCENIC: single-cell regulatory network inference and clustering. Nat Methods. 2017;14(11):1083–6. Epub 20171009. doi: 10.1038/nmeth.4463. PubMed PMID: 28991892; PMCID: PMC5937676.

17. Browaeys R, Saelens W, Saeys Y. NicheNet: modeling intercellular communication by linking ligands to target genes. Nat Methods. 2020;17(2):159–62. Epub 20191209. doi: 10.1038/s41592-019-0667-5. PubMed PMID: 31819264.

18. Chaffin M, Papangeli I, Simonson B, Akkad AD, Hill MC, Arduini A, Fleming SJ, Melanson M, Hayat S, Kost-Alimova M, Atwa O, Ye J, Bedi KC, Jr., Nahrendorf M, Kaushik VK, Stegmann CM, Margulies KB, Tucker NR, Ellinor PT. Single-nucleus profiling of human dilated and hypertrophic cardiomyopathy. Nature. 2022;608(7921):174–80. Epub 20220622. doi: 10.1038/s41586-022-04817-8. PubMed PMID: 35732739; PMCID: PMC12591363.

19. Koenig AL, Shchukina I, Amrute J, Andhey PS, Zaitsev K, Lai L, Bajpai G, Bredemeyer A, Smith G, Jones C, Terrebonne E, Rentschler SL, Artyomov MN, Lavine KJ. Single-cell transcriptomics reveals cell-type-specific diversification in human heart failure. Nat Cardiovasc Res. 2022;1(3):263–80. Epub 20220316. doi: 10.1038/s44161-022-00028-6. PubMed PMID: 35959412; PMCID: PMC9364913.

20. Reichart D, Lindberg EL, Maatz H, Miranda AMA, Viveiros A, Shvetsov N, Gartner A, Nadelmann ER, Lee M, Kanemaru K, Ruiz-Orera J, Strohmenger V, DeLaughter DM, Patone G, Zhang H, Woehler A, Lippert C, Kim Y, Adami E, Gorham JM, Barnett SN, Brown K, Buchan RJ, Chowdhury RA, Constantinou C, Cranley J, Felkin LE, Fox H, Ghauri A, Gummert J, Kanda M, Li R, Mach L, McDonough B, Samari S, Shahriaran F, Yapp C, Stanasiuk C, Theotokis PI, Theis FJ, van den Bogaerdt A, Wakimoto H, Ware JS, Worth CL, Barton PJR, Lee YA, Teichmann SA, Milting H, Noseda M, Oudit GY, Heinig M, Seidman JG, Hubner N, Seidman CE. Pathogenic variants damage cell composition and single cell transcription in cardiomyopathies. Science. 2022;377(6606):eabo1984. Epub 20220805. doi: 10.1126/science.abo1984. PubMed PMID: 35926050; PMCID: PMC9528698.

21. Hahn VS, Chaffin M, Simonson B, Jenkin SC, Mulligan AS, Rezaee M, Bedi KC, Jr., Margulies KB, Klattenhoff CA, Sharma K, Kass DA, Ellinor PT. Single-Cell Analysis of Human Heart Failure With Preserved Ejection Fraction. Circ Res. 2026;139(2):e327433. Epub 20260515. doi: 10.1161/CIRCRESAHA.125.327433. PubMed PMID: 42137938; PMCID: PMC13286016.

22. Guo Z, Ataran A, Ma P, Yu W, Hajirezaei H, Valenzuela Ripoll C, Pedersen LN, Rashidi O, Chan MM, Shen M, Kelley S, Sargazi A, Ozcan M, Lotfinaghsh A, Cho Y, Diab A, Grogan F, Klaas A, Pompian A, Imam A, Lodhi R, Zelleke AB, Kovacs A, Margulies KB, Szymanski J, Sardiello M, Razani B, Asnani A, Kopecky BJ, Signore PE, Yi BA, Basson CT, Ravichandran KS, Prabhu SD, Bergom C, Schilling JD, Lavine KJ, Javaheri A. Human Single-Nucleus RNA Sequencing Identifies CD47 as a Therapeutic Target for Doxorubicin-Induced Cardiomyopathy. Circulation. 2025;152(10):661–81. Epub 20250814. doi: 10.1161/CIRCULATIONAHA.124.071217. PubMed PMID: 40808662; PMCID: PMC12756910.

23. Xia P, Lee S, Roh K, Griffith J, Zhou Y, Guzman E, Shi Y, Yang Z, Castro C, Li H, Guo YY, Singh A, Knipe RS, Raji I, Xu JH, Babbs RK, Fisher F, Lachey J, Seehra J, Yu PB, Lee SJ, Anderson DG, Aguirre A, Rosenzweig A, Malhotra R, Roh JD. Endothelial ActRIIA inhibition protects the cardiac microvasculature in severe viral respiratory infection. Res Sq. 2025. Epub 20250401. doi: 10.21203/rs.3.rs-6306417/v1. PubMed PMID: 40235477; PMCID: PMC11998776.

24. Kanemaru K, Cranley J, Muraro D, Miranda AMA, Ho SY, Wilbrey-Clark A, Patrick Pett J, Polanski K, Richardson L, Litvinukova M, Kumasaka N, Qin Y, Jablonska Z, Semprich CI, Mach L, Dabrowska M, Richoz N, Bolt L, Mamanova L, Kapuge R, Barnett SN, Perera S, Talavera-Lopez C, Mulas I, Mahbubani KT, Tuck L, Wang L, Huang MM, Prete M, Pritchard S, Dark J, Saeb-Parsy K, Patel M, Clatworthy MR, Hubner N, Chowdhury RA, Noseda M, Teichmann SA. Spatially resolved multiomics of human cardiac niches. Nature. 2023;619(7971):801–10. Epub 20230712. doi: 10.1038/s41586-023-06311-1. PubMed PMID: 37438528; PMCID: PMC10371870.

25. Lee SE, Joo JH, Hwang HS, Chen SF, Evans D, Lee KY, Kim KH, Hyun J, Kim MS, Jung SH, Kim JJ, Lee JS, Torkamani A. Spatial transcriptional landscape of human heart failure. Eur Heart J. 2025;46(31):3098–114. doi: 10.1093/eurheartj/ehaf272. PubMed PMID: 40335066; PMCID: PMC12349961.

26. Zolla V, Nizamutdinova IT, Scharf B, Clement CC, Maejima D, Akl T, Nagai T, Luciani P, Leroux JC, Halin C, Stukes S, Tiwari S, Casadevall A, Jacobs WR, Jr., Entenberg D, Zawieja DC, Condeelis J, Fooksman DR, Gashev AA, Santambrogio L. Aging-related anatomical and biochemical changes in lymphatic collectors impair lymph transport, fluid homeostasis, and pathogen clearance. Aging Cell. 2015;14(4):582–94. Epub 20150515. doi: 10.1111/acel.12330. PubMed PMID: 25982749; PMCID: PMC4531072.

27. Vieira JM, Norman S, Villa Del Campo C, Cahill TJ, Barnette DN, Gunadasa-Rohling M, Johnson LA, Greaves DR, Carr CA, Jackson DG, Riley PR. The cardiac lymphatic system stimulates resolution of inflammation following myocardial infarction. J Clin Invest. 2018;128(8):3402–12. Epub 20180709. doi: 10.1172/JCI97192. PubMed PMID: 29985167; PMCID: PMC6063482.

28. Heron C, Dumesnil A, Houssari M, Renet S, Lemarcis T, Lebon A, Godefroy D, Schapman D, Henri O, Riou G, Nicol L, Henry JP, Valet M, Pieronne-Deperrois M, Ouvrard-Pascaud A, Hagerling R, Chiavelli H, Michel JB, Mulder P, Fraineau S, Richard V, Tardif V, Brakenhielm E. Regulation and impact of cardiac lymphangiogenesis in pressure-overload-induced heart failure. Cardiovasc Res. 2023;119(2):492–505. doi: 10.1093/cvr/cvac086. PubMed PMID: 35689481; PMCID: PMC10064842.

29. Treasure CB, Vita JA, Cox DA, Fish RD, Gordon JB, Mudge GH, Colucci WS, Sutton MG, Selwyn AP, Alexander RW, et al. Endothelium-dependent dilation of the coronary microvasculature is impaired in dilated cardiomyopathy. Circulation. 1990;81(3):772–9. doi: 10.1161/01.cir.81.3.772. PubMed PMID: 2306829.

30. Petersen SE, Jerosch-Herold M, Hudsmith LE, Robson MD, Francis JM, Doll HA, Selvanayagam JB, Neubauer S, Watkins H. Evidence for microvascular dysfunction in hypertrophic cardiomyopathy: new insights from multiparametric magnetic resonance imaging. Circulation. 2007;115(18):2418–25. Epub 20070423. doi: 10.1161/CIRCULATIONAHA.106.657023. PubMed PMID: 17452610.

31. Yang JH, Obokata M, Reddy YNV, Redfield MM, Lerman A, Borlaug BA. Endothelium-dependent and independent coronary microvascular dysfunction in patients with heart failure with preserved ejection fraction. Eur J Heart Fail. 2020;22(3):432–41. Epub 20191215. doi: 10.1002/ejhf.1671. PubMed PMID: 31840366.

32. Beets K, Staring MW, Criem N, Maas E, Schellinx N, de Sousa Lopes SM, Umans L, Zwijsen A. BMP-SMAD signalling output is highly regionalized in cardiovascular and lymphatic endothelial networks. BMC Dev Biol. 2016;16(1):34. Epub 20161010. doi: 10.1186/s12861-016-0133-x. PubMed PMID: 27724845; PMCID: PMC5057272.

33. Dunworth WP, Cardona-Costa J, Bozkulak EC, Kim JD, Meadows S, Fischer JC, Wang Y, Cleaver O, Qyang Y, Ober EA, Jin SW. Bone morphogenetic protein 2 signaling negatively modulates lymphatic development in vertebrate embryos. Circ Res. 2014;114(1):56–66. Epub 20131011. doi: 10.1161/CIRCRESAHA.114.302452. PubMed PMID: 24122719; PMCID: PMC4047637.

