## Supplemental Information for "Physiological Aging and Cardiovascular Disease Distinctly Remodel the Human Cardiac Lymphatic Endothelium"

Supplementary Methods: 4,086 Words

Supplementary Figures: 8;

Supplementary Tables: 7;

### Corresponding author:

Peng Xia, PhD

Cardiovascular Research Center

Massachusetts General Brigham

Harvard Medical School

Boston, MA, USA

**Acknowledge:** This work was supported by the National Institutes of Health (NIH)/National Institute on Aging (K01AG080077 to P.X.); the NIH/National Heart, Lung, and Blood Institute (R01HL170058 to J.D.R. and R01HL159443 to P.B.Y.); the NIH/National Institute of Arthritis and Musculoskeletal and Skin Diseases (R01AR057374 to P.B.Y.); the American Heart Association (25SFRNPC-KMS1463897 to J.D.R.); and the Fred and Ines Yeatts Fund for Innovative Research (to J.D.R.).

**Conflict of Interest:** PBY is a co-founder and consultant for Keros Therapeutics, which develops therapies for cardiovascular, hematologic, and musculoskeletal diseases targeting bone morphogenetic protein and TGF-β signaling pathways. PBY is a co-founder of Modal Therapeutics, which develops therapies for vascular and metabolic diseases. The Mass General Brigham corporation has applied for patents on behalf of PBY on the use of saracatinib for the treatment of FOP. The interests of PBY are reviewed and managed by Massachusetts General Hospital in accordance with their conflict-of-interest policies. JDR has received research support from Keros Therapeutics and Amgen Pharmaceuticals and consulting fees from Takeda.

#### SUPPLEMENTARY METHODS

##### Code availability and computational reproducibility

Custom analysis code used to generate the principal computational and statistical results will be made publicly available through GitHub upon publication. Scripts will be organized according to major analytical components, including data processing and quality control, LEC-state characterization, differential-expression and pathway analyses, SCENIC, MultiNicheNet, spatial transcriptomic analyses,

Analyses were performed primarily in Python using Scanpy and related open-source packages. R was used for selected downstream analyses, including edgeR-based pseudobulk differential-expression analysis and NicheNet/MultiNicheNet analyses. Software versions and major analytical parameters are summarized in Supplementary Table S6. Statistical inference was performed at the appropriate biological-replicate level. Individual cells, imaging objects, spatial spots, or repeated AOIs were not treated as independent biological replicates when donor-, well-, or animal-level inference was required. Analysis-specific quality-control procedures, inclusion criteria, normalization approaches, statistical models, and multiple-testing corrections are described below.

Publicly available human transcriptomic datasets remain accessible through their original repositories using the accession or portal identifiers provided in Supplementary Table S1. Spatial datasets and corresponding cohort information are summarized in Supplementary Table S5.

##### Human cardiac single-cell and single-nucleus RNA-sequencing datasets

Publicly available human cardiac single-cell and single-nucleus RNA-sequencing datasets were assembled to characterize LEC heterogeneity during physiological aging and across cardiovascular diseases. Dataset-level metadata, including repository and accession or portal identifier, tissue and cardiac region, sequencing modality, disease classification, and available donor characteristics, were harmonized across studies (Supplementary Table S1). Physiological aging and cardiovascular disease atlases were analyzed independently to distinguish physiological age-associated remodeling from pathological LEC states.

For physiological aging analyses, non-failing donors were assembled from five component datasets (EGAS00001006374, ERP123138, GSE156703, GSE183852, and GSE224995). Donors were stratified into four age groups (Q1–Q4) according to the resolved donor-age distribution: Q1, 35–48 years; Q2, 48–54 years; Q3, 55–62.5 years; and Q4, 62.5–75 years. When exact age was unavailable and age was reported as a range, the midpoint of the reported range was used for age-group assignment. Donor identity and source dataset were retained throughout the analyses to enable donor-level and dataset-aware assessment of robustness.

Cardiovascular disease analyses were performed independently using datasets representing non-failing (NF) controls, dilated cardiomyopathy (DCM), hypertrophic cardiomyopathy (HCM), doxorubicin-associated cardiomyopathy (DoxCM), and heart failure with preserved ejection fraction (HFpEF). LEC-state identities derived from physiological aging were not transferred to the disease cohort. Instead, disease LECs were independently reclustered and annotated, thereby avoiding an a priori assumption that physiological aging and pathological cardiovascular remodeling generate equivalent LEC states. Detailed dataset provenance, donor characteristics, disease classifications, and cohort composition are provided in Supplementary Table S1.

##### Data preprocessing, integration, and endothelial-cell identification

Single-cell and single-nucleus RNA-sequencing datasets were processed in R (version 4.5.3), primarily using Seurat (version 5.5.1). Dataset-specific quality-control procedures were applied before integration to remove low-quality cells or nuclei according to available sequencing and feature-level metrics. Because datasets originated from multiple studies and sequencing platforms, quality-control criteria accounted for the characteristics of individual source datasets rather than imposing a single universal threshold across all cohorts. Donor-level cell numbers, endothelial-cell and LEC recovery, and analysis inclusion/exclusion information are provided in Supplementary Table S2.

Following preprocessing and integration, major cardiac cell populations were identified using established lineage-associated transcriptional markers. Endothelial cells were isolated from the integrated cardiac cellular population and independently reclustered to improve resolution of endothelial heterogeneity and facilitate identification of the relatively rare lymphatic endothelial population. Subsequent LEC reclustering and state-level analyses were conducted independently for the physiological aging and cardiovascular disease cohorts.

##### LEC identification and reclustering

Cardiac LECs were identified within the endothelial compartment using combined expression of canonical lymphatic endothelial markers, including PROX1, FLT4, LYVE1, PDPN, CCL21, and MMRN1, together with assessment of endothelial identity and potential non-endothelial transcriptional contamination. Expression of blood endothelial, mural-cell, fibroblast, immune-cell, and cardiomyocyte-associated genes was evaluated during quality control and interpretation.

For each cohort, highly variable genes were identified and dimensionality reduction was performed using 30 principal components. Harmony correction was applied to account for dataset-of-origin effects, using Project as the batch variable for the aging cohort and dataset for the disease cohort. Neighborhood graphs were constructed using 15 nearest neighbors, followed by Leiden clustering. Clustering resolutions of 0.3 and 0.6 were used for the aging and disease analyses, respectively. UMAP was used for visualization.

State-associated genes were identified using cell-level Wilcoxon rank-sum testing comparing each state with the remaining LEC population. Complete marker results are provided in Supplementary Data S1 and S5. State annotations were based on integrated consideration of marker expression, pathway enrichment, transcriptional regulatory architecture, and endothelial/lymphatic biological context. Donor distribution and resampling analyses were subsequently used to assess robustness rather than to define state identity. Final aging and disease LEC-state definitions are summarized in Supplementary Tables S3 and S4. Rare states and states containing substantial non-endothelial-associated transcriptional features were interpreted conservatively.

**Donor-level LEC-state abundance and donor inclusion**

Because the number of recovered LECs varied among donors, LEC-state abundance was calculated independently for each donor:

LEC-state abundance = number of donor LECs assigned to a given state / total number of LECs recovered from that donor.

Thus, donors rather than individual cells represented the primary biological units for state-abundance comparisons. Donor-level distributions and heatmaps were examined to determine whether apparent differences were reproducible across individuals rather than being disproportionately influenced by donors with high LEC recovery.

For primary donor-level analyses, donors with ⩾25 recovered LECs were retained to provide sufficient representation for estimation of within-donor state composition while maintaining an adequate number of independent donors for group-level comparisons. Donor-level total cell, endothelial-cell and LEC numbers and corresponding inclusion status are provided in Supplementary Table S2.

For disease-specific MultiNicheNet analyses, a minimum of 25 receiver LECs per donor was used for DCM, HCM, and DoxCM. Because LEC recovery was lower in the HFpEF cohort, the primary exploratory HFpEF analysis used a threshold of ⩾10 LECs per donor, with ⩾15 LECs per donor evaluated as a more stringent sensitivity analysis. HFpEF communication analyses were therefore interpreted cautiously. The ≥25-LEC donor threshold was applied to donor-level state-abundance comparisons and other analyses requiring donor-level representation; it was not used to reconstruct the integrated LEC atlas or rerun the primary state-marker, GSEA, or SCENIC analyses.

**Pathway and gene-set enrichment analysis**

State-associated transcriptional programs were evaluated using ranked gene-set enrichment analysis. Ranked gene lists were generated from state-versus-rest Wilcoxon marker analyses, with genes ordered according to the corresponding marker-test statistic. Enrichment was evaluated against MSigDB Hallmark, Gene Ontology Biological Process, KEGG Medicus, and Reactome gene-set collections. Gene sets containing 10–500 genes were evaluated using 1,000 permutations, and enrichment was considered statistically significant at an FDR <0.05.

Aging and disease LEC states were analyzed independently. Pathway enrichment was integrated with marker expression and complementary regulatory analyses to support biological characterization rather than being used as an independent determinant of state identity.

**SCENIC analysis**

Transcription-factor regulatory programs were evaluated using the SCENIC framework. Gene regulatory networks were inferred from LEC expression profiles, followed by motif-based evaluation of transcription factor–

target relationships and calculation of regulon activity. Regulon activity was summarized across transcriptionally defined LEC states to identify state-associated regulatory programs. Aging and disease cohorts were analyzed independently.

Robustness of state-associated regulon activity was evaluated using resampling strategies designed to reduce sensitivity to unequal cell and donor representation. For the aging cohort, equal-cell resampling was performed over 200 repetitions, together with 500 donor-bootstrap repetitions. For the disease cohort, complementary donor- and dataset-aware balancing procedures were applied, including 500 donor-bootstrap repetitions and 500 dataset–donor bootstrap repetitions. Regulons emphasized in the manuscript were selected with consideration of both state-associated activity and reproducibility across resampling strategies. Software and analytical parameters are summarized in Supplementary Table S6.

**NicheNet and MultiNicheNet analyses**

Intercellular communication between the cardiac microenvironment and LECs was evaluated using NicheNet (version 2.2.1.1) and MultiNicheNet (version 2.1.0). LECs were designated as receiver cells, while major cardiac populations were evaluated as potential sender populations. Candidate signaling relationships were prioritized by integrating ligand expression in sender populations, receptor availability in LECs, established ligand–receptor and signaling-network information, and the ability of candidate ligands to explain transcriptional changes observed in receiver LECs.

To minimize pseudoreplication, disease-associated differential-expression analyses underlying MultiNicheNet were performed using donor-level pseudobulk counts rather than treating individual cells as independent replicates. Raw integer counts were aggregated by donor, filtered using filterByExpr, normalized by trimmed mean of M-values (TMM), and analyzed using robust quasi-likelihood models in edgeR. Sender populations were required to contain >10 cells per donor and to be represented in at least four donors per comparison group. Receiver-cell inclusion thresholds were applied as described above.

For physiological aging, candidate signaling programs associated with age-dependent LEC transcriptional changes were evaluated across the aging cohort. Disease analyses were performed independently for DCM, HCM, DoxCM, and HFpEF, allowing disease-specific cardiac signaling environments to be evaluated without assuming a universal pathological communication network.

NicheNet/MultiNicheNet results were interpreted as computationally prioritized candidate signaling relationships and not as evidence of direct ligand–receptor binding, biochemical pathway activation, or causal signaling in cardiac LECs.

**Mouse studies**

**Endothelial Acvr2a deletion**

Endothelial Acvr2a-deficient mice were generated by crossing Acvr2a^fl/fl mice with Tie2-Cre mice[1]. Cre-negative Acvr2a^fl/fl littermates served as controls. Genotypes were confirmed by PCR according to established protocols. Mice were studied at approximately 18 months of age.

**Doxorubicin cardiotoxicity**

Mice received doxorubicin at 5 mg/kg by intraperitoneal (i.p.) injection once weekly for four consecutive weeks, corresponding to a cumulative dose of 20 mg/kg. Cardiac phenotyping was performed 1 day before the first doxorubicin administration (pre-treatment) and 1 week after the final doxorubicin administration (post-treatment). Cardiac vascular phenotyping was performed at the post-treatment endpoint. Individual-animal measurements underlying the cardiac functional and vascular analyses are provided in Supplementary Table S7.

**Echocardiography**

Transthoracic echocardiography was performed using a GE ultrasound imaging system in awake mice. Standard cardiac dimensions and functional parameters were acquired according to a standardized imaging protocol, including fractional shortening and the cardiac functional parameters reported in Fig. 5. Echocardiographic measurements were obtained 1 day before the first doxorubicin administration (pre-treatment) and 1 week after the final doxorubicin administration (post-treatment). Image acquisition and analysis were performed with investigators blinded to genotype when feasible. Each mouse constituted one biological replicate.

**Cardiac vascular immunostaining and quantification**

At the post-treatment endpoint, hearts were collected, embedded in OCT compound, fresh-frozen, and processed for cryosection immunostaining. Cardiac lymphatic and blood vascular structures were identified by immunostaining for LYVE1 and CD31, respectively, together with nuclear counterstaining as appropriate.

Open cardiac lymphatic vessels[2] were identified and quantified according to prespecified morphological criteria, and blood-vessel abundance was quantified independently. Measurements obtained from multiple microscopic fields and/or tissue sections from the same heart were first summarized at the individual-animal level before statistical analysis. Thus, individual imaging fields or sections were not treated as independent observations, and each mouse constituted one biological replicate.

**Spatial transcriptomic analyses**

**Aging Visium analysis**

An independent human cardiac Visium spatial transcriptomic dataset comprising eight sections from four donors was used for tissue-level assessment of aging-associated LEC programs. Sections and spatial spots were subjected to dataset-specific quality-control procedures before analysis.

LEC-enriched spots were identified using a prespecified criterion requiring detectable expression of at least two canonical lymphatic endothelial markers (CCL21, LYVE1, PDPN, FLT4, and PROX1). A total of 858 LEC-enriched spots met the primary criteria.

Because Visium spots can contain transcripts from multiple neighboring cells, LEC-enriched spots were not assigned directly to discrete snRNA-seq-defined LEC states. Instead, continuous program scores representing selected State 0-, State 4-, and State 5-associated signatures were calculated and spatially mapped. Program scores were subsequently summarized at the section and donor levels. Sensitivity analyses evaluated alternative LEC-enrichment criteria, score adjustment, spatial-block bootstrap sampling, leave-one-gene-out robustness, and potential dependence on individual donors or anatomical regions.

Because the spatial cohort contained only one younger donor, these analyses were considered supportive and descriptive rather than an independent population-level replication of age-associated differences.

**Disease GeoMx analysis**

An independent human cardiac GeoMx whole-transcriptome dataset containing control and DCM tissue was used to evaluate tissue-level support for disease-associated signaling programs. After anatomical, segmentation, and dataset-specific quality-control criteria were applied, 16 left-ventricular vessel AOIs from 11 donors comprised the primary cohort. LEC-like AOIs were identified using prespecified lymphatic-marker detection criteria, resulting in seven LEC-like AOIs from five donors.

A predefined 47-gene BMP/TGF-β/activin-family panel was interrogated, of which 16 genes passed the dataset-specific detection criteria and were retained for testing. Probe- and AOI-level quality control was performed according to the GeoMx dataset specifications, followed by donor-aware evaluation of detectable pathway components and prespecified sensitivity analyses using alternative anatomical and control definitions.

ACVR2A was represented in the original assay panel but did not satisfy the dataset-specific detection criterion and was therefore excluded from inferential spatial analysis. Failure to detect ACVR2A above the assay threshold was treated as a technical limitation and was not interpreted as evidence for biological absence of the receptor.

**Statistical analysis**

Statistical analyses were performed using R and GraphPad Prism 11.0.2. The biological unit of inference was defined according to experimental design. For human transcriptomic analyses, donors were treated as independent biological units for donor-level comparisons; individual cells were not considered independent biological replicates. For animal experiments, each mouse represented one biological replicate.

For LEC-state abundance analyses, state proportions were calculated independently for each donor before group-level comparison. The final aging pairwise analyses compared Q2+Q3 and Q4 separately with Q1 using two-sided Mann–Whitney U tests, with Benjamini–Hochberg (BH) FDR correction across all 22 state-by-contrast tests (11 states × two contrasts). Three-group omnibus comparisons used Kruskal–Wallis tests with BH correction across 11 states. Disease-versus-NF comparisons used two-sided Mann–Whitney U tests within dataset (DCM/HCM, dataset01; DoxCM, dataset08; HFpEF, dataset11). The final disease effect panels used global BH correction across all testable state-by-comparison tests, including non-displayed states, rather than the within-comparison FDR values also retained in the upstream results. This family contained 35 tests: nine states × four comparisons, excluding the all-zero HFpEF State 4 contrast. Aging and DCM/HCM/DoxCM analyses required ≥25 LECs per donor. HFpEF and its same-dataset NF controls used ≥10 LECs per donor for exploratory analysis and ≥15 for sensitivity analysis; each threshold was evaluated in a separate four-comparison BH family. Mann–Whitney tests used asymptotic P values with tie and continuity corrections. Group contrasts were reported as differences in equal-donor mean percentages, with 95% percentile confidence intervals from 10,000 bootstrap resamples of donors separately within each group. Retained donors contributed zero proportions for absent states; cells were not resampled as independent replicates. These were unadjusted donor-level group comparisons, without covariate adjustment. Multiple-testing correction was performed using the Benjamini–Hochberg procedure where applicable.

Spatial transcriptomic analyses were summarized at the donor level whenever donor replication permitted, and sensitivity analyses were used to evaluate robustness to alternative enrichment, filtering, anatomical, and cohort definitions. Data are presented as mean ± SEM or other distributional summaries as specified in the individual figure legends. Unless otherwise indicated, statistical tests were two-sided and $P<0.05$ was considered statistically significant.

**Supplementary Figure legends**

**Figure S1. Dataset-level quality control and identification of cardiac LECs.** (A) Donor representation and quality-control retention across physiological aging and cardiovascular disease datasets. (B) Endothelial cell (EC)-to-LEC selection yield across datasets, showing post-QC cells/nuclei, EC candidates, and selected LECs. (C) Distribution of final LEC numbers across LEC-positive donors in the aging and disease cohorts; donors with ⩾25 LECs were retained for donor-level analyses. (D) Cumulative donor contribution to the final aging and disease LEC atlases, illustrating the distribution of LEC representation across donors. (E) Expression of endothelial and LEC identity markers and exclusion markers in the selected LEC clusters across datasets, supporting lymphatic endothelial identity and limited contribution from non-endothelial lineages.

**Figure S2. LEC-number threshold rationale and sensitivity analyses.** (A) Effects of minimum LEC-per-donor thresholds on donor and LEC retention in the aging and disease cohorts. (B) Donor-level abundance resolution based on observed LEC counts, illustrating improved percentage resolution with increasing LEC number. (C) Sensitivity analysis of donor and LEC retention across LEC-number thresholds within individual aging groups. (D) Corresponding threshold analysis across individual disease and matched non-failing (NF) cohorts; alternative thresholds were examined for the lower-yield D11 dataset. (E) Sensitivity of representative aging- and disease-associated LEC state differences across alternative donor inclusion thresholds. Overall patterns remained directionally consistent across the evaluated thresholds, supporting the primary donor-selection strategy.

**Figure S3. Extended aging LEC-state characterization and robustness analyses.** (A) Continuous-age associations of donor-level LEC state abundance across all states; q values represent BH-adjusted FDR. (B) Maximum contribution of an individual donor to each LEC state across younger (Q1), middle-aged (Q2+Q3), and older (Q4) groups. (C) Donor-level abundance of rare LEC States 7–10 across age groups. (D) Additional marker and pathway enrichment evidence supporting characterization of rare LEC states. (E) Sensitivity analysis comparing aging-associated state-abundance differences before and after removal of the highest-contributing donor for each state.

**Figure S4. Extended aging SCENIC and MultiNicheNet analyses.** (A) Extended state-specific SCENIC regulon activity across aging-associated LEC states. (B) Donor-bootstrap analysis of additional state-associated regulons, showing median regulon specificity scores (RSS) and 95% intervals. (C) Robustness of state–regulon associations across alternative bootstrap analysis schemes. (D) Extended MultiNicheNet ligand rankings for LEC gene programs downregulated or upregulated with aging. (E) Relative contributions of sender-cell populations to prioritized sender-to-LEC communication. (F) Additional prioritized ligand–receptor and ligand–target relationships associated with aging. (G) Sensitivity analyses comparing primary donor-pseudobulk differential-expression estimates with dataset-restricted and project- and sex-adjusted models.

**Figure S5. Extended disease LEC-state characterization and robustness analyses.** (A) Additional donor-level disease effects for LEC states not highlighted in the main disease-state analysis; bars indicate 95% donor-bootstrap intervals. (B) Leave-one-donor-out analysis showing the influence of individual donors on state abundance across disease and matched non-failing (NF) cohorts. (C) Donor-level distributions of less-abundant LEC States 5–8 across cohorts. (D) Additional state-marker expression and pathway enrichment supporting disease LEC-state characterization. (E) LEC/endothelial identity and cardiac/contractile-marker detection in States 4 and 7. (F) Focused technical assessment of contractile-associated States 4 and 7, including endothelial versus cardiomyocyte signature balance, relationship of cardiac signals to doublet scores, and all-state technical context. These analyses support cautious interpretation of the contractile-associated states but do not exclude contributions from ambient RNA.

**Figure S6. Extended disease SCENIC and MultiNicheNet analyses.** (A) Extended state-specific SCENIC regulon activity across disease-associated LEC states. (B) Reproducibility of additional state-associated regulons across dataset- and donor-balancing schemes. (C) Extended MultiNicheNet ligand-activity rankings for disease-upregulated and disease-downregulated LEC gene programs across DCM, HCM, DoxCM, and HFpEF. (D) Representation of sender-cell populations among prioritized sender-to-LEC signaling axes across diseases. (E) Additional prioritized ligand–receptor and ligand–target relationships for each disease. (F) Cross-disease context of additional prioritized ligands and sensitivity analysis of HFpEF signaling priorities across alternative LEC-per-donor thresholds. DCM, HCM, and DoxCM analyses used ⩾25 LECs/donor; HFpEF analyses using ⩾ 10 LECs/donor were considered exploratory, with ⩾15 LECs/donor evaluated as a sensitivity analysis.

**Figure S7. Quality control and sensitivity analyses for spatial assessment of aging-associated LEC programs.** (A) Section-level quality-control metrics for eight Visium sections from four donors. (B) Spatial distribution of LEC-enriched spots, defined by detection of ⩾2 canonical LEC markers (CCL21, LYVE1, PDPN, FLT4, and PROX1). (C) Expression and detection frequency of canonical LEC markers before and after LEC-enrichment gating. (D) Sensitivity of LEC-enriched spot identification to alternative marker and technical quality-control criteria. (E) Sensitivity of State 0-, State 4-, and State 5-like spatial program scores to enrichment criteria, score adjustment, and spatial-block bootstrap sampling. (F) Leave-one-gene-out analyses and spatial detection frequencies of genes contributing to each program score. (G) Section-level program scores and sensitivity analyses examining older-donor dependence and paired anatomical-region effects. (H) Summary matrix of prespecified robustness and sensitivity criteria. These analyses support cautious interpretation of the spatial data: none of the three aging-associated programs met all criteria for positive spatial validation, and the availability of only one younger donor limits inference regarding age-related differences.

**Figure S8. Quality control and sensitivity analyses for spatial assessment of disease-associated BMP/TGF-β/activin signaling.** (A) Overall GeoMx dataset and AOI-level quality-control metrics. (B) Sequential anatomical, segmentation, and disease criteria used to define the primary LV vessel AOI cohort. (C) Identification of LEC-like AOIs based on detection of ⩾2 canonical lymphatic markers above the AOI-specific limit of quantification (LOQ). (D) Expression and detection frequency of canonical LEC markers in primary and LEC-like AOIs. (E) Probe detection and quality assessment for prespecified BMP/TGF-β/activin pathway components. (F) Assay coverage of ligands, receptors/co-receptors, and downstream SMAD/readout components across the BMP/TGF-β/activin pathway. (G) Donor-level sensitivity analyses of pathway-associated genes using alternative LV, pre-LVAD, and control definitions. (H) Robustness of selected pathway components across major sensitivity analyses. (I) Assessment of ACVR2A assay coverage. ACVR2A was present in the raw panel but detected above the AOI-specific LOQ in only 1 of 178 AOIs and was absent from the post-QC expression matrix; therefore, ACVR2A expression and disease-associated differences were not evaluated in this spatial dataset.

**Supplementary Tables**

Table S1. Human cardiac single-cell and single-nucleus RNA-sequencing datasets and donor characteristics.

Table S2. Donor-level quality control, endothelial and lymphatic endothelial cell numbers, and inclusion criteria.

Table S3. Final aging-associated lymphatic endothelial cell state definitions.

Table S4. Final disease-associated lymphatic endothelial cell state definitions.

Table S5. Spatial transcriptomic datasets, donors, sections/AOIs, and inclusion information.

Table S6. Software, package versions, and major analysis parameters.

Table S7. Individual-animal cardiac functional and vascular measurements from the endothelial Acvr2a doxorubicin experiment.


1. Xia P, Lee S, Roh K, Griffith J, Zhou Y, Guzman E, Shi Y, Yang Z, Castro C, Li H, Guo YY, Singh A, Knipe RS, Raji I, Xu JH, Babbs RK, Fisher F, Lachey J, Seehra J, Yu PB, Lee SJ, Anderson DG, Aguirre A, Rosenzweig A, Malhotra R, Roh JD. Endothelial ActRIIA inhibition protects the cardiac microvasculature in severe viral respiratory infection. Res Sq. 2025. Epub 20250401. doi: 10.21203/rs.3.rs-6306417/v1. PubMed PMID: 40235477; PMCID: PMC11998776.
2. Henri O, Pouehe C, Houssari M, Galas L, Nicol L, Edwards-Levy F, Henry JP, Dumesnil A, Boukhalfa I, Banquet S, Schapman D, Thuillez C, Richard V, Mulder P, Brakenhielm E. Selective Stimulation of Cardiac Lymphangiogenesis Reduces Myocardial Edema and Fibrosis Leading to Improved Cardiac Function Following Myocardial Infarction. Circulation. 2016;133(15):1484-97; discussion 97. Epub 20160301. doi: 10.1161/CIRCULATIONAHA.115.020143. PubMed PMID: 26933083.

**Figure S1 | Dataset-level QC and LEC identification**

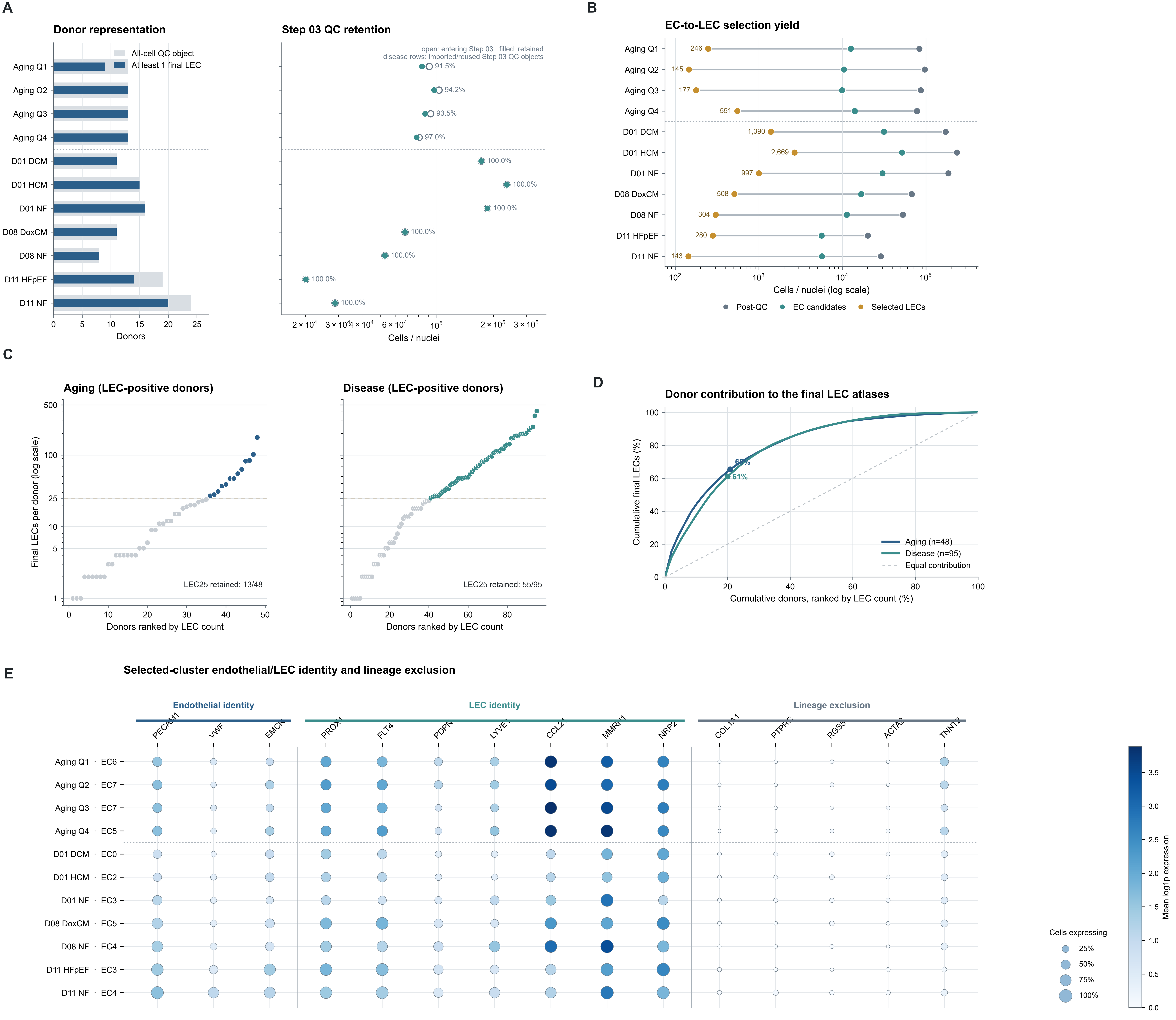

Figure S2 | LEC-number threshold rationale and sensitivity

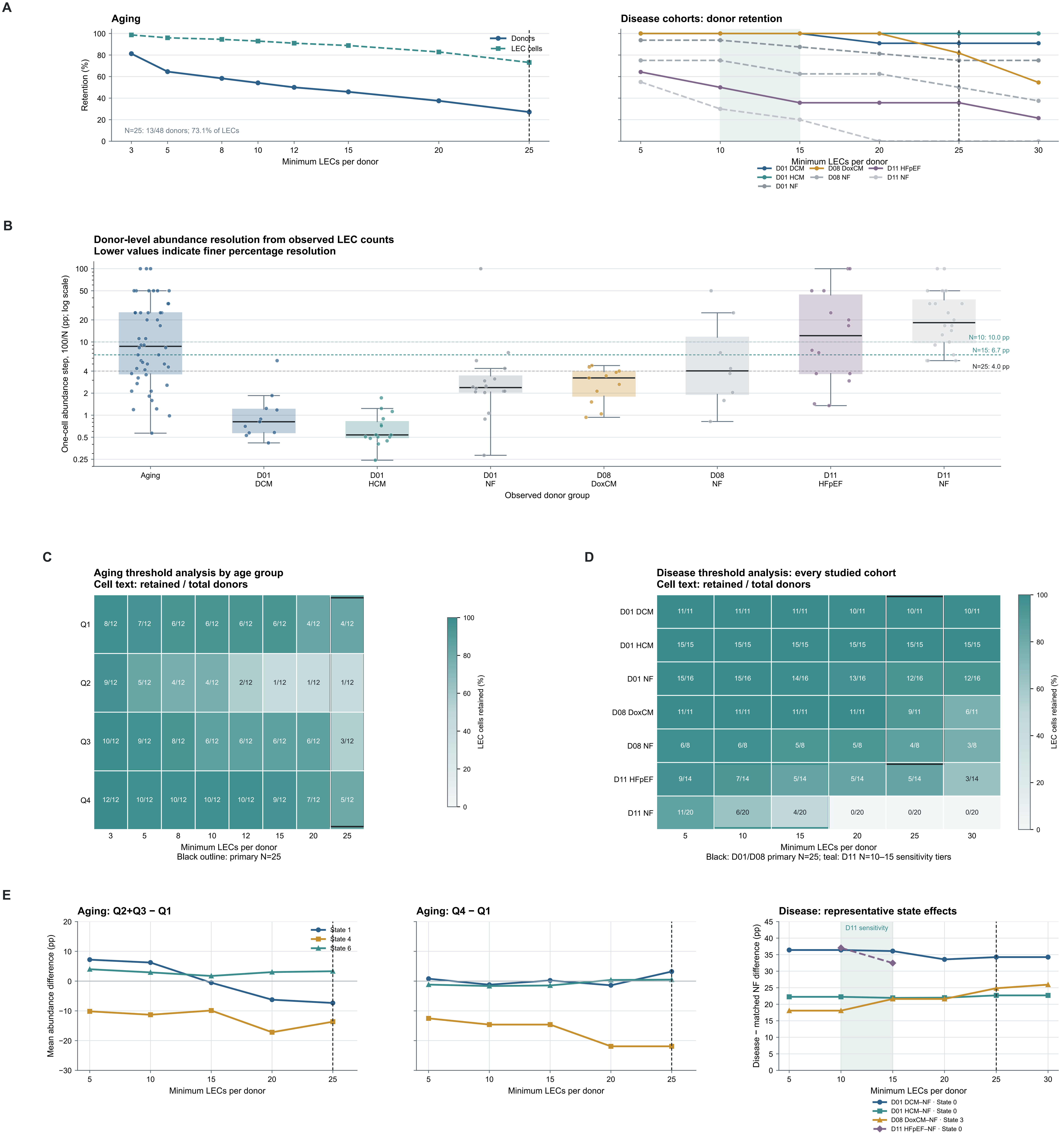

#### Figure S3 | Extended aging LEC-state characterization and robustness

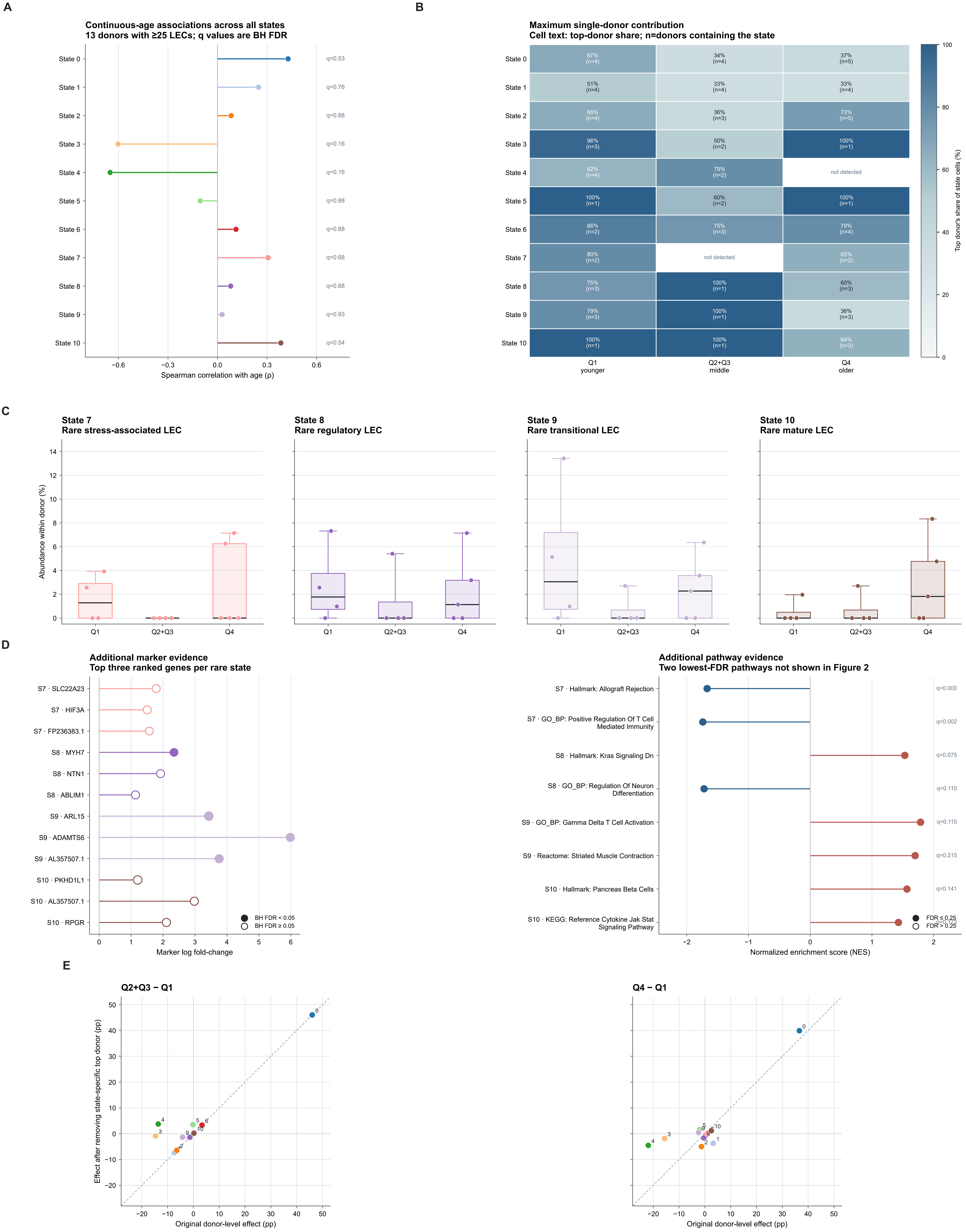

#### Figure S4 | Extended aging SCENIC and MultiNicheNet analyses

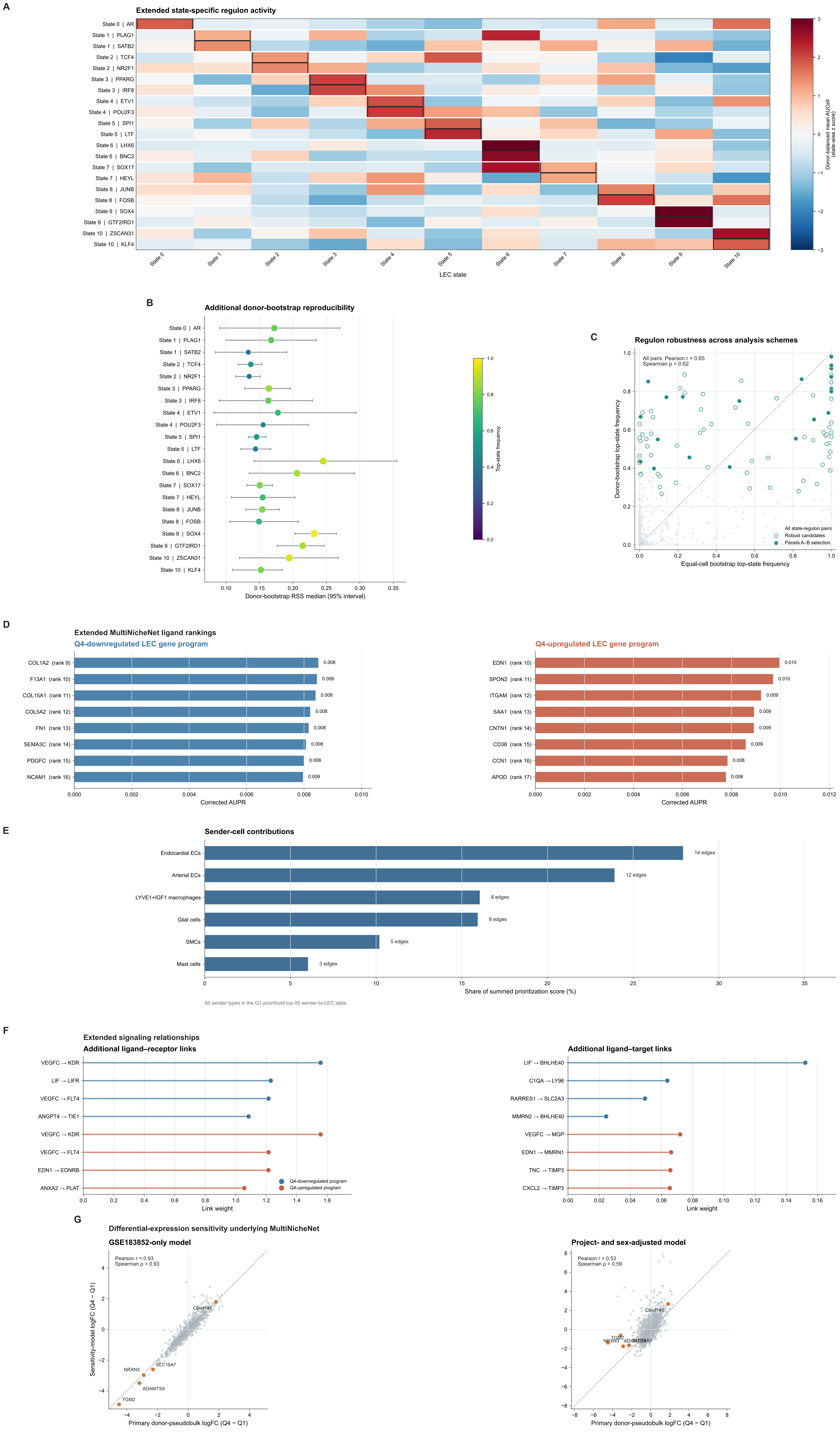

#### Figure S5 | Extended disease LEC-state characterization and robustness

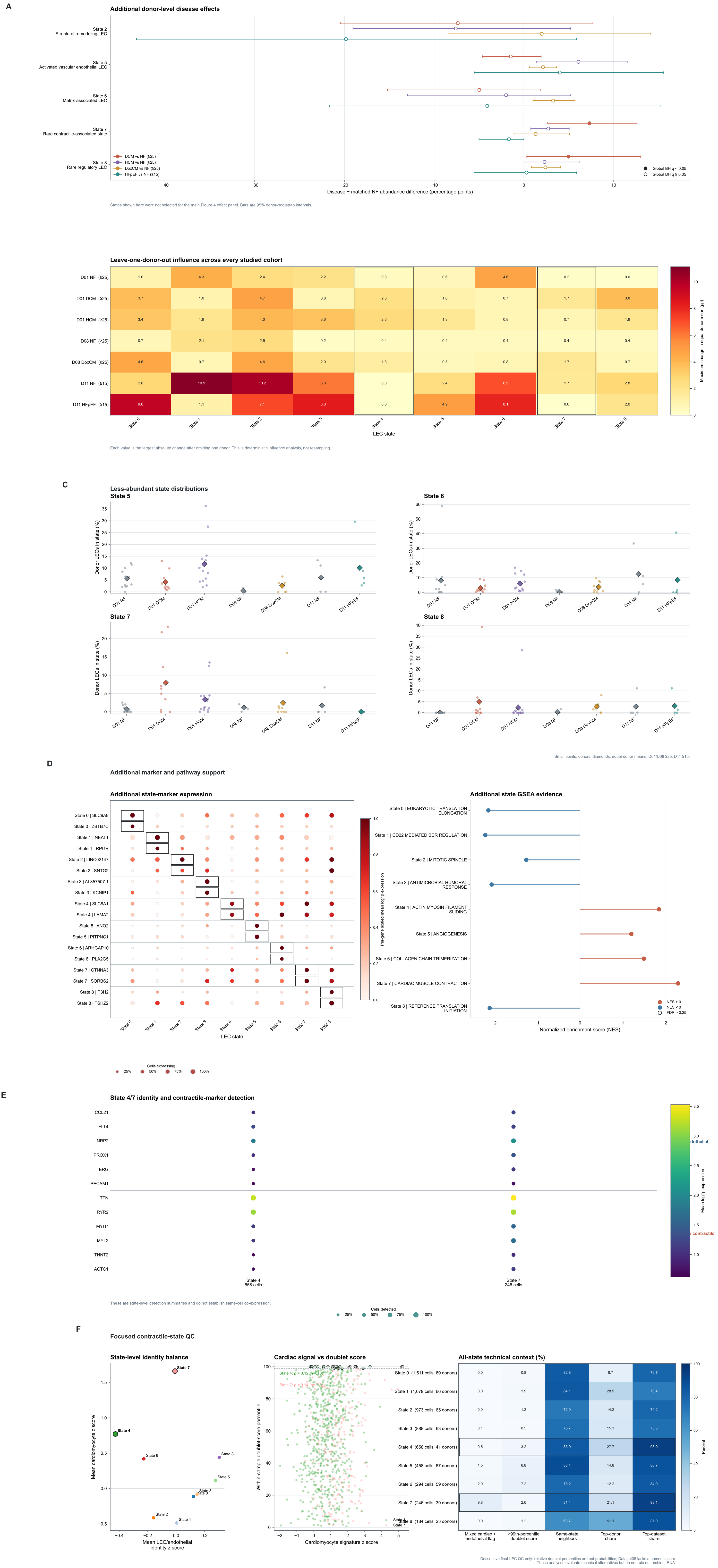

#### A Extended disease-state regulons

Two additional consensus-ranked robust candidates per state; main-figure pairs excluded

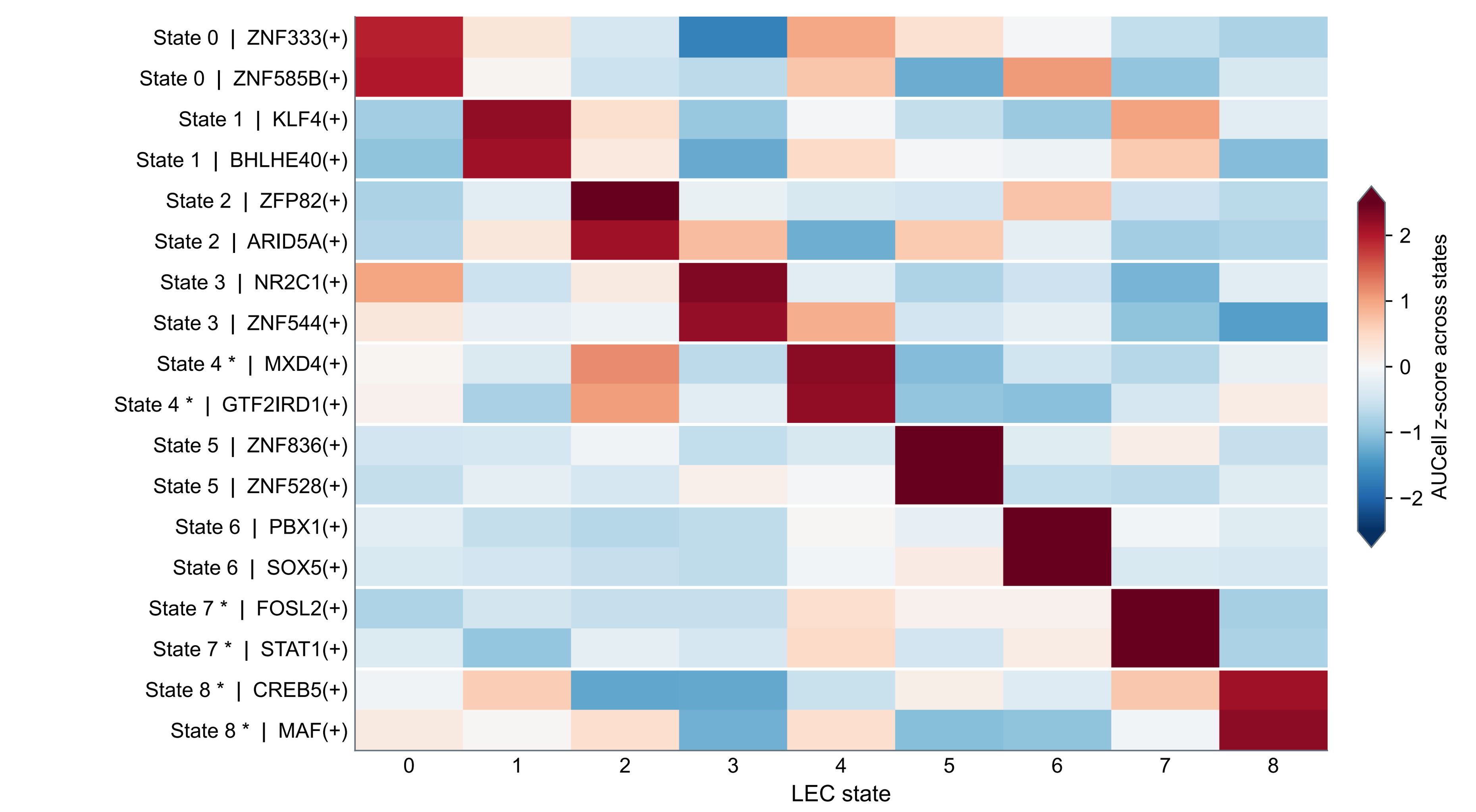


Saved strict dataset/donor-balanced activity: datasets 01 and 08.
* States 4/7: contractile-marker caution; State 8: donor-dominance caution.
Z-scores describe relative regulon activity, not disease-condition effects.

#### B Regulon reproducibility across balancing schemes

Existing 500-replicate bootstrap summaries for the additional regulons in A

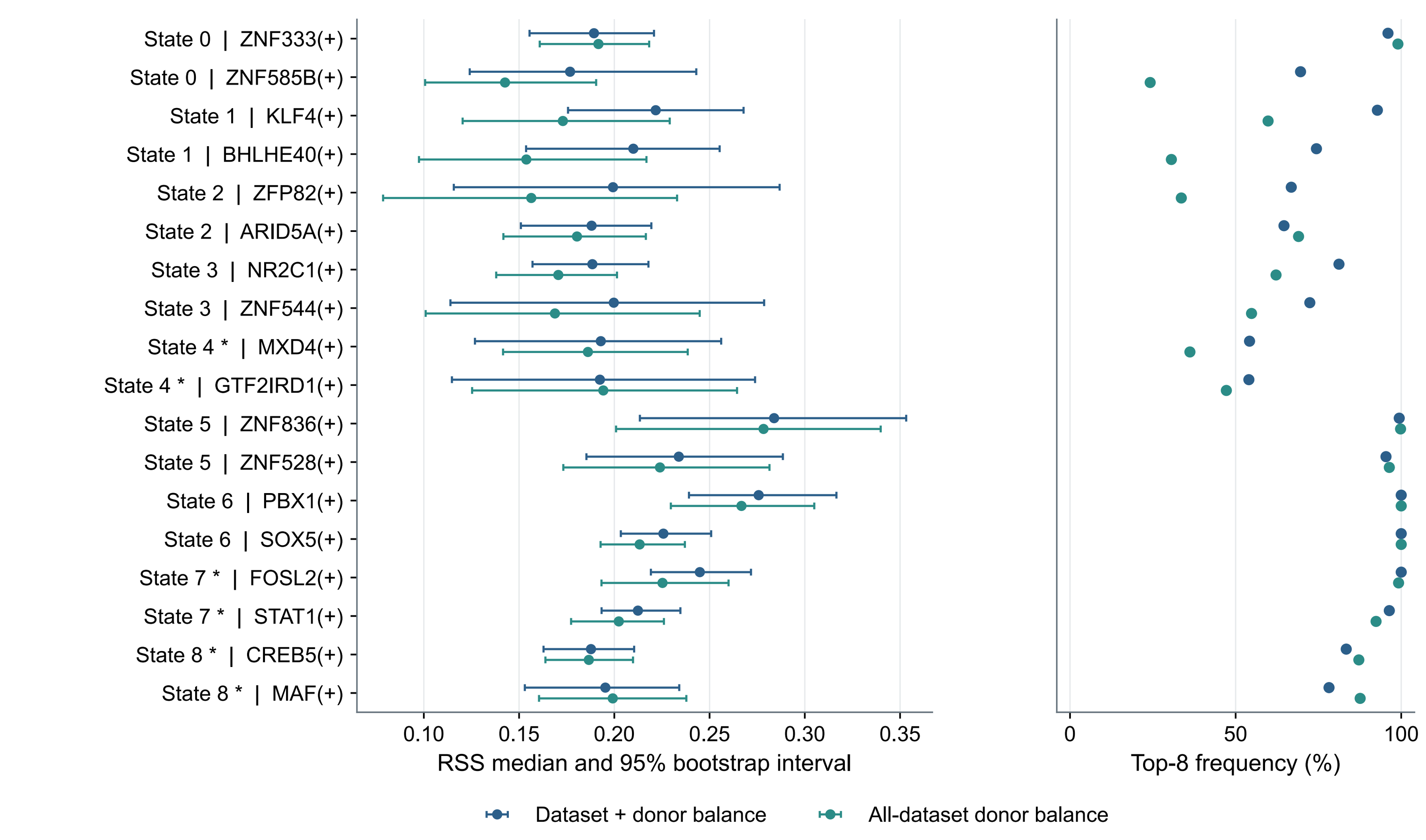


Strict balance: datasets 01/08; all-dataset sensitivity also includes dataset 11.
Top-8 frequency: fraction of saved replicates ranking the regulon among the top 8 for its state.
* State cautions as in A. Intervals are not p-values; no new resampling was run.

#### C Extended ligand activity rankings

Top five additional ligands per direction, ranked by corrected AUPR

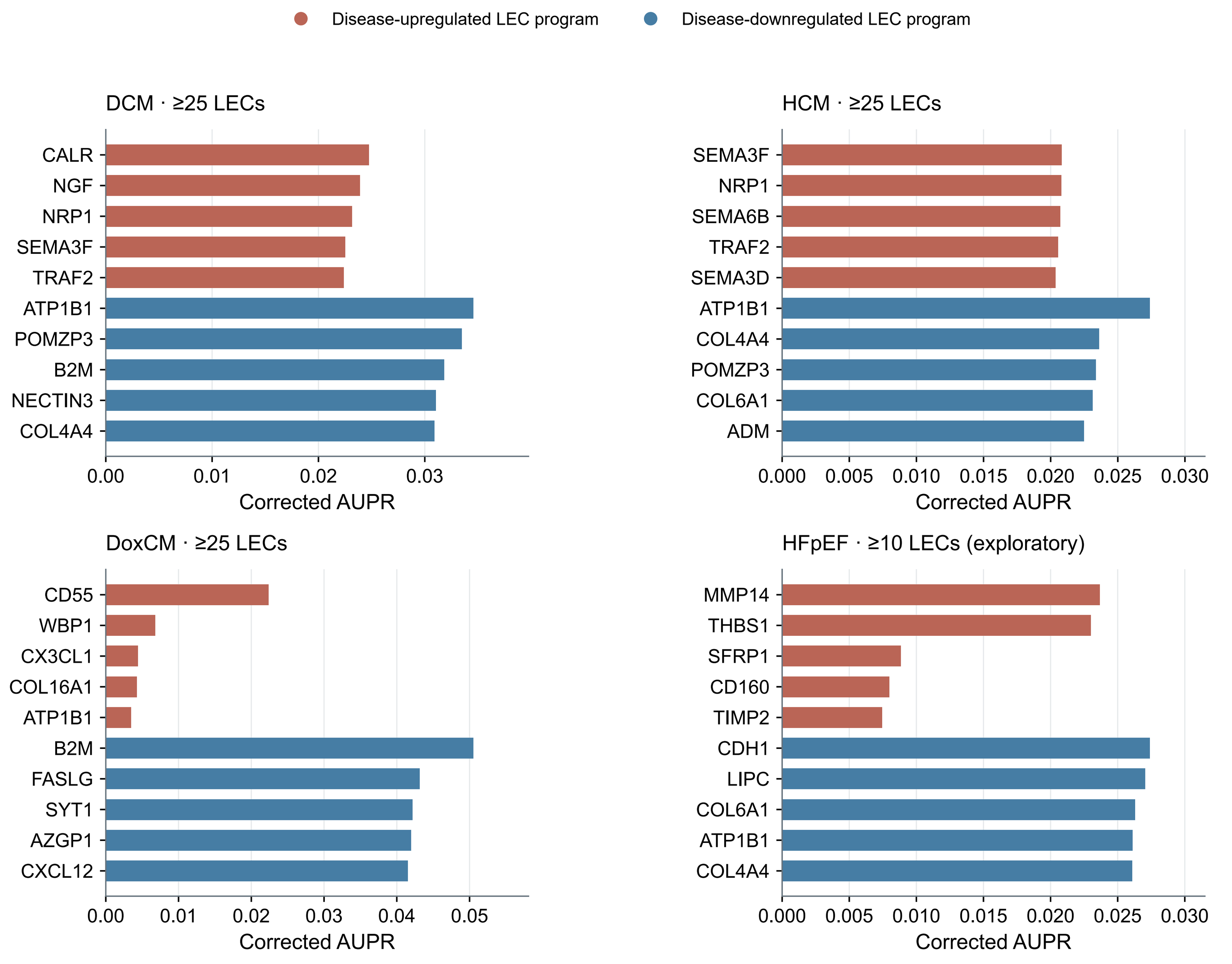


DCM, HCM, DoxCM: ≥25 LECs/donor; HFpEF: exploratory ≥10. Matched NF uses the same cutoff.
Direction refers to the receiver gene program, not ligand up/downregulation.
Activity is model-based ranking evidence; complete saved rankings are in the panel source CSV.

#### D Sender representation beyond selected axes

All sender types eligible in at least one disease; summary of the top 50 prioritized axes

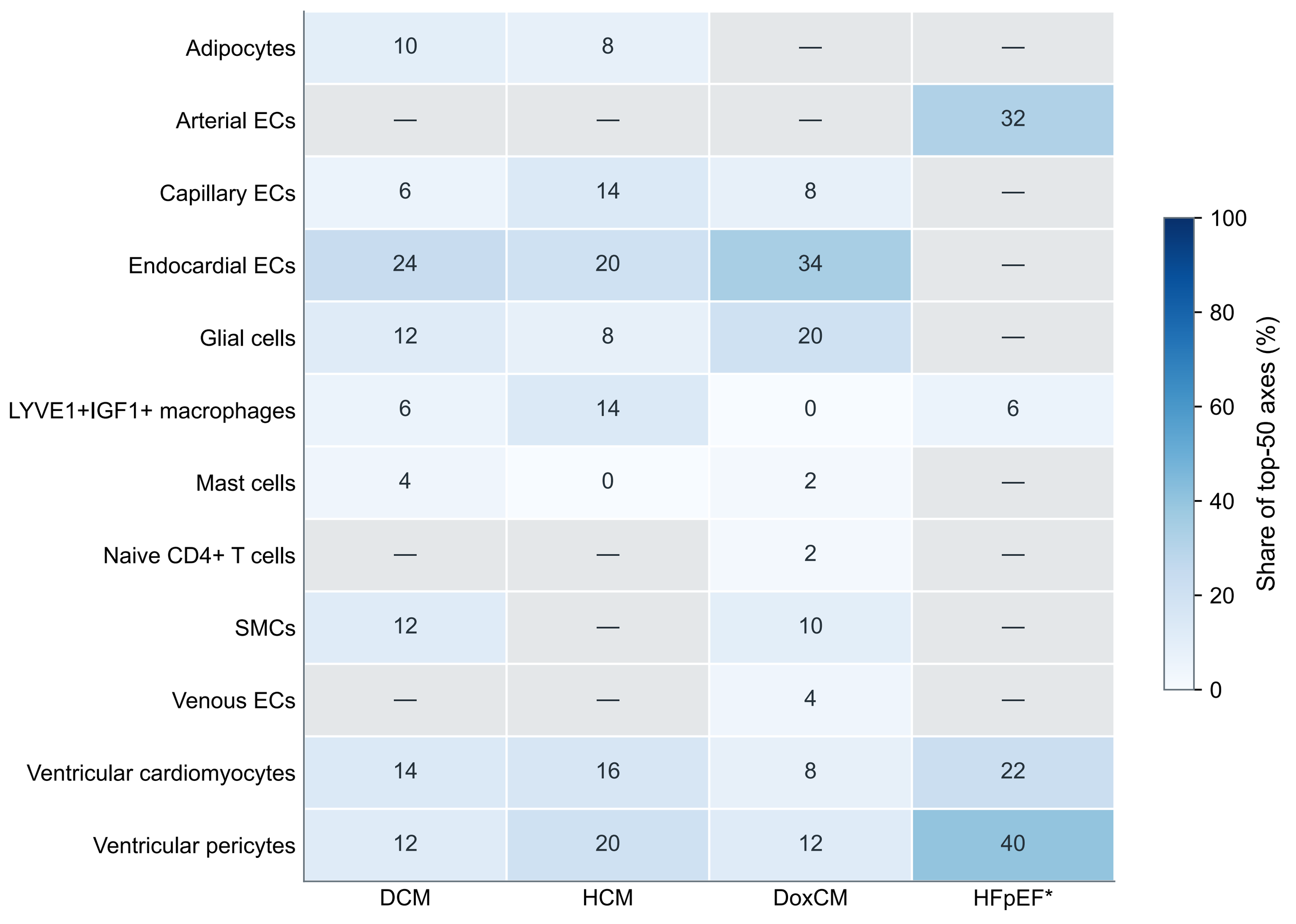


Each column sums to 100%. Share counts sender–ligand–receptor axes, not cells or signaling strength.
Gray/—: sender not eligible or not tested; 0: eligible but no axis among the top 50.
* HFpEF ≥10 LECs/donor; other diseases ≥25. Selection uses disease-prioritized axes only.

#### E Additional receptor and target links

Two strongest additional saved links per direction and type, for each disease

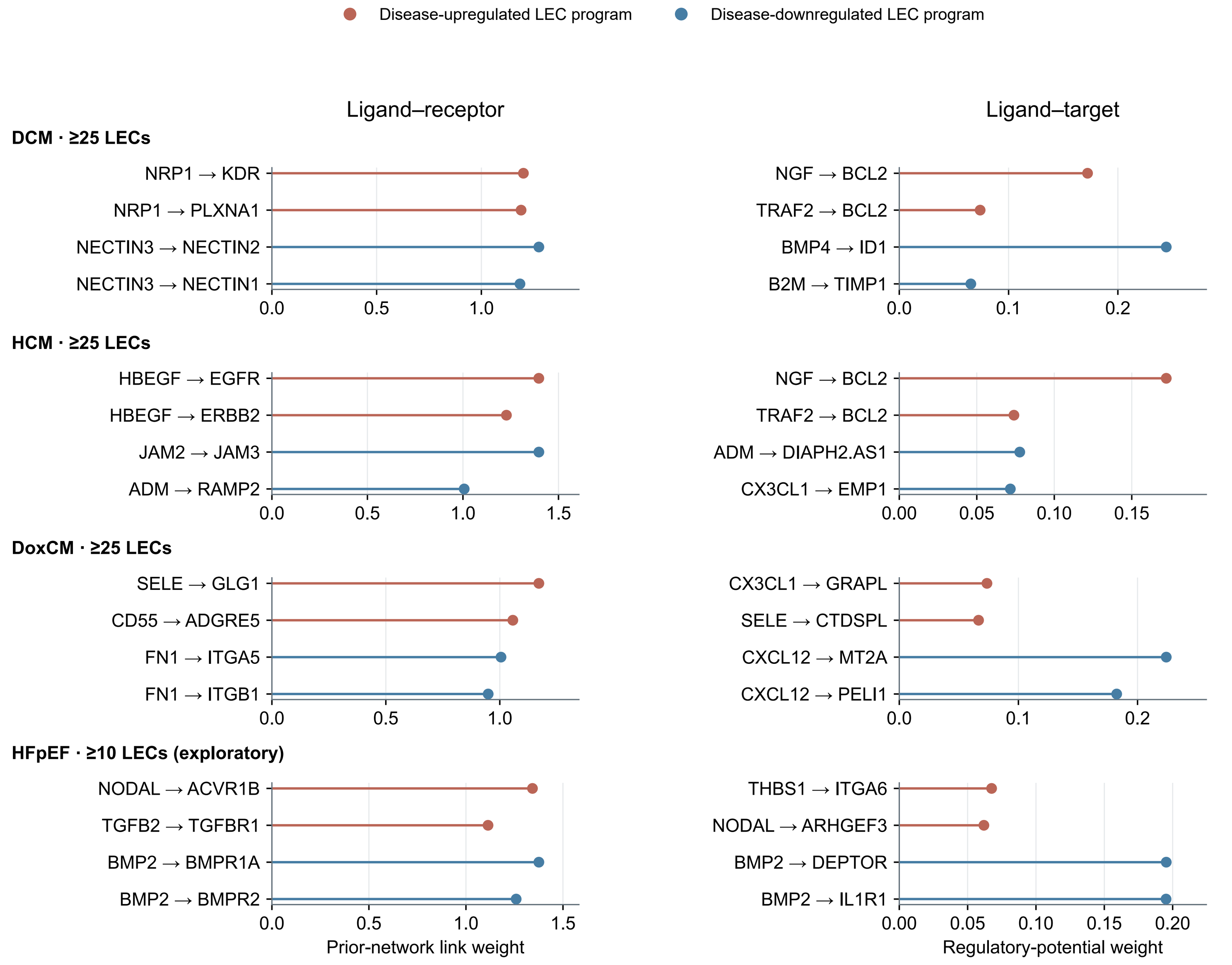


Receptor links already selected in Fig. 5 are excluded. These are extended saved candidate-link tables.
Weights are model/prior-network evidence, not measured interaction strength or validation.
Direction refers to the LEC gene program; weight scales differ between receptor and target links.
DCM, HCM, DoxCM: ≥25 LECs/donor; HFpEF: exploratory ≥10. Matched NF uses the same cutoff.

#### F Cross-disease context and HFpEF sensitivity

Additional ligand context across all diseases, with a saved cutoff sensitivity comparison

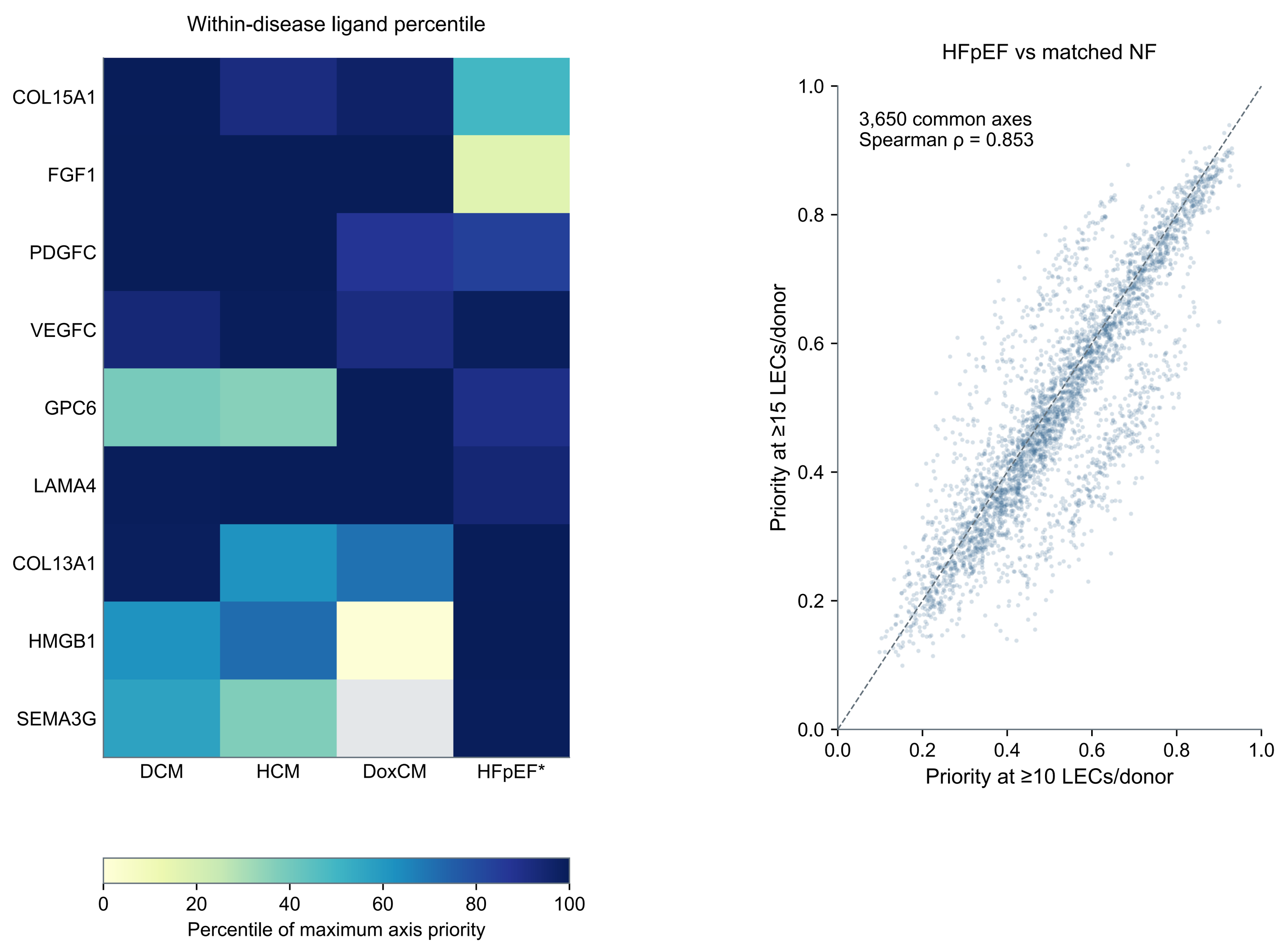


Left: union of the top 3 additional ligands per disease; all main-figure ligand selections excluded at selection.
Ranks are within-disease, not comparable effect sizes. Gray: no disease-prioritized qualifying axis.
Right: all common HFpEF-contrast axes, not only top-ranked axes; shared donors mean this is not replication.
* HFpEF exploratory ≥10, sensitivity ≥15; matched NF uses the same cutoff. Other diseases ≥25.

**A** Section-level Visium quality control

Eight sections from four donors; section-relative QC, not single-cell filtering

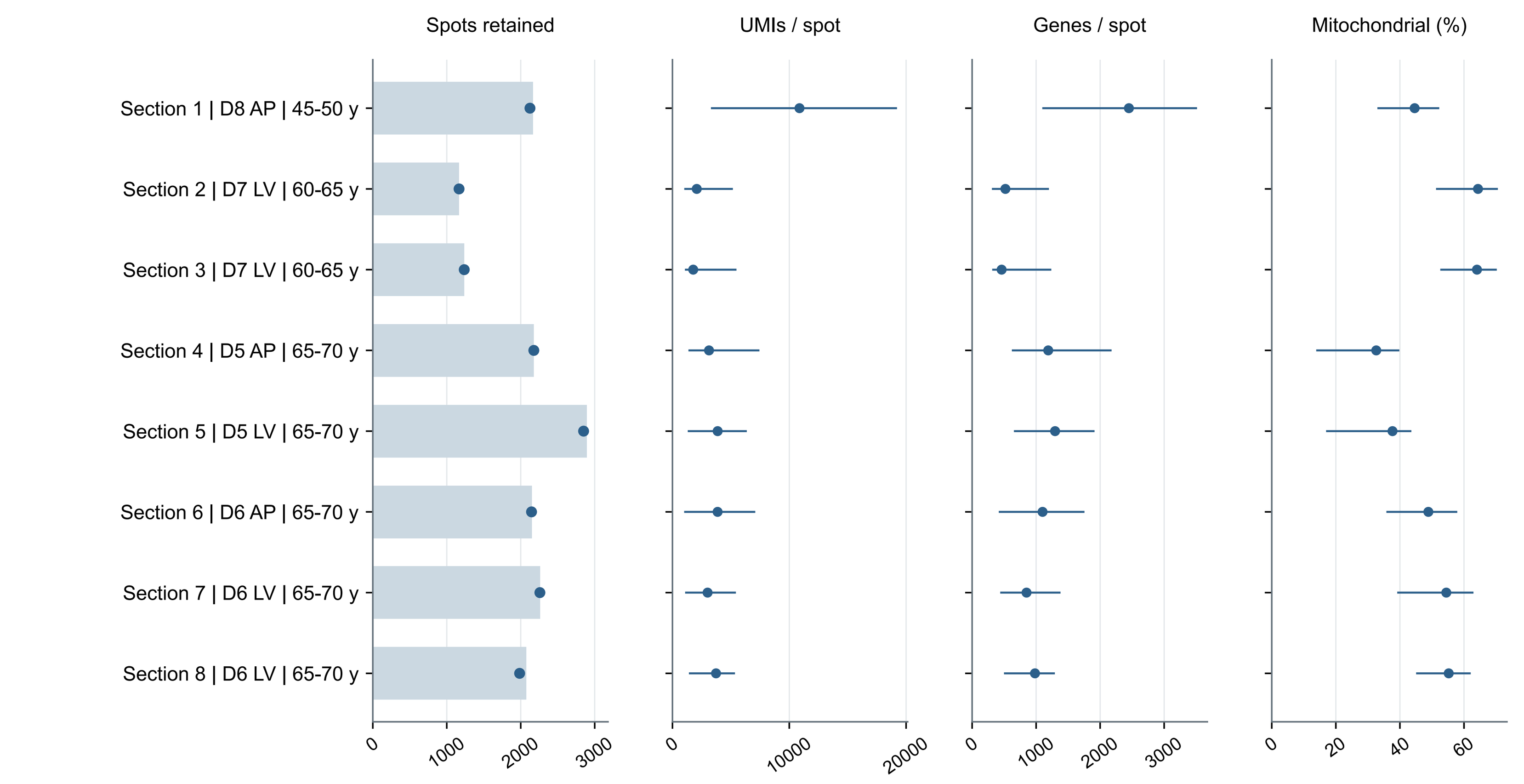


Bars: primary-QC spots; dots in first column: stricter technical-QC retention. Other dots: median; lines: 5th–95th percentiles.
Mitochondrial fraction alone is not an exclusion. AX is standardized to AP; section IDs are mapped in the source CSV.
D8 (45–50 years) is the only younger donor; repeated sections do not provide independent donor replication.

**B** LEC-enriched mixed-spot identification

Primary-QC spots with ≥2 of CCL21, LYVE1, PDPN, FLT4 and PROX1 detected

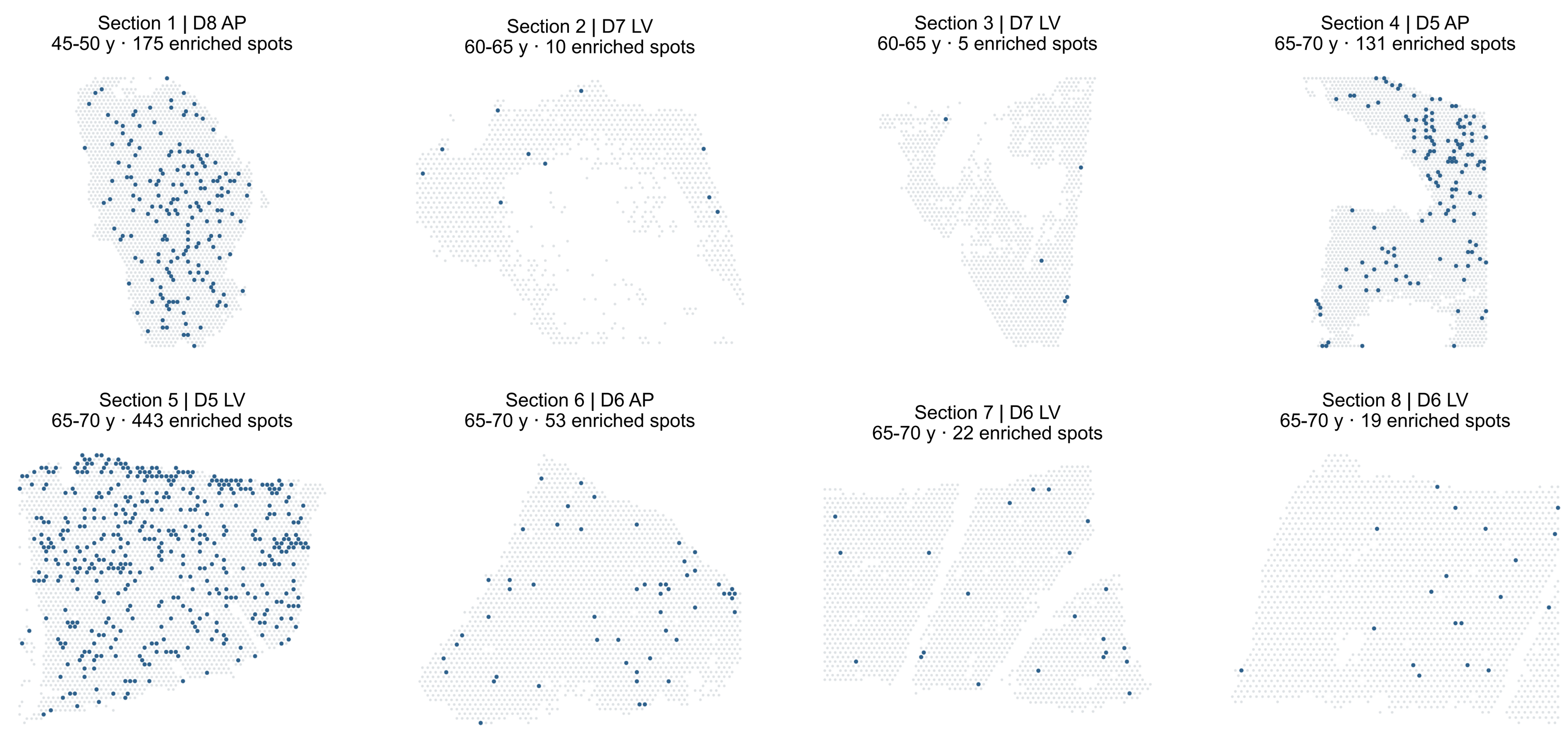


Blue: LEC-enriched; gray: other QC-passing spots. Detection means raw count >0; section coordinate scales are independent.
Cell2location EC8 estimates are positive throughout the resource, so positivity is not an LEC-selection rule.
These are mixed-cell Visium spots, not purified LECs or categorical LEC-state assignments.

**C** Canonical LEC-marker expression and coverage

Dot area: marker detection; color: mean log1p(CP10K) across the indicated spots

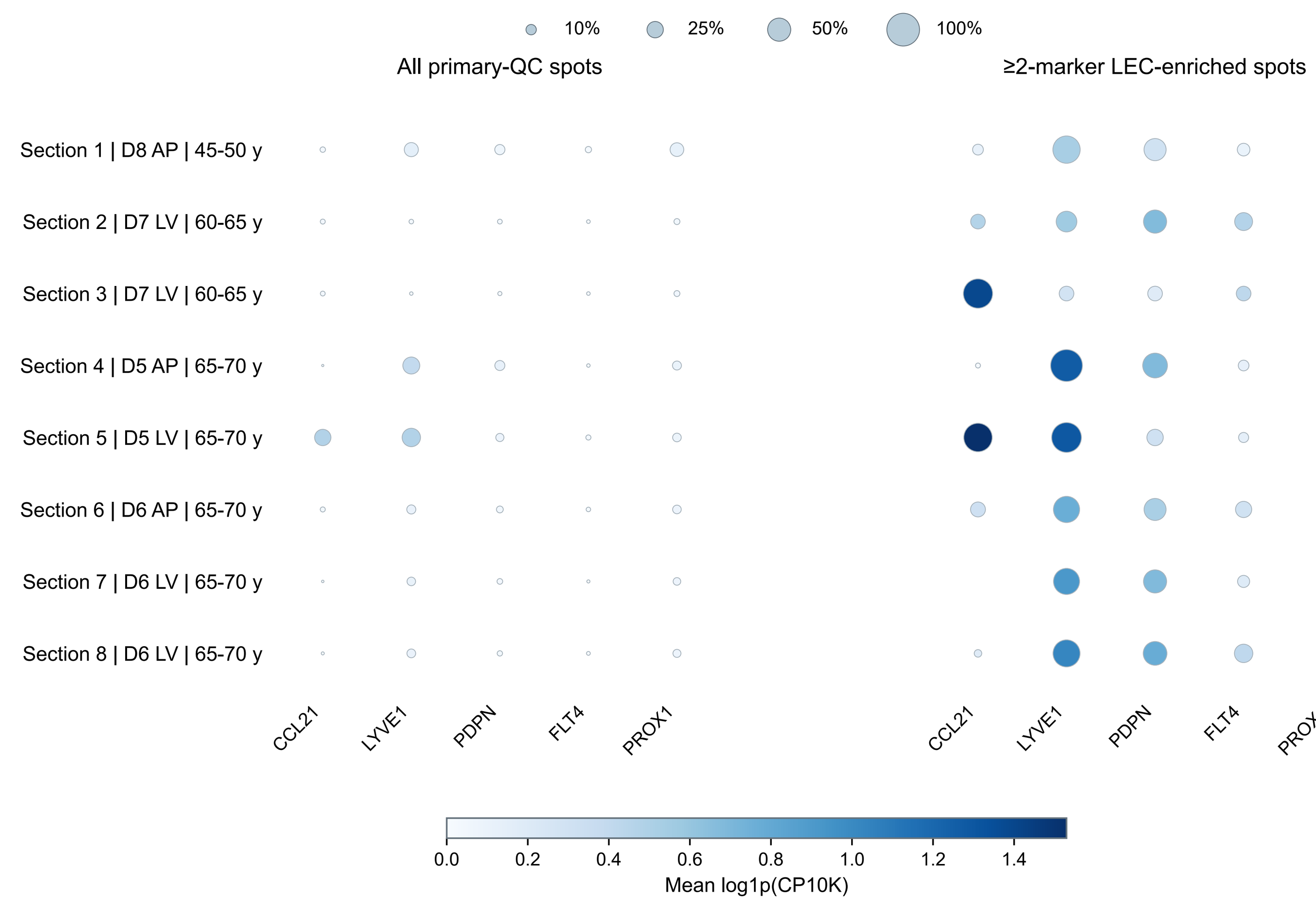


Detection denominator: all spots in the stated section/gate, including zero-expression spots. Blank dots denote zero detection.
Expression is the mean of saved log1p(CP10K), including zeros—not log1p of the mean.
Higher marker coverage after gating is expected by construction and is not independent validation.

**D** Sensitivity to LEC-enrichment criteria

Compare the existing ≥2-marker gate with ≥3 markers and stricter technical QC

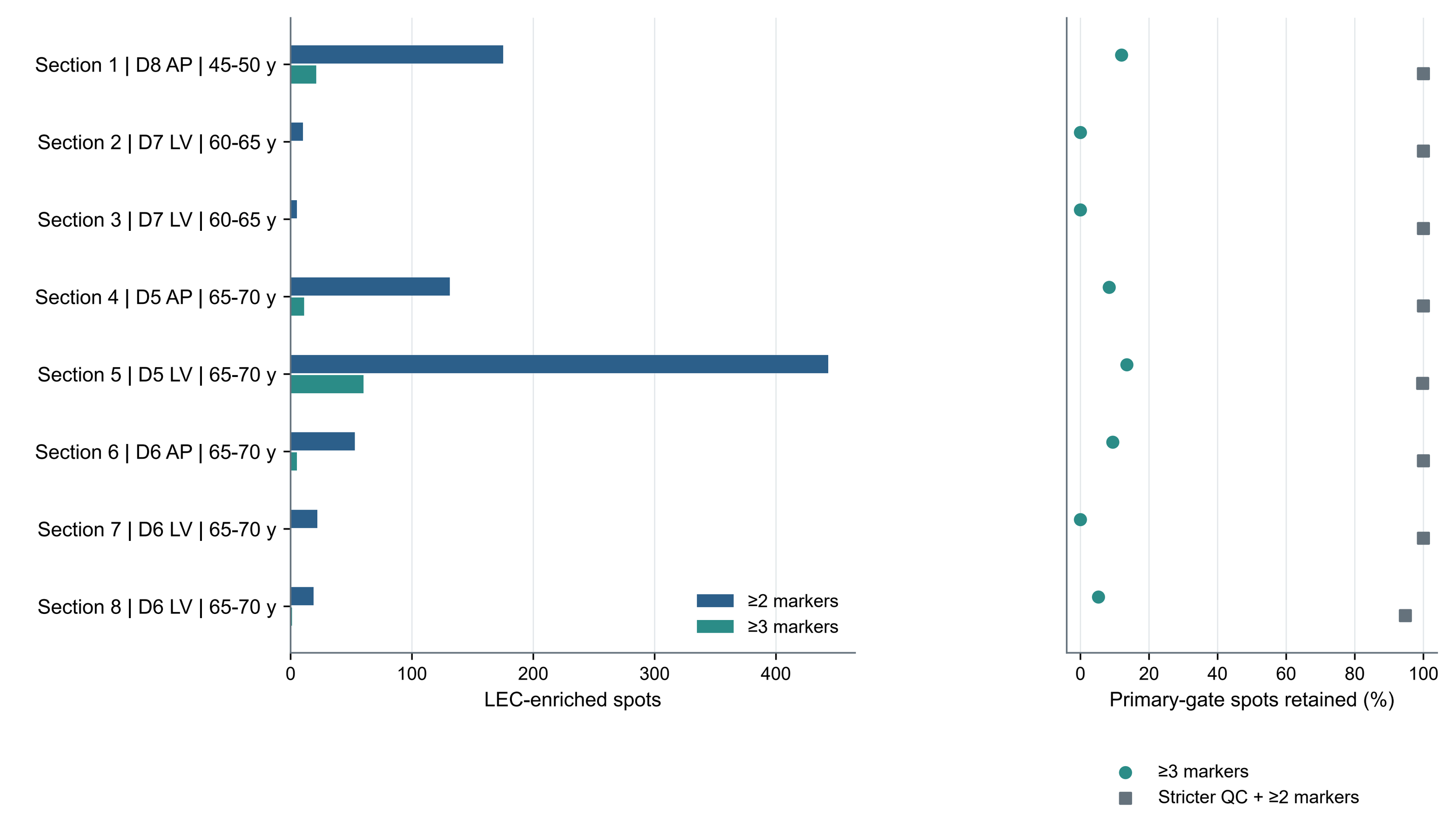


≥2 markers: 858 spots; ≥3 markers: 98 spots. All sections remain shown, including three with no ≥3-marker spots.
Stricter technical QC retains 856/858 primary-gate spots; this is a tabulation of saved flags, not a score/model refit.
A one-marker enrichment analysis was not available and is not introduced here.

**E** Program-score and spatial-sampling sensitivity

All saved gate/model settings and within-section spatial-block uncertainty

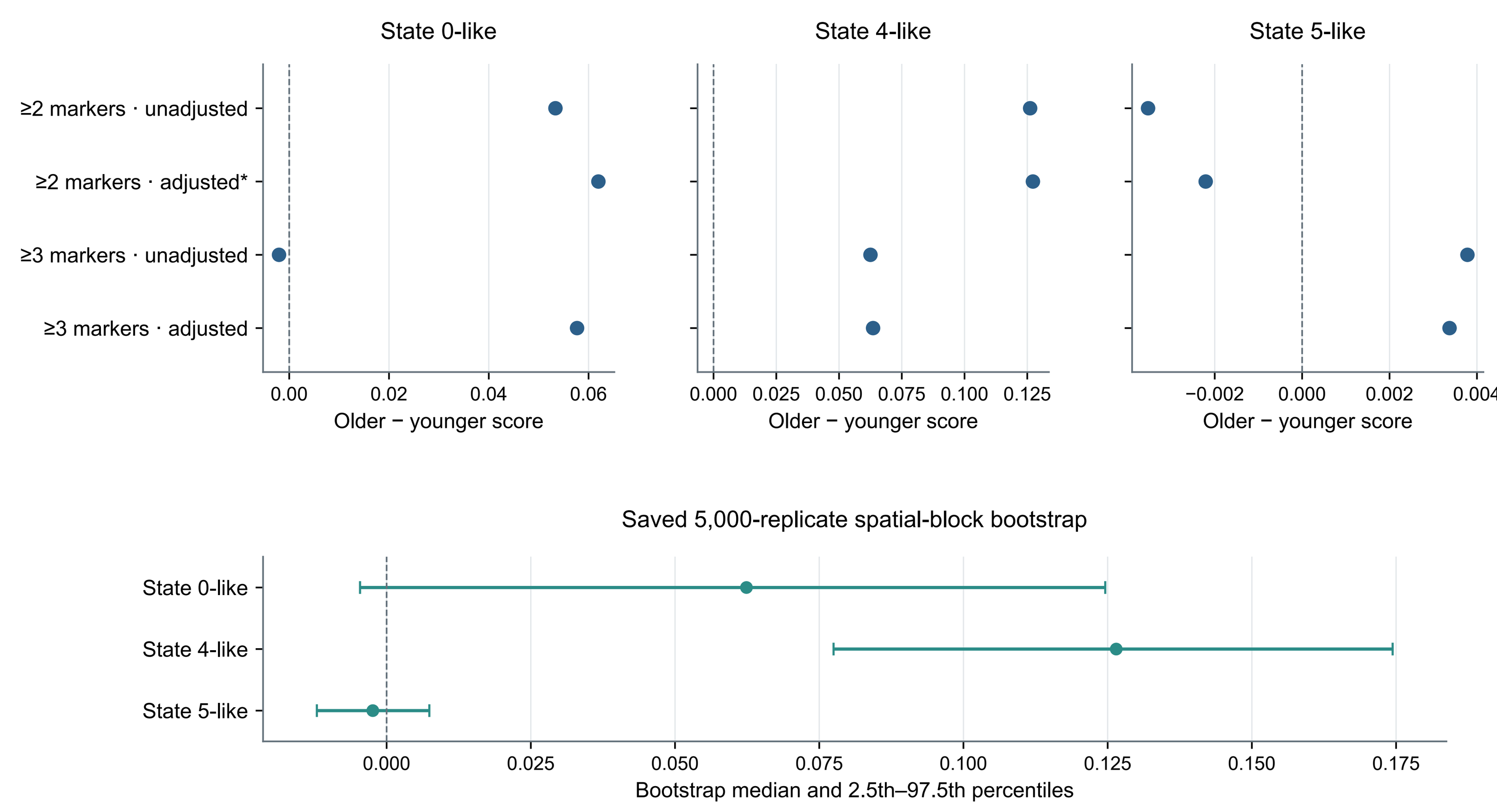


* Fixed primary endpoint: AP D8 versus the equal donor mean of D5/D6, ≥2 markers, mixture-adjusted score.
Unadjusted denotes the matched-control score before competitor-mixture adjustment; these are not raw counts.
Bootstrap intervals describe within-section sampling only, not donor uncertainty or an age test. No new resampling was run.

**F** Leave-one-gene-out program sensitivity

All nine saved gene-omission analyses; program-gene coverage remains visible

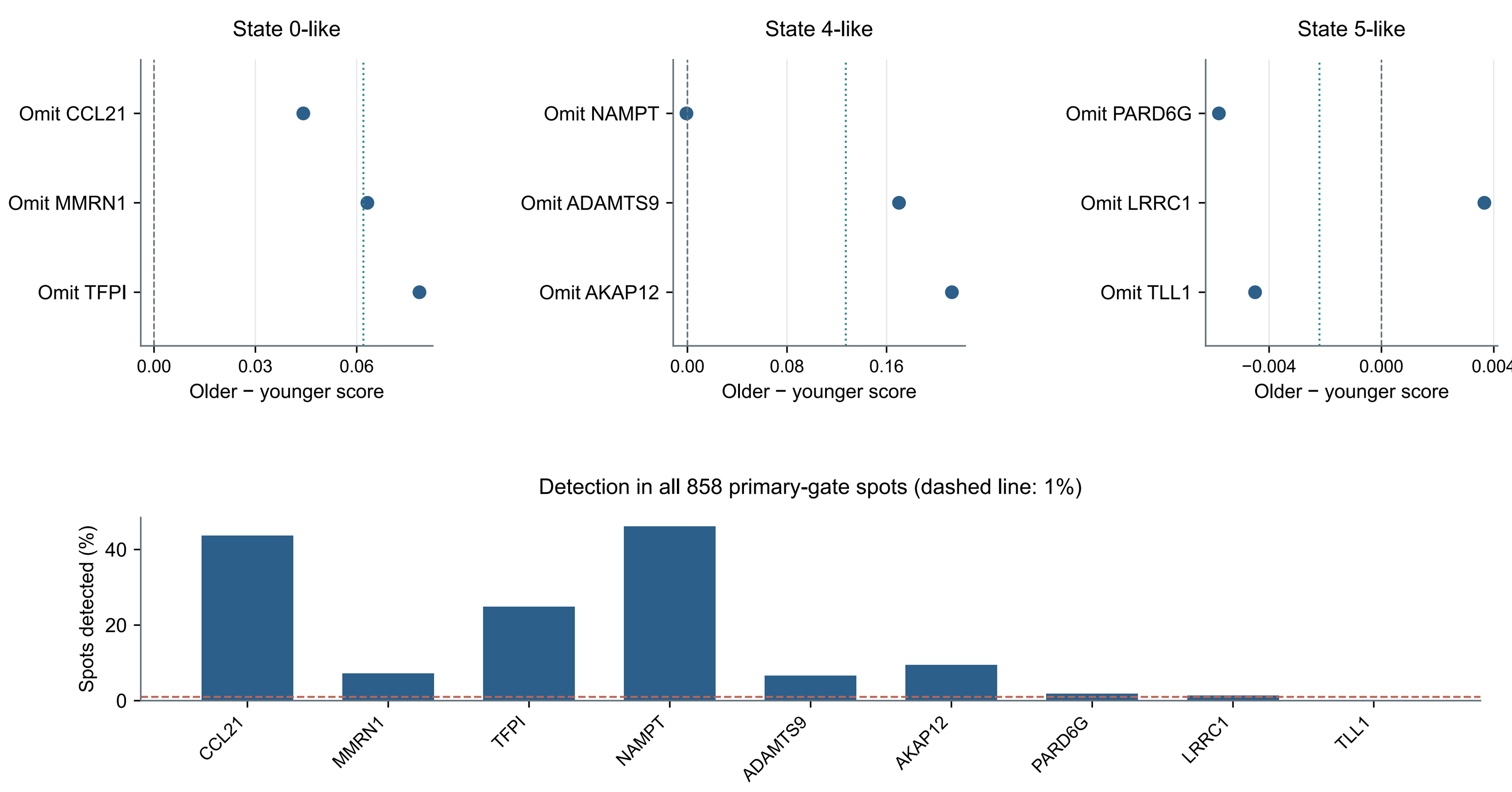


Dotted teal lines: fixed full-panel effect. Each omitted-gene score was previously recomputed and mixture-adjusted upstream.
CCL21 participates in both selection and the State 0-like score; omitting it from the score does not remove it from selection.
TLL1 is sparsely detected. The saved robustness rule requires ≥2 of 3 gene-omission directions, not unanimity.

**G** Donor dependence and section context

All-section program context, older-donor omission and paired AP/LV sensitivity

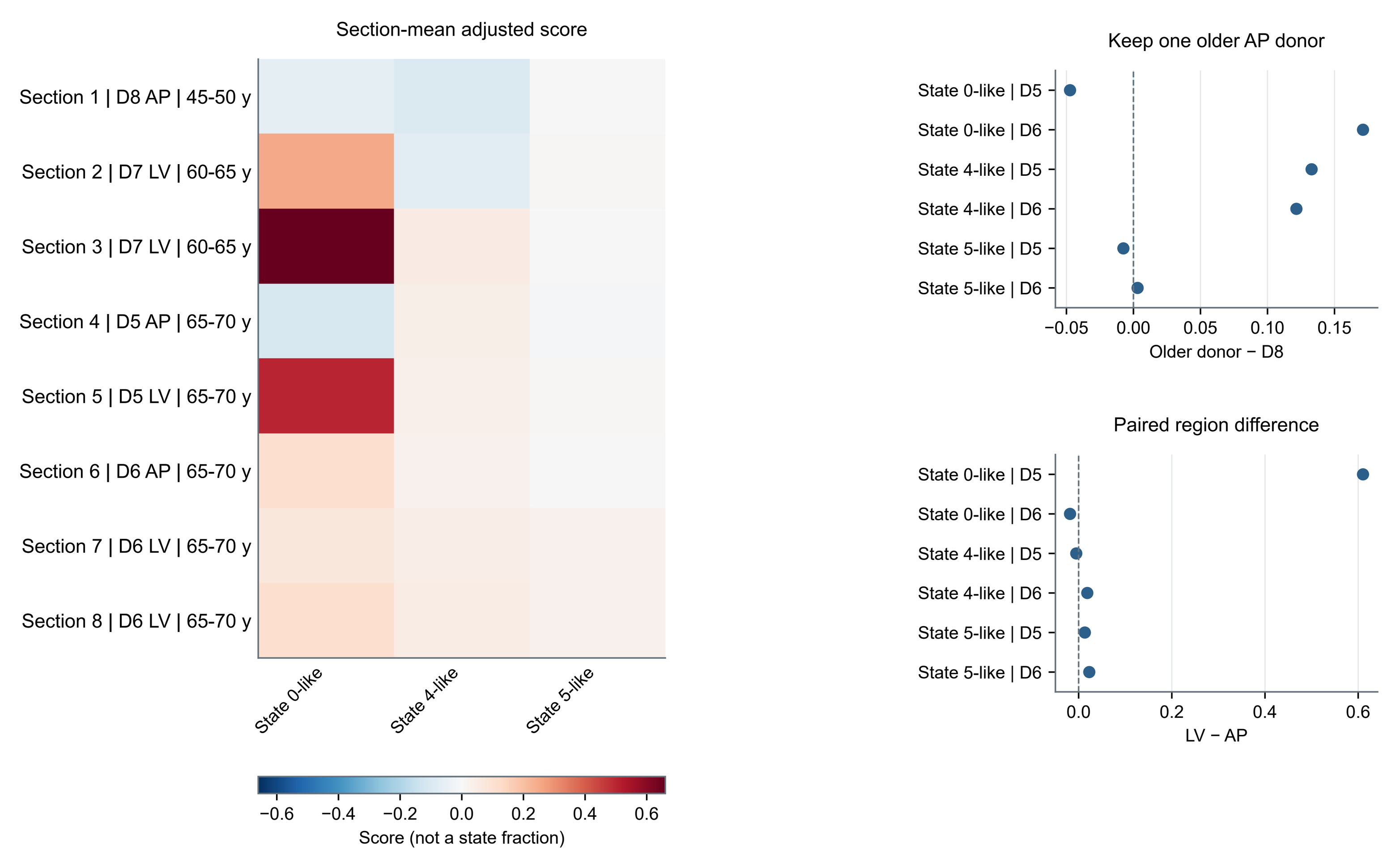


Section means are mixed-spot scores; sections with few enriched spots remain visible (counts in D).
AP/LV contrasts pair regions within D5 or D6; LV sections receive equal weight within each donor.
D7 contributes LV context only. No independent younger-donor omission check is possible with one younger donor.

**H** Detailed robustness and sensitivity matrix

Original configured checks and canonical conclusions; all three programs reported

| | State 0-like | State 4-like | State 5-like |
|---|---|---|---|
| Primary expected direction | Yes | No | Yes |
| Both adjusted enrichment gates | 2/2 | 0/2 | 1/2 |
| Both score methods (≥2-marker gate) | 2/2 | 0/2 | 2/2 |
| Both older AP donors separately | 1/2 | 0/2 | 1/2 |
| Gene omissions: ≥2/3 agree | 3/3 | 1/3 | 2/3 |
| Detectability: ≥2/3 genes at ≥1% | 3/3 | 3/3 | 2/3 |
| Block-bootstrap direction probability ≥0.80 | 0.966 | 0.000 | 0.678 |
| All configured grading rules met | No | No | No |
| Region sensitivity flag* | Flagged | Not flagged | Flagged |
| **Canonical outcome** | **Fragile / inconclusive** | **Not spatially corroborated** | **Fragile / inconclusive** |

Teal: configured rule met; pale coral: not met. These checks are not significance tests.
* Region flag is separate context, not an additional criterion in the original composite grading rule.
None of the three programs qualifies as positive spatial validation; one younger donor limits all age interpretation.

#### A Overall GeoMx dataset and AOI quality control

GSE271676: 178 segmented AOIs from 44 donors; all passed the saved AOI-level QC gate

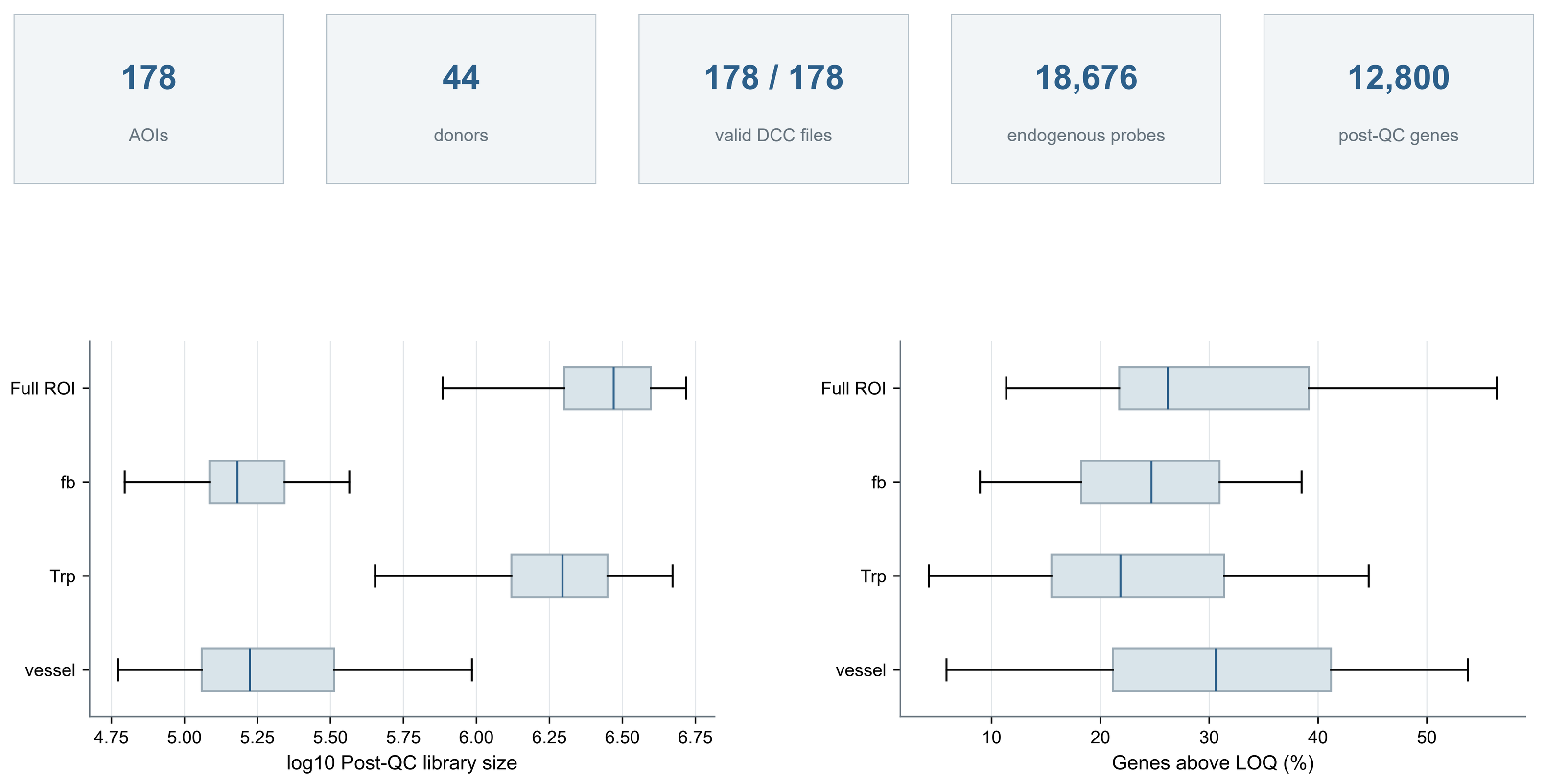


AOIs are segmented regions, not cells. Vessel AOIs are CD31-enriched endothelial regions, not purified LECs.
QC used raw DCC validation, negative-probe background, post-probe-QC library size and above-LOQ gene coverage.
No AOI hard failures occurred; repeated AOIs within a donor are not independent biological replicates.

#### B Rationale for selecting LV vessel AOIs

Anatomical, segmentation and phenotype restrictions define the 16-AOI primary pathway cohort

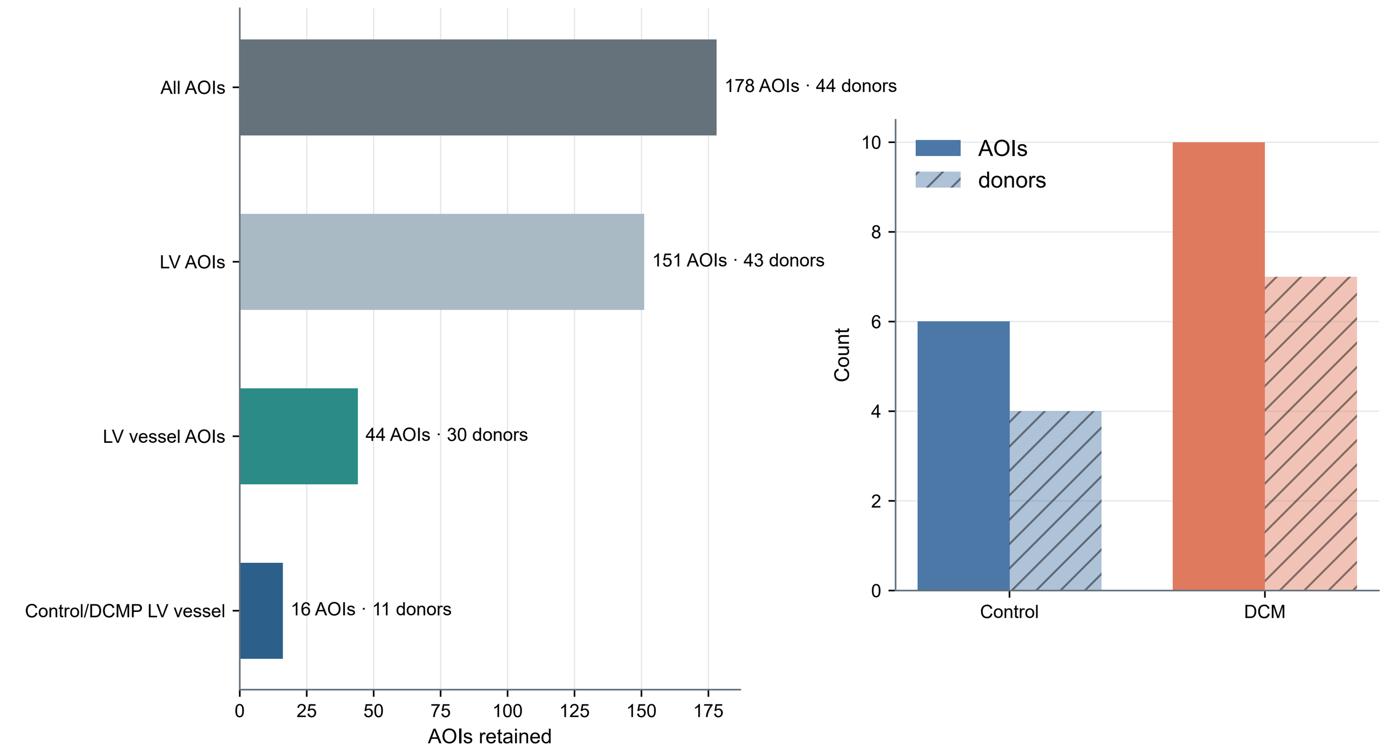


Primary screen: Ventricle=LV, vessel segment, Clinical_phenotype_LV in Control/DCMP.
This matches cardiac anatomy, endothelial segmentation and the DCM contrast; it is not an expression-selected cohort.
Six Control AOIs from four donors and ten DCM AOIs from seven donors permit donor-level testing in the pathway screen.

#### C Criteria for identifying LEC-like AOIs

Prespecified rule: ≥2 of five canonical lymphatic markers above each AOI-specific LOQ

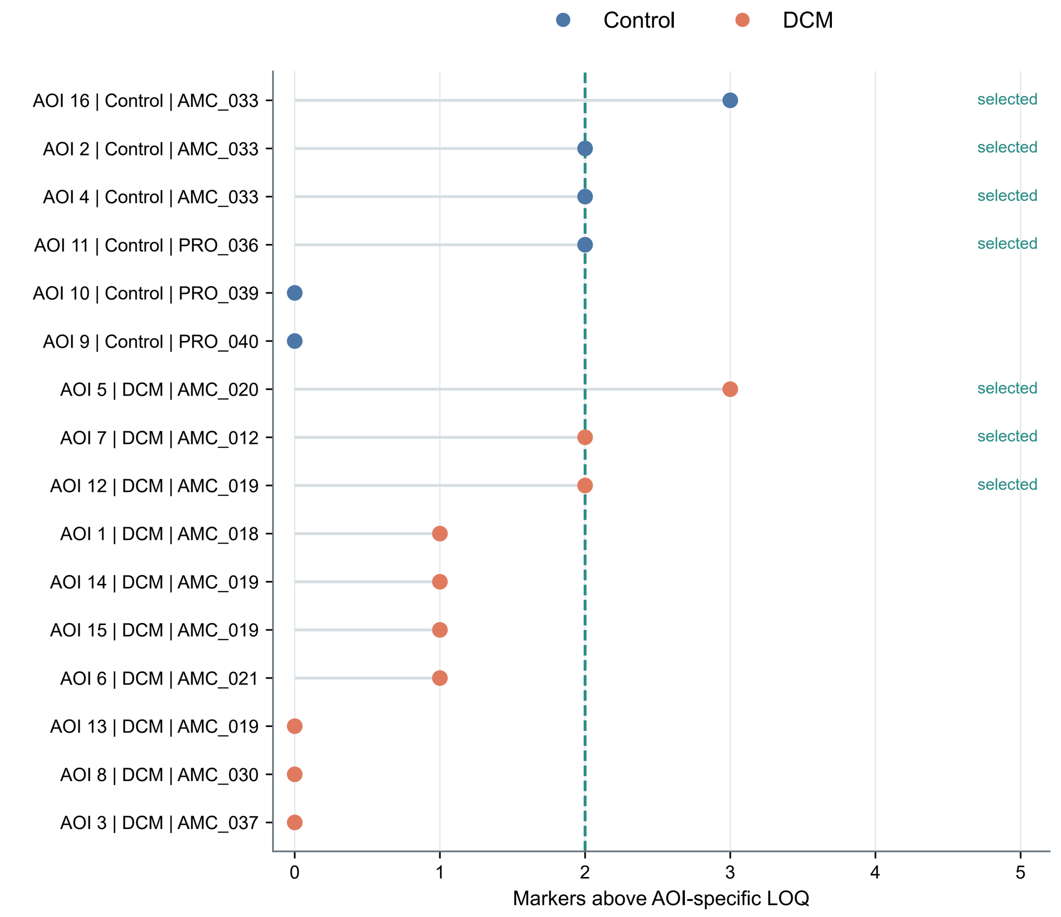


Markers: CCL21, LYVE1, PDPN, FLT4 and PROX1. Detection means target count above the AOI-specific GeoMx LOQ.
Seven of 16 AOIs were selected: four Control AOIs from two donors and three DCM AOIs from three donors.
Two markers in a mixed vessel AOI do not demonstrate co-expression in one cell; no formal selected-cohort test is permitted.

#### D Lymphatic-marker expression and coverage

Dot area: percentage above LOQ; color: mean log2 upper-quartile CPM + 1

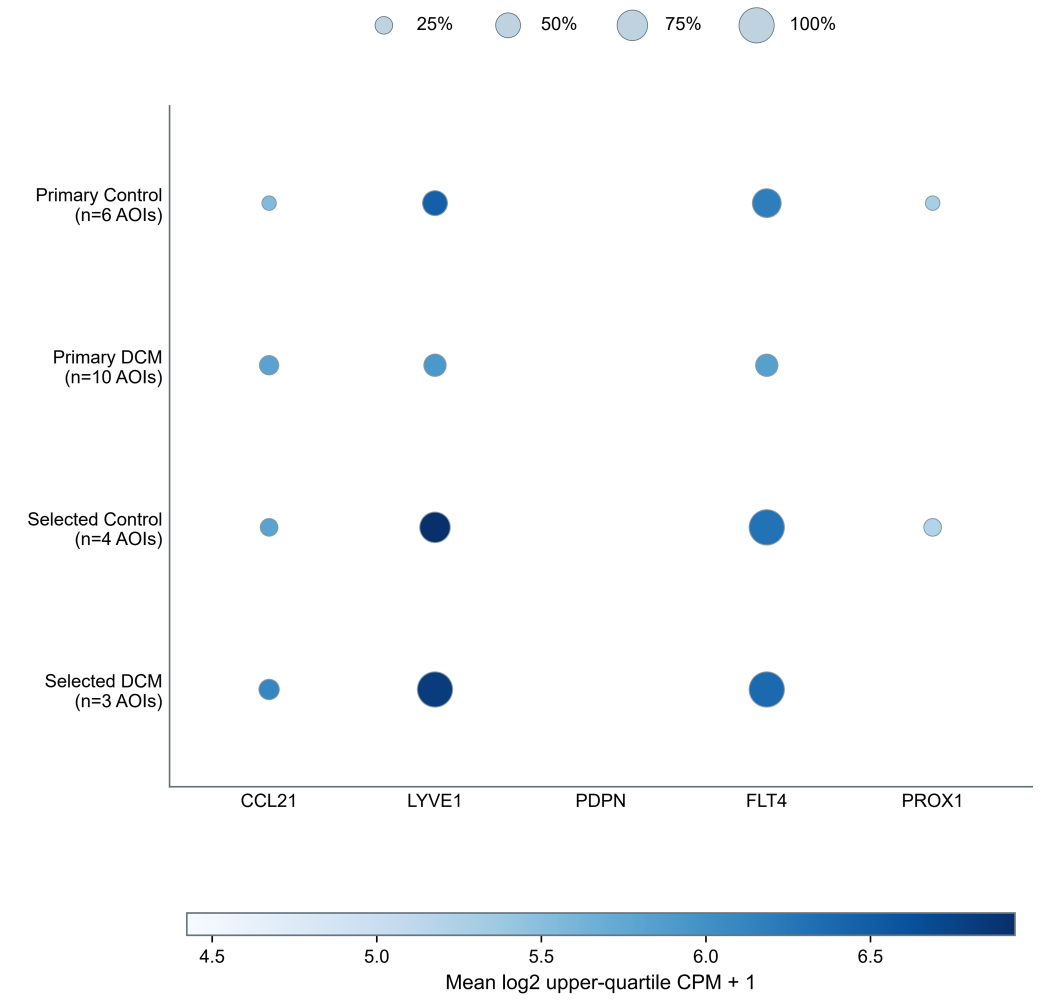


Detection uses raw target count > AOI-specific LOQ; expression is the mean normalized value including below-LOQ AOIs.
Selected-group marker enrichment is expected because these five markers define selection; it is not independent validation.
CCL21 also appears in the homeostatic endpoint, creating explicit selection–outcome overlap.

#### E Probe detection and quality assessment

Raw detection is shown before the authors' 1%-probe-QC expression filter

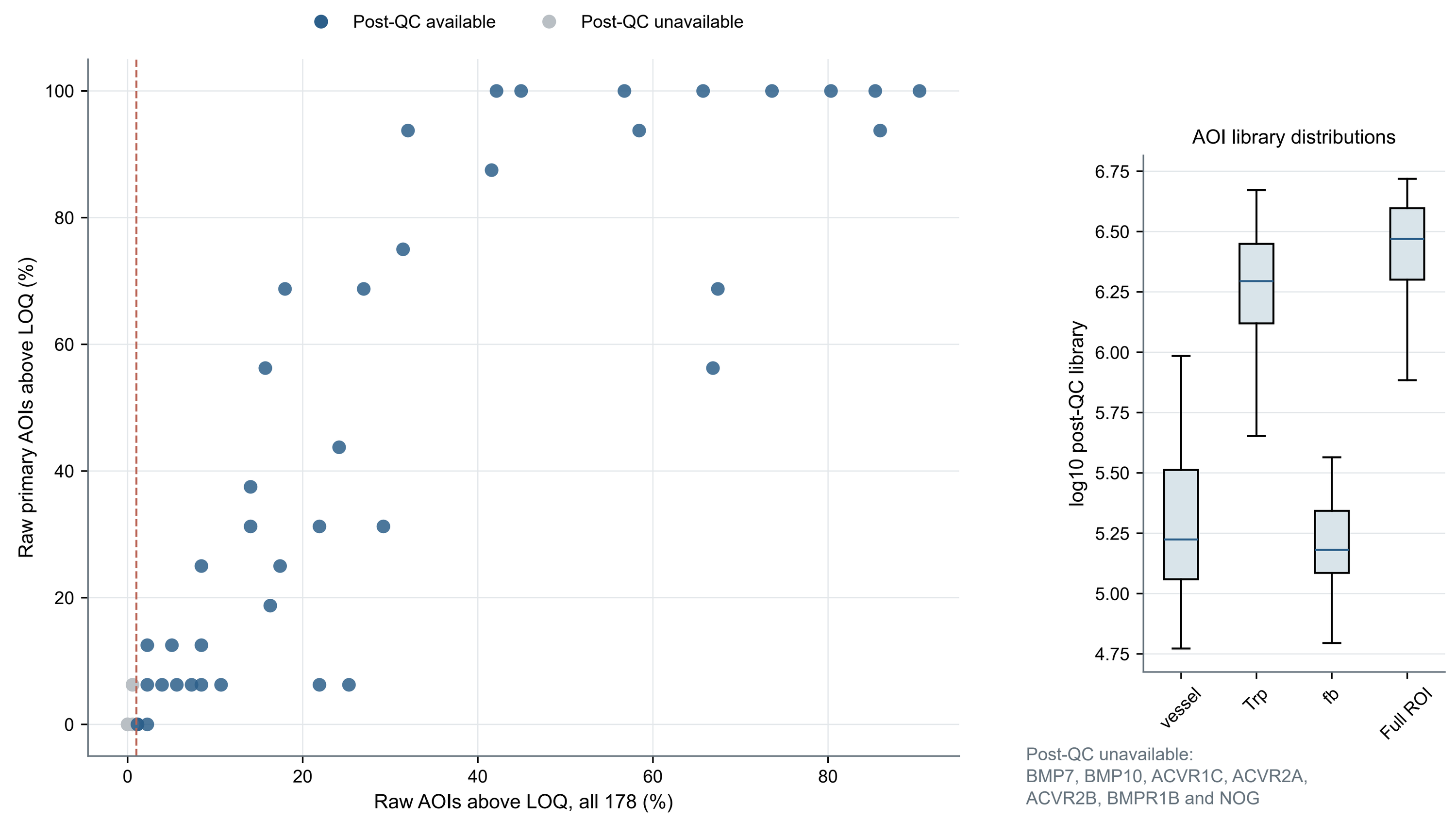


Each left point is one of 47 prespecified pathway genes. Gray points failed the authors' probe-QC expression matrix.
The dashed 1% line contextualizes sparse raw detection; probe retention is read from the supplied filtered matrix.
Unavailable genes remain in the audit but are not reintroduced for expression inference.

#### F BMP/TGF-β/activin pathway assay coverage

Complete 47-gene audit: raw above-LOQ coverage and post-probe-QC availability

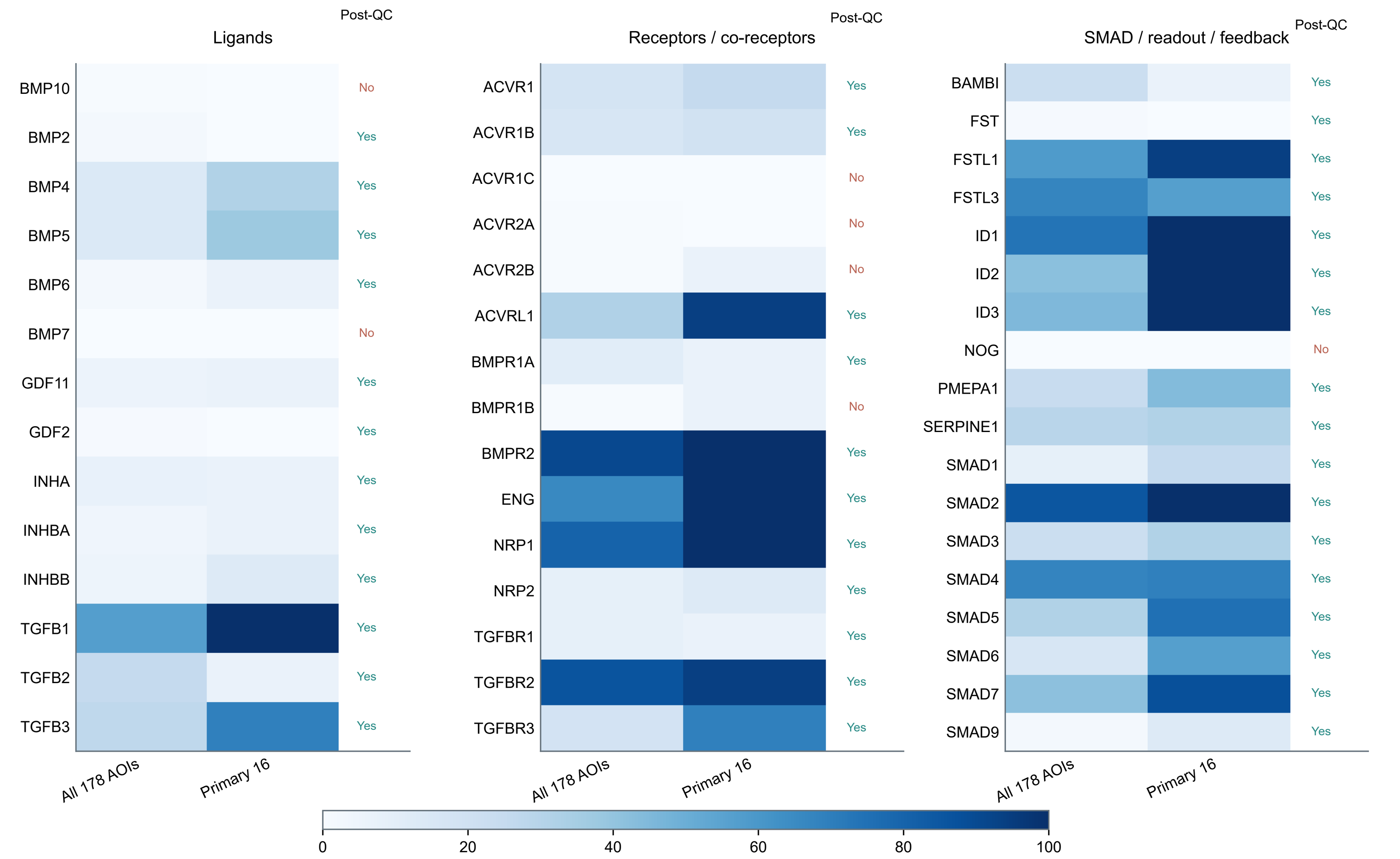


Post-QC Yes: available in the authors' 12,800-gene matrix; No: unavailable and excluded from expression testing.
Coverage is assay detectability, not differential expression; all 47 predefined family components remain in the source CSV.
The focused main-figure heatmap is not repeated here.

#### G Strict-LV, pre-LVAD and control sensitivities

All 16 detection-eligible pathway genes across five prespecified donor-level cohorts

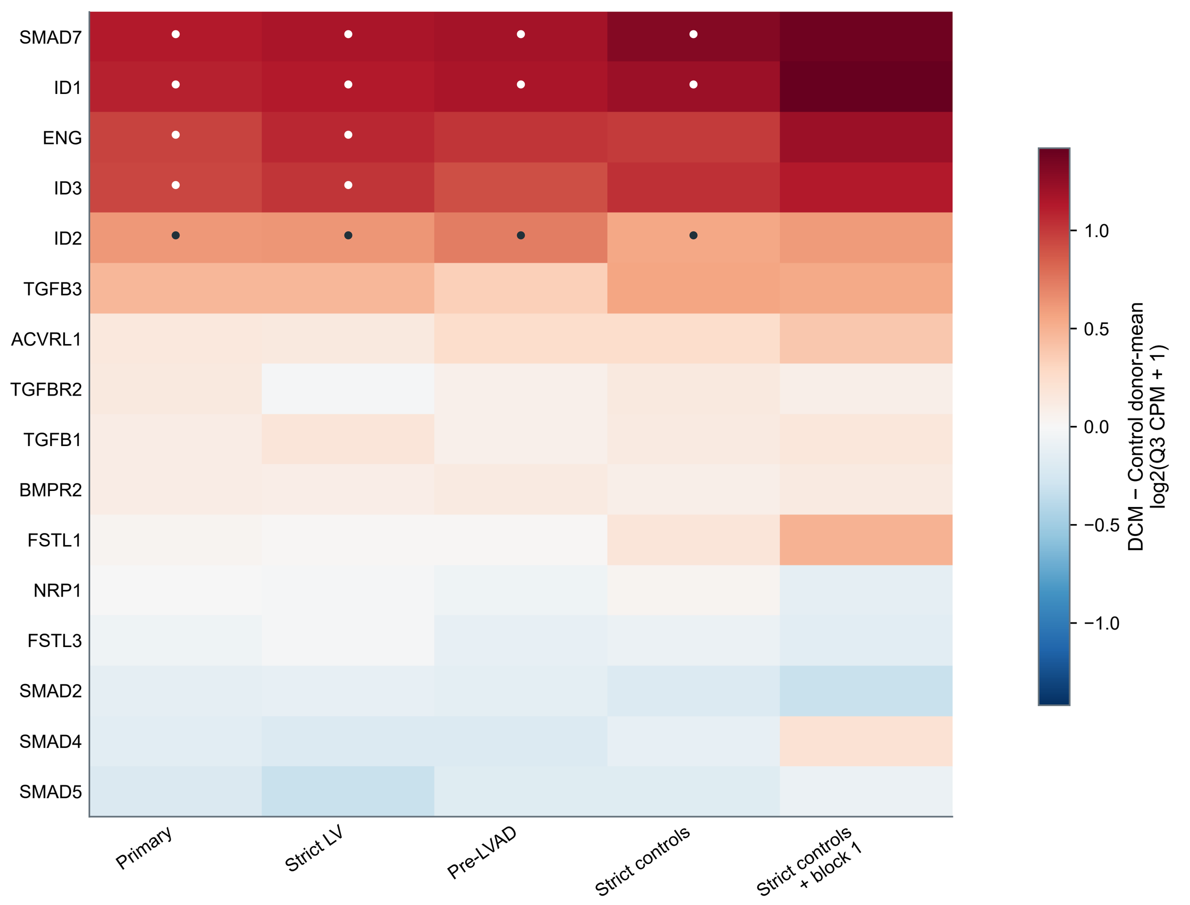


Dot: BH FDR <0.05 within that 16-gene cohort analysis. Biological replicate is donor; repeated AOIs are averaged.
Cohorts: primary 4/7 Control/DCM donors; strict LV 4/6; pre-LVAD 4/6; strict controls 3/7; block-1 strict controls 3/4.
The final cohort has limited permutation resolution. This sensitivity panel does not replace the prespecified primary analysis.

#### H Robustness of selected pathway components

Direction and BH-FDR support in the four major cohorts used for the main-figure robustness label

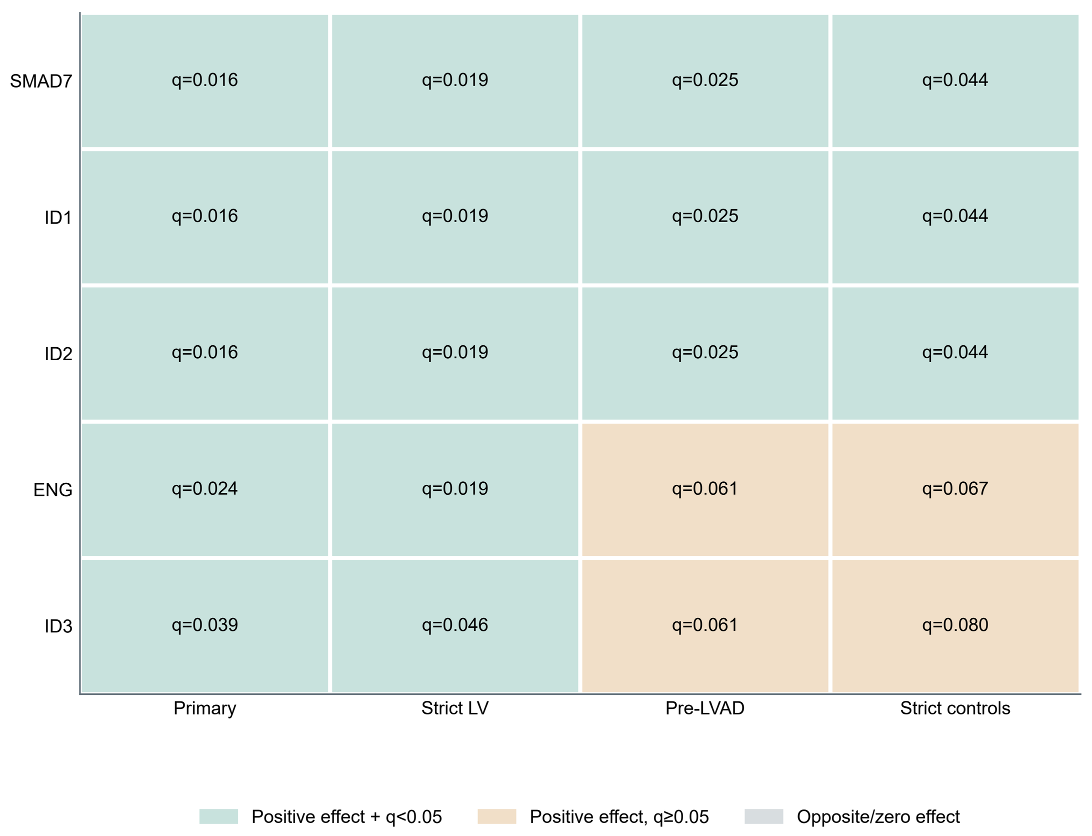


All five selected components have positive DCM−Control effects across major cohorts; FDR support varies by component/set.
"Robust across major sensitivities" requires q<0.05 in primary, strict LV, pre-LVAD and strict-control analyses.
This is the detailed source for that label, not a new composite score or separate validation experiment.

#### I Why ACVR2A could not be evaluated

ACVR2A was assayed, but raw detection was below the authors' expression-QC requirement

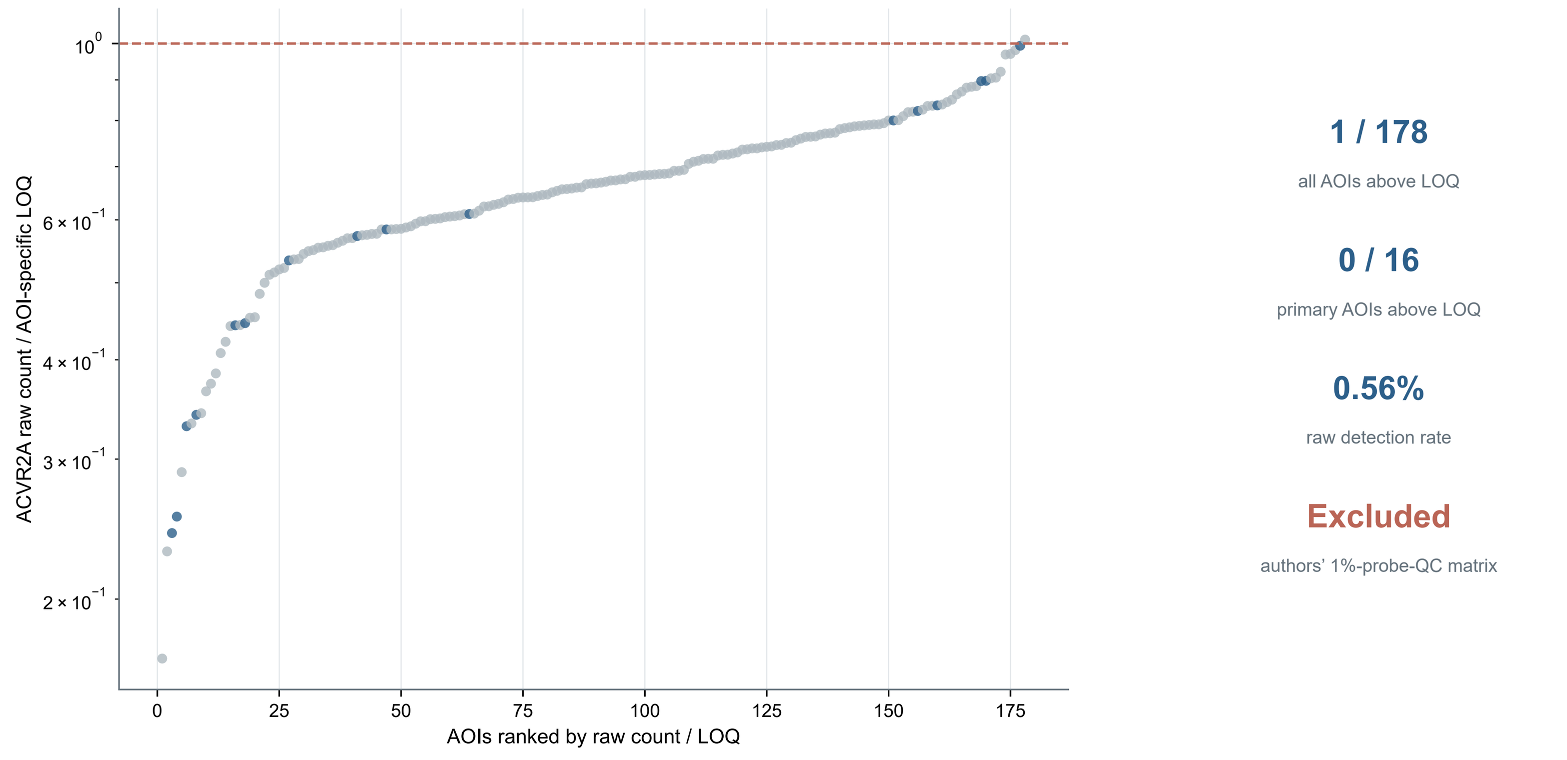


Dashed line: raw count equals the AOI-specific LOQ. Blue: primary 16 LV vessel AOIs; gray: all other AOIs.
ACVR2A had one endogenous probe and was present in the raw panel, but it was absent from the authors' post-QC matrix.
Raw counts are used only to document assay failure. ACVR2A expression, group effects and receptor validation are not evaluated.
